# Pangenomes reveal genomic signatures of microbial adaptation to experimental soil warming

**DOI:** 10.1101/2023.03.16.532972

**Authors:** Mallory J. Choudoir, Achala Narayanan, Damayanti Rodriguez-Ramos, Rachel Simoes, Alon Efroni, Abigail Sondrini, Kristen M. DeAngelis

## Abstract

Below-ground carbon transformations represent a natural climate change mitigation solution, but newly-acquired traits adaptive to climate stress may alter microbial climate feedback mechanisms. To better define microbial evolutionary responses to long-term climate warming, we study microorganisms from an ongoing *in situ* soil warming experiment at the Harvard Forest Long-term Ecological Research (LTER) site where, for over three decades, soils are continuously heated 5 °C above ambient temperatures. We hypothesize that across generations of chronic warming, genomic signatures within diverse bacterial lineages reflect trait-based adaptations related to growth and carbon utilization. From our bacterial culture collection isolated from experimental heated and control plots, we sequenced genomes representing taxa dominant in soil communities and sensitive to warming, including lineages of Alphaproteobacteria, Actinobacteria, and Betaproteobacteria. We investigated differences in genomic attributes and patterns of functional gene content to identify genomic signatures of adaptation. Comparative pangenomics revealed accessory gene clusters related to central metabolism, competition, and carbon substrate degradation. Overall, genomes from control plots were relatively enriched in carbon and fatty acid metabolism pathways, while genomes from heated plots were relatively enriched in nitrogen metabolism pathways. We also observed differences in global codon usage bias between heated and control genomes, suggesting potential adaptive traits related to growth or growth efficiency. This effect was more varied for organisms with fewer 16S rrn operons, suggesting that these organisms experience different selective pressures on growth efficiency. Together, these data illustrate the emergence of lineage-specific traits as well as common ecological-evolutionary microbial responses to climate change.

## 0.1 Importance

Anthropogenic climate change threatens soil ecosystem health in part by altering below-ground carbon cycling carried out by microbes. Microbial evolutionary responses are often overshadowed by community-level ecological responses, but adaptive responses represent irreversible changes in traits and functional potential. We predict that microbes are adapting to climate change stressors, like soil warming. To test this, we analyzed the genomes of bacteria from a soil warming experiment where soil plots have been experimentally heated 5 °C above ambient for over 30 years. While genomic attributes were unchanged by long-term warming, warming altered functional gene content related to carbon and nitrogen usage and genomic indicators of growth efficiency. These responses may represent new parameters in how soil ecosystems feed back to the climate system.

## 1 Introduction

Humanity faces an existential threat with accelerating soil loss due to habitat degradation and climate change Handelsman and Cohen (2021), but the irreversible consequences of microbial adaptation to climate change remain insufficiently explored. Microbes are both impacted by climate change and are themselves agents of global change Cavicchioli et al (2019); Gillings and Paulsen (2014); Tiedje et al (2022). Soils are home to the most diverse microbial communities on the planet, and soil health is predicated on microbial diversity and activity Wagg et al (2014); Delgado-Baquerizo et al (2017). Ultimately, how soil microbes respond to and evolve with changing soil habitats will determine terrestrial ecosystem function under future climate change scenarios.

Rising soil temperatures due to climate change are altering microbial global nutrient cycles in complex ways Butler et al (2012); Jansson and Hofmockel (2020). Worse, chronic warming may disrupt the microbial mechanisms governing carbon fluxes between the biosphere and atmosphere, mechanisms which may provide a natural attenuation of the self-reinforcing feedbacks to climate Griscom et al (2017). Soils release considerable amounts of atmospheric carbon dioxide (*CO*_2_) through microbial decomposition of organic matter Schlesinger and Andrews (2000) but are also the largest terrestrial reservoir of global carbon Bahiri-Elitzur and Tuller (2021); Oelkers and Cole (2008). Microbes mediate key roles in carbon fluxes between terrestrial and atmospheric systems, evidenced by the fact that incorporating microbial processes into Earth system models improves global carbon pool projections Wieder et al (2013, 2014, 2015). Temperate forests actively sequester 0.7 Pg C year-1, but the potential for forests to mitigate climate change is uncertain Dixon et al (1994); Lal (2005); Pan et al (2011). The fate of terrestrial forest carbon pools largely depends on how microbes respond to global change factors.

Long-term field warming experiments allow us to make insights into what soils could look like in a warming world. For over three decades, temperate forest soils at the Harvard Forest Long-term Ecological Research (LTER) site are continuously heated *in situ* 5 °C above ambient temperatures Peterjohn et al (1994). After 26 years, warming degraded the soil organic carbon quantity and quality, increased net *CO*_2_ emissions, and altered microbial activity in heated versus control plots Pold et al (2017); Melillo et al (2017). Though fungal biomass and community structure were drastically changed with warming Frey et al (2008); Pec et al (2021), bacterial biomass and community structure remained relatively unperturbed by warming, with significant changes observed only after 20 years in the organic horizon DeAngelis et al (2015). Finally, bacteria isolated from the heated Harvard Forest plots showed greater potential to degrade lignin and other plant polymers compared to those isolated from control plots Pold et al (2015, 2016), suggesting that traits related to substrate use may explain long-term dynamics of climate stress on microbial systems.

Both ecological and evolutionary mechanisms shape how environmental microbes respond to climate change. The metacommunity concept Leibold et al (2004) describes how regional pools of microbial diversity, high dispersal rates, and a propensity for dormancy all determine how microbial communities change, or not, in response to environmental disturbance Shade et al (2012); Allison and Martiny (2008). In general, ecological filtering strongly shapes microbial community assembly Nemergut et al (2013); Freedman and Zak (2015). Canonically we expect to observe similar ecological forces shaping microbial community rapid responses to soil warming. But due to large population sizes and short generation times, co-occurring ecological and evolutionary forces can act on more similar time scales for natural microbial populations across years or decades Koskella and Vos (2015). Evolution introduces heritable traits into microbial populations, and we don’t know the cascading impacts of local or regional adaptive traits on global soil ecosystem functions. Theoretical carbon models need to account for ecological and evolutionary feedbacks across spatiotemporal scales within these dynamic selective landscapes Abs et al (2022).

Based on our previous works, ecological filtering in heated plots likely selects for microbes able to degrade more recalcitrant carbon substrates Pold et al (2015, 2016) in response to warming-induced decreases in carbon quality and quantity Pold et al (2017). Evolutionary processes may also contribute to differences in carbon metabolism between treatment plots. Long-term warming causes thermal acclimation of soil respiration Bradford et al (2008, 2019), though this effect may be seasonal Domeignoz-Horta et al (2022). Microbial thermal adaption likely involves physiological trade-offs altering growth rate and efficiency Bradford (2013). Life history trade-offs between growth efficiency, resource acquisition, and stress tolerance shape organismal fitness Malik et al (2020) linking population and community-level trait evolution.

We hypothesize that Harvard Forest microbial lineages have evolved adaptive traits related to carbon substrate metabolism and growth efficiency across generations of chronic soil warming. To maximize our ability to identify genomic signatures of adaption, we used comparative pangenome approaches on collections of closely related bacterial taxa across multiple independent lineages spanning Actinobacteria, Alphaproteobacteria, and Betaproteobacteria. We explored genomic attributes, patterns of functional gene content presence and absence, and codon usage between heated and control genomes to define adaptive traits which have emerged in this system since the onset of the long-term warming experiment.

## 2 Results and Discussion

### 2.1 Harvard Forest soil warming experiment bacterial genome collection

We conducted comparative pangenomic analyses on bacteria isolated from a long-term warming field experiment to test our hypothesis that microbes adapt to local environmental change stressors. All bacterial isolates originated from a multi-decade climate manipulation experiment Melillo et al (2017) at the Harvard Forest Long-Term Ecological Research (LTER) site (Petersham, MA, USA). The Prospect Hill warming experiment was established in 1991 in a mixed deciduous forest stand. Heated cables buried 10 cm below the surface continuously maintain soil temperatures in heated plots at 5 °C above the ambient temperatures of control plots (6 replicate plots per treatment of 6 x 6 m) Peterjohn et al (1994). Between 2013-2020, we built a diverse culture collection of bacteria isolated from heated and control plots (see Methods and Table S1 for isolation details) targeting taxa dominant in soil communities and sensitive to long-term warming DeAngelis et al (2015).

Isolates for whole genome sequencing were selected from our culture collection based on their full-length 16S ribosomal RNA gene phylogeny and taxonomy. Our goal was to build collections of closely related genomes representing multiple independent lineages of dominant soil taxa with replicates from heated and control plots. The final data set contained 91 genomes belonging to 5 clades spanning three phyla, including Actinobacteria *Kitasatospora* spp. (n=28), Alphaproteobacteria *Bradyrhizobium* spp. (n=15), Alphaproteobacteria *Rhizobium* spp. (n=14), Betaproteobacteria *Paraburkholderia* spp. (n=16), and Betaproteobacteria *Ralstonia* spp. (n=18) (Figure 1, Table S1). We choose these clades because we previously observed increased abundances of Actinobacteria and Alphaproteobacteria in Harvard Forest heated plots compared to controls DeAngelis et al (2015); Pold et al (2015). We also included Betaproteobacteria here as a contrasting example of abundant soil taxa less responsive to warming. Partial 16S rRNA gene sequences from all *Bradyrhizobium* and some *Paraburkholderia* genomes mapped to the most abundant OTUs from our previous study DeAngelis et al (2015). While these particular Alphaproteobacteria and Betaproteobacteria OTUs had higher relative abundances in heated and control plots, respectively, none of these were indicator taxa of warming treatment (Figure S1).

**Fig. 1.**
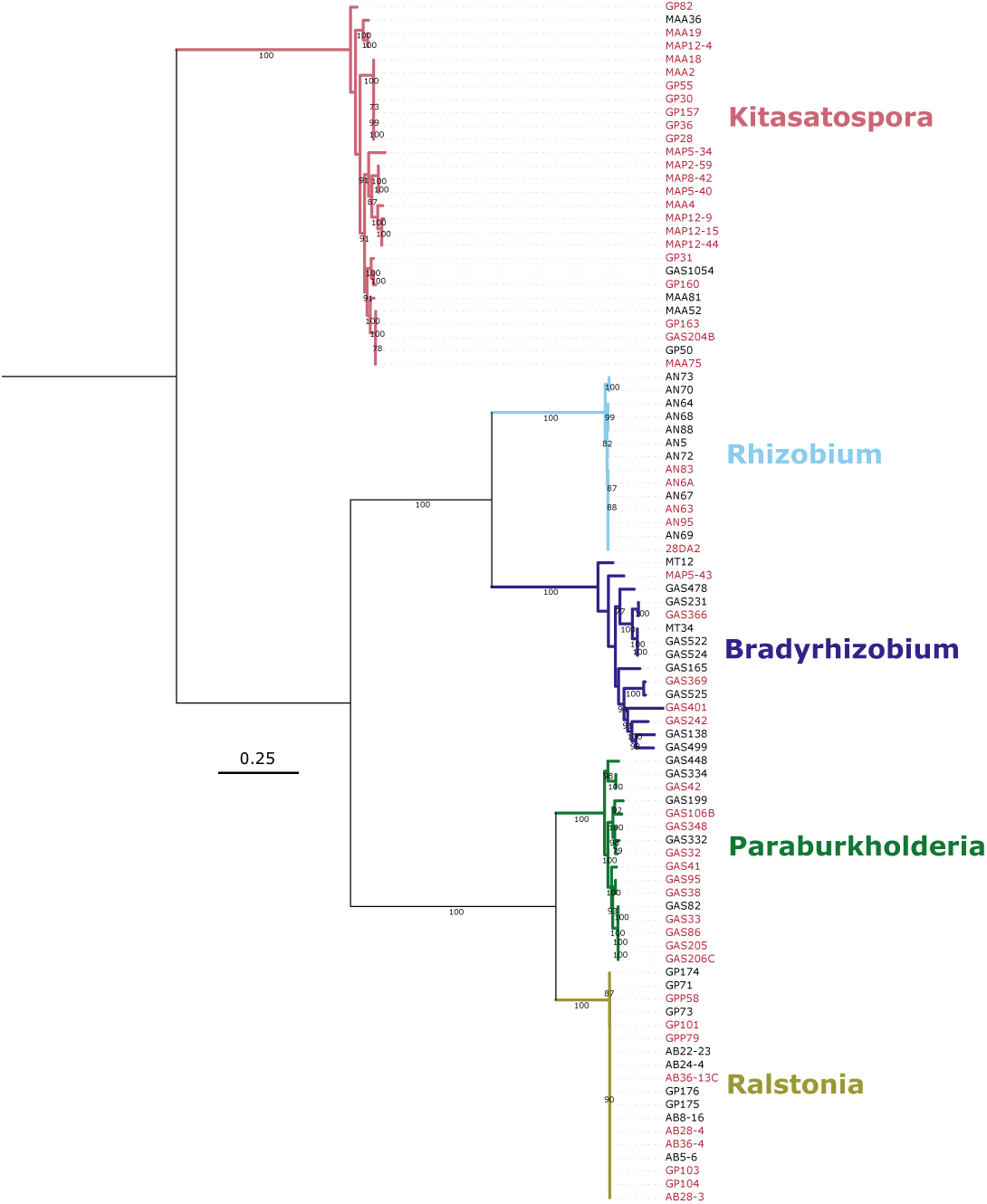
Ribosomal protein gene phylogeny. Tree shows the phylogenetic relationships between all 91 isolates (Table S1) constructed from 39 concatenated ribosomal protein gene nucleotide sequences derived from anvi’o HMM searches. Genome names are colored according to treatment group, with heated in red and control in black. Phylogenetic clades are also labeled and colored according to their pangenome group. Scale bar indicates nucleotide substitutions per site. Bootstrap values greater than 70 from 100 iterations are noted on tree branches. Tree is mid-rooted.

Actinobacteria, Alphaproteobacteria, and Betaproteobacteria are all globally dominant soil phyla Janssen (2006); Delgado-Baquerizo et al (2018) and represent broad functional guilds delineated by varying life history strategies Schimel and Schaeffer (2012); Fierer et al (2007). Though many ecosystem carbon models generally consider all bacteria as copiotrophic, operationally there exists a spectrum of growth efficiencies among phyla. Soil-dwelling Betaproteobacteria and Alphaproteobacteria are generally considered more copiotrophic with rapid growth and strong responses to labile carbon, while Actinobacteria typically embody more oligotrophic traits including slower growth and catabolic enzyme production Fierer et al (2007); Trivedi et al (2013). Actinobacterial genomes also harbor abundant cellulases that breakdown plant-derived biomass in soils Lewin et al (2016); Berlemont and Martiny (2013). This aligns with a temperature sensitivity lab incubation study by Oliverio et al (2017) who found that across all sites, nearly all Actinobacteria taxa and approximately half of all Alphaproteobacteria taxa responded positively to warming.

The objective of our study was to identify genomic signatures that may link functional potential to trait-based adaptation. We generated high quality draft genomes according to community standards Bowers et al (2017) using a combination of sequencing platforms (Table S2 and Table S3). Total contigs in assembled genomes varied considerably between sequencing platforms (Table S2). For *Paraburkholderia* spp., the number of assembled contigs strongly correlated with average gene length (Spearman’s rho = -0.85, P-value *<* 0.0001) and genes per kb (Spearman’s rho = 0.92, P-value *<* 0.0001). We also observed that assemblies with approximately *>* 20 contigs had markedly fewer 16S rRNA gene copies than expected based on clade membership, so we omitted these genomes from downstream 16S rRNA gene analyses (see Table S2). However, there was no relationship between the number of assembled contigs and total genes or total gene clusters. We also assessed draft assembly quality by estimating completion based on the presence of bacterial single-copy genes, and all genomes have high estimated completion *>* 94% (Table S2-S3). Given our focus on functional gene content, we concluded that these draft genomes were adequate for our study.

### 2.2 Phylogeny and genomic attributes

Phylogenetic reconstruction of concatenated single-copy ribosomal protein genes showed five monophyletic clades with varying levels of intra-group relatedness (Figure 1). All clades shared greater than 97% similarity at 16S rRNA gene sequences (Table 1, Figure S1) suggesting that pangenomes represent intra-species comparisons. However, genome wide average nucleotide identity (ANI) ranged from 78-99% (Table 1) indicating that some groups represent more distant genus-level comparisons Barco et al (2020) while other groups span inter-species to intra-species comparisons (ANI *≥* 95%) Konstantinidis and Tiedje (2005) to nearly identical isolates (ANI *>* 99%). Our goal here is not to demarcate species or taxonomic boundaries, only to frame the phylogenetic and genomic relatedness of our genomes within the ongoing discussion of bacterial diversity.

**Table 1.**
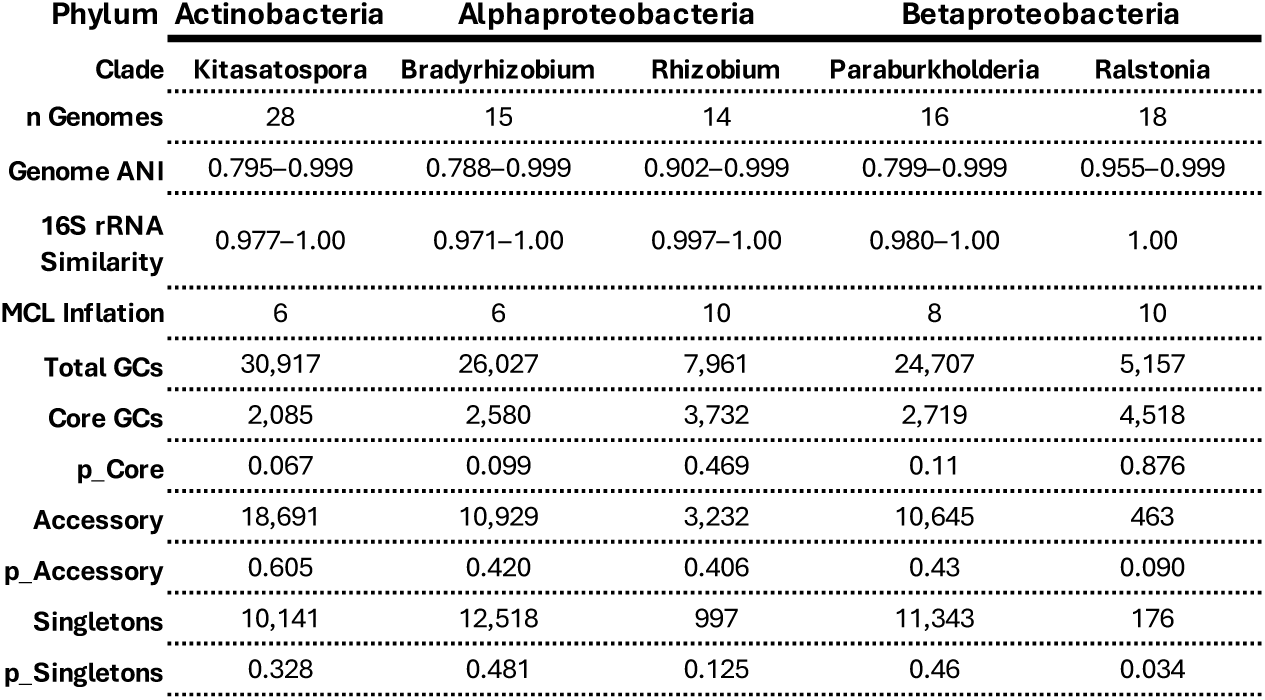
Pangenome characteristics for each clade. Columns list pangenome phylum, clade name, number of genomes (n) in each pangenome, genome wide average nucleotide identity (ANI), 16S rRNA gene similarity, Markov Cluster Algorithm (MCL) inflation values used for generating gene clusters, total number of gene clusters (GCs), number of core GCs, portion of GCs that belong to the core gene pool (pCore), number of accessory GCs, portion of GCs that belong to the accessory gene pool (pAccessory), number of singletons GCs, and portion of GCs that are singletons (pSingletons).

To identify adaptive genomic traits associated with long-term soil warming that are shared among the clades, we first determined if broad genome attributes varied by warming treatment. While all genome traits differed significantly between clades, they did not vary by treatment within any of the clades (Kruskal-Wallis rank sum test on the interaction between clade and treatment followed by post-hoc pairwise Dunn test across intra-clade treatment groups, P-value *>* 0.1). Traits tested included genome size, total number of genes, genes per kb, average gene length, G+C content, 16S rRNA gene copy number, total number of pangenome gene clusters, number of unique or singleton genes, and the total number of genes annotated as carbohydrate-active enzymes (CAZymes) (Table S2). When controlling for phylogeny, these attributes also did not vary by treatment (phylogenetic ANOVA, P-value *>* 0.1).

### 2.3 Pangenome structure, gene content, and functional enrichment between warming treatments

We used comparative pangenomics to determine if genomes from heated treatments harbored different patterns of functional gene content compared to control treatments and whether these differences were consistent between clades. Pangenome structure and composition varied across phyla and between clades, ranging from more open to more closed pangenomes (Figure 2, Table 1). For instance, the *Ralstonia* pangenome contained relatively few gene clusters (GCs) (n=5,175 GCs) with nearly all of these belonging to the conserved core genome (n=4,518 GCs) and very few strain-specific or singleton GCs. Conversely, the other Betaproteobacteria clade *Paraburkholderia*, showed considerably higher gene richness with approximately five times more gene clusters (n=24,707) with only 11% of GCs (n=2,719) belonging to the core gene pool, and many singletons. We observed a contrasting pattern between the more open pangenome of Alphaproteobacteria *Bradyrhizobium* and the more closed pangenome of Alphaproteobacteria *Rhizobium*. We also observed a high portion of accessory and singletons gene clusters in the *Kitasatospora* pangenome, which is unsurprising as Actinobacteria are well known for their genomic and ecological diversity Lewin et al (2016).

**Fig. 2.**
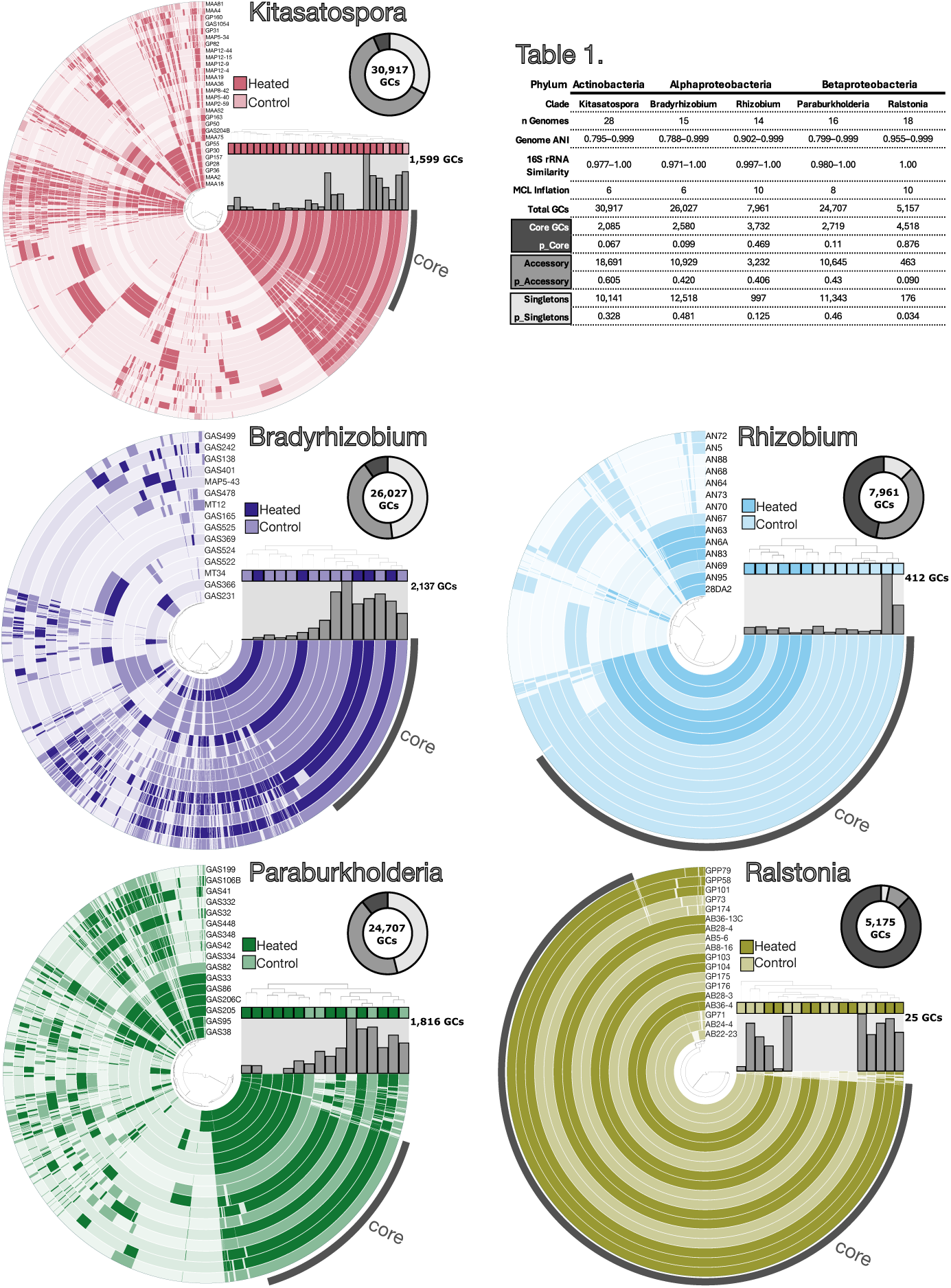
Pangenome composition and structure across clades. For each pangenome plot, concentric circles show gene cluster presence/absence across all genomes. The internal dendrogram orders genomes by gene cluster frequencies. For visualization singletons were removed before plotting, and core gene clusters are labeled on each pangenome. Pangenomes are colored by clade, and genomes are colored by warming experiment treatment as shown in the legend. Bar graphs show the number of singleton gene clusters present in each genome. Finally, donut plots illustrate the portion of GCs belonging to the core, accessory, and singleton gene pools for each pangenome. See Table 1 for exact values.

Patterns of gene variation across populations evolve through various mechanisms that depend on both population parameters and ecological interactions McInerney et al (2017); Domingo-Sananes and McInerney (2021), and often these mechanisms are impossible to disentangle Baumdicker and Kupczok (2023). Pangenomes are constantly in flux and are themselves shaped by environmental dynamics and ecological-evolutionary feedbacks Cummins et al (2022). For example, lineages with large accessory gene pools have a deeper reserve of genes that may be shared within a population during times of stress or environmental change, facilitating ecological adaptation Conrad et al (2022). However, a recent pangenome study found that environmental stressors including acidity, heat, drought, and salt stress all led to reductions in gene richness of *Bradyrhizobium diazoefficiens* Simonsen (2022), with accessory genes more prone to habitat-specific loss. These dynamics may reflect a transition between niche generalists possessing more open pangenomes and niche specialists more closed pangenomes Brockhurst et al (2019).

We did not observe gene clusters strongly associating with heated or control treatments for any of the clades (Figure 2). In other words, a discrete “heated core” or “control core” does not exist in our pangenome collection. Because we didn’t see a strong treatment-specific signal for individual gene clusters, we also investigated differences in functional gene content between treatment to discover annotations that were over- or underrepresented in genomes derived from chronically heated soils. Genomes were annotated hierarchically: gene cluster annotations varied by functional orthologs (n=334 KOfams, Table S4) or by functional units of genes within metabolic pathways (n=35 KEGG pathway modules, Table S5). Finally, the estimated metabolic capabilities of strains also varied between treatments (n=15 metabolic pathways, Table S6).

Gene clusters with functional orthologs related to carbon and nitrogen metabolism, gene regulation, and heavy metal metabolism were differently associated with heated and control genomes (Table S4). Overall, genomes from control plots were relatively enriched in functional genes related to carbon and fatty acid metabolism, while genomes from heated plots were relatively enriched in functional genes related to nitrogen metabolism (Table S4). Although these patterns differed across individual clades. For control genomes, this included relative enrichment of fatty acid, methane, and acetyl-CoA pathways for *Kitasatospora*; amino acid metabolism (polyamine biosynthesis) for *Bradyrhizobium*; purine degradation, nitrate assimilation, and amino acid degradation for *Rhizobium*; and central carbohydrate metabolism for *Paraburkholderia* (Table S5-S6). Conversely, genomes from heated plots were relatively enriched for metabolic pathways involved in central carbohydrate metabolism and amino acid (cysteine) biosynthesis for *Kitasatospora*; central metabolism and the GABA (gamma-aminobutyrate) shunt for *Bradyrhizobium*; and methionine and organic compound (phthalate) degradation for *Rhizobium* (Table S5-S6). *Ralstonia* genomes lacked functional genes and metabolic pathways differently associated with warming treatment. Since it’s unlikely that individual gene gain or loss events will lead to differences in metabolic functional potential between genomes, these pathway differences between heated and control groups likely reflect either longer timescales or ecological and evolutionary feedbacks mechanisms shaping genome evolution.

Pathways associated with natural product biosynthesis and competition varied across warming treatment for the Proteobacteria clades. *Rhizobium* and *Bradyrhizobium* genomes from heated plots were relatively enriched in pathways related to beta-lactam biosynthesis and resistance, respectively, while *Paraburkholderia* genomes from control plots were enriched in the vancomycin resistance pathway (Table S5-S6). For *Kitasatospora*, fosfomycin and betacyanin biosynthetic pathways were enriched in genomes from control plots, while calicheamicin and dihydrokalafungin biosynthetic pathways were enriched in genomes from heated plots (Table S6). Differences in secondary metabolite repertoires could indicate different biotic competition pressures due to warming stress, or an indirect effect of the severe soil organic carbon depletion in the chronically heated soils. Over three decades, heated soils have lost 34% of soil organic matter Melillo et al (2017), accompanied by greater depletion of carbohydrates and other relatively more accessible forms of heterotrophic substrates Pold et al (2017).

### 2.4 Carbohydrate-active enzymes

Given that differences in available carbon is the major environmental determinate driving ecological differentiation between heated and control soils, next we identified genes functionally annotated as carbohydrate-active enzymes Drula et al (2022), or CAZymes. These enzymes catalyze the breakdown, degradation, or modification of carbohydrates and glycoconjugates. The total number of CAZyme annotations did not differ across warming treatment for all clades (Table S2). Clade membership, but not warming treatment or the interaction between treatment and clade, was a strong predictor for CAZyme composition (PERMANOVA, P-value=0.001, R2=0.85) (Figure S2). Within clades, treatment is a significant predictor of CAZyme composition for *Rhizobium* (PERMANOVA, P-value=0.019, R2=0.27), but not for the other clades.

While warming treatment did not explain variation in the total number or composition of CAZyme annotations across genomes, we observed individual CAZyme families and sub-families with different associations across heated and control plots (Table S7). For example, glycosyl-transferase (GT) enzymes were depleted in heated *Paraburkholderia* genomes. Heated genomes of *Rhizobia* and *Kitasatospora* were relatively enriched in polysaccaride lyases enzymes involved in the breakdown of alginate (Table S7). Conversely, glycoside hydrolase (GH) enzymes with complex substrates such as xylan, chitin, and glycogen were relatively depleted in heated *Kitasatospora* genomes. Chronic soil warming depleted total soil organic matter and altered carbon composition, resulting in lower quantity as well as quality of carbon substrates in the heated versus control plots Pold et al (2017). Further, a previous study found that a greater portion of bacterial strains isolated from heated plots utilized complex carbon substrates (including xylan and cellulose) compared to strains isolated from control plots Pold et al (2016). Together, these data suggest that microbial substrate availability and carbon utilization shape microbial adaptation to soil warming.

### 2.5 Codon usage bias

Codon usage bias (CUB) is an estimate of the degree to which an organism demonstrates non-random, or preferential, use of synonymous codons in genes. Codon bias varies with microbial physiology through mechanisms involving protein translation accuracy and/or efficiency Sharp et al (2005, 2010); Bennetzen and Hall (1982). Different measurements of codon usage bias offer various insights into codon usage across genes and gene expression levels across genomes Bahiri-Elitzur and Tuller (2021). We measured CUB as the effective number of codons (ENC) Wright (1990) and as the Measurement Independent of Length and Composition (MILC) Supek and Vlahoviček (2005). CUB varied between heated and control genomes for both metrics.

ENC calculates the frequencies of different codons in a gene sequence, and ranges from 20 (extreme bias, only a single codon is used for each amino acid) to 61 (no bias, all synonymous codons are equally used). We calculated CUB distributions between heated and control strains and across core and accessory genes, which may reflect differing selection pressures across gene pools or differences in horizontal gene exchange dynamics Moszer et al (1999). Genome wide gene-level ENC distributions differed between heated and control genomes for all clades (Wilcox rank sum test, P-value *<* 0.0001) except *Paraburkholderia* spp. and (P-value = 0.57) *Ralstonia* spp. (P-value = 0.73) (Figure S3A). Core (Figure S4A) and accessory (Figure S5A) gene ENC distributions differed between heated and control genomes for all groups (P-value *<* 0.01) except *Ralstonia* (P-value *>* 0.6).

MILC calculates codon usage of a sequence as the distance against the expected distribution of a reference gene set. When determined against set of highly expressed genes, like ribosomal proteins, higher values indicate more dissimilar patterns of codon usage. Genome wide gene-level MILC distributions differed between heated and control genomes for all clades (Wilcox rank sum test, P-value *<* 0.001) except *Rhizobium* spp. (P-value = 0.18) (Figure S3B). Core gene MILC distributions differed between heated and control genomes for all clades (P-value *<* 0.03) besides *Bradyrhizobium* (P-value = 0.40) (Figure S4B). Accessory gene MILC distributions differed between heated and control genomes for all clades (P-value *<* 0.01) except *Bradyrhizobium* spp. and *Rhizobium* (P-value *>* 0.1) (Figure S5B).

While gene-level CUB metrics show genome wide distribution patterns, codon usage summarized for each genome provides a single metric to compare between treatment and gene pools. Strain-level median ENC codon usage differed between heated and control *Kitasatopora* genomes for “All” and “Accessory” gene pools (Wilcox rank sum test, P-value *≤* 0.05) (Figure 3A). Strain-level MILC codon usage nearly differed between heated and control *Paraburkholderia* genomes for core and accessory gene pools (P-value *≤* 0.1). Median ENC and MILC varied significantly across gene pools for all clades (ANOVA, P-value *<* 0.001) (Figure 3), but treatment also contributed to differences in median ENC across gene pools for *Kitasatopora* (P-value = 0.0067) and *Rhizobium* (P-value = 0.037) and median MILC across gene pools for *Paraburkholderia* (P-value = 0.0016) and *Ralstonia* (P-value = 0.033). In general, core genes tend to have codon bias patterns more similar to the ribosomal reference set relative to accessory genes, and these differences contribute to global codon usage patterns.

**Fig. 3.**
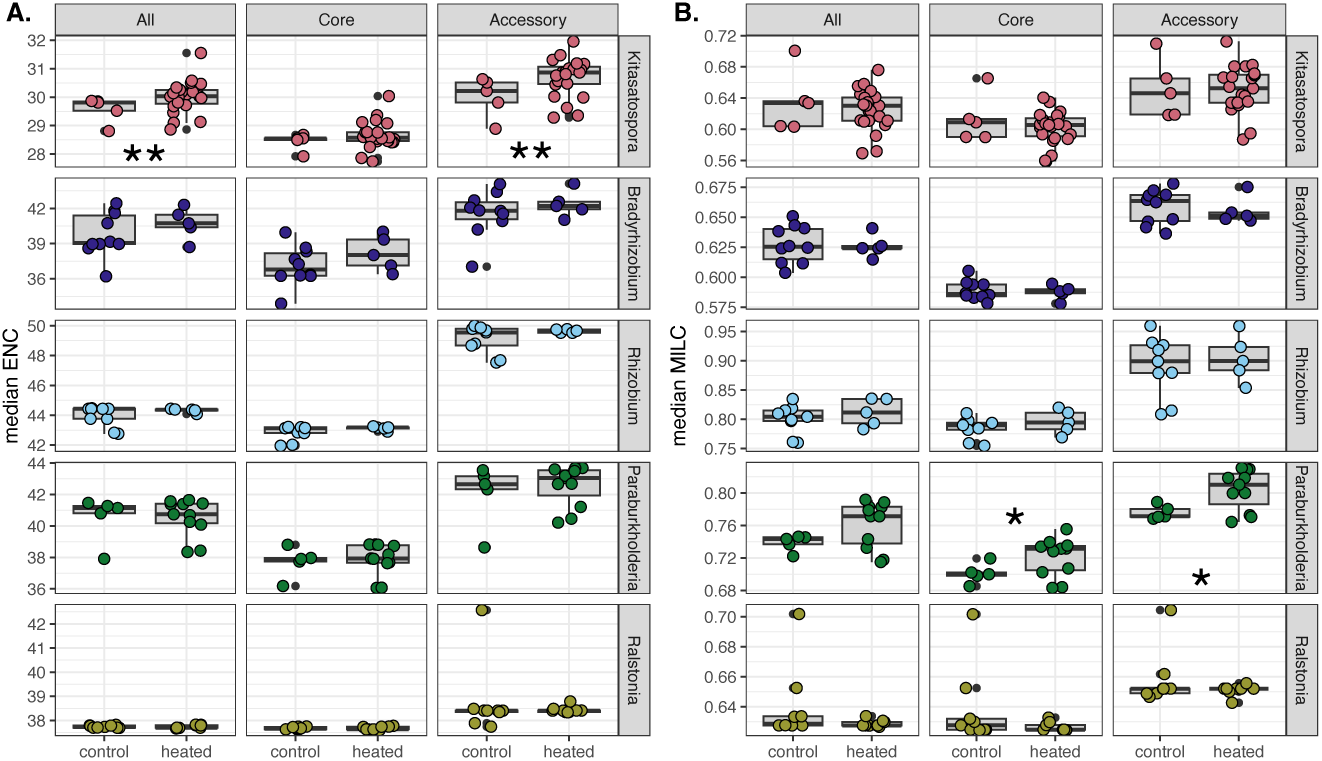
Strain-level median CUB values across gene pools. Plots illustrate the strain-level (i.e., summarized per genome) median codon usage between treatment groups across clades and across gene pools. Columns are codon usage summary for all genes (All), core genes (Core), or accessory genes (Accessory). Core genes belong to gene clusters that are present in all genomes within a pangenome clade. Accessory genes belong to gene clusters that are shared by at least two, but not all genomes within a pangenome clade. Note, singleton genes were not included in the accessory gene pool for this analysis. Boxplots show the interquartile range of codon usage values for control and heated genomes, and the line is the median, whiskers are 1.5 interquartile range, and black points are outliers. Circles show median codon usage values for each strain and are colored by clade. Figure 4A shows ENC (Wright 1990) values, and Figure 4B shows MILC (Supek et al. 2005) values. We calculated MILC values against a set of conserved ribosomal protein genes. Asterisks indicate Wilcox rank sum test P-values for differences in median codon usage between treatment *<*= 0.05 (*); *<* 0.10 *>* 0.05 (**).

Long-term warming was associated with less ENC codon bias (higher values) for Actinobacteria and Alphaprotebacteria clades but not Betaproteobacteria clades (Figure 3a). High MILC values, or codon usage less similar to highly expressed ribosomal protein coding genes, were observed in *Paraburkholderia* heated genomes compared to control genomes (Figure 3b). Genomes with different MILC distributions likely differ in their protein expression profiles, which may relate to growth rate and efficiency. This may indicate that genomes from chronically heated soils have lower global expression levels (i.e., related to maximum growth rate), or possess a smaller pool of genes with relatively less efficient or slower expression patterns.

Genomes with more 16S rRNA gene copies have stronger codon usage bias (ENC). Small subunit ribosomal (SSU) rRNA copy number reflects microbial growth traits and ecological strategies Klappenbach et al (2000). We observed a significant negative relationship between median ENC codon bias and 16S rRNA gene copy number for heated (Spearman’s rho = -0.67, P-value *<* 0.0001) but not control genomes (rho = -0.079, P-value = 0.53). This relationship differs between treatment (slope of linear model differs between groups, P-value = 0.048) (Figure 4). While SSU copy number does not vary between heated and control genomes, there may be other codon usage or ribosomal operon traits that result in growth efficiency adaptations related to resource availability Roller et al (2016) in warming soils.

**Fig. 4.**
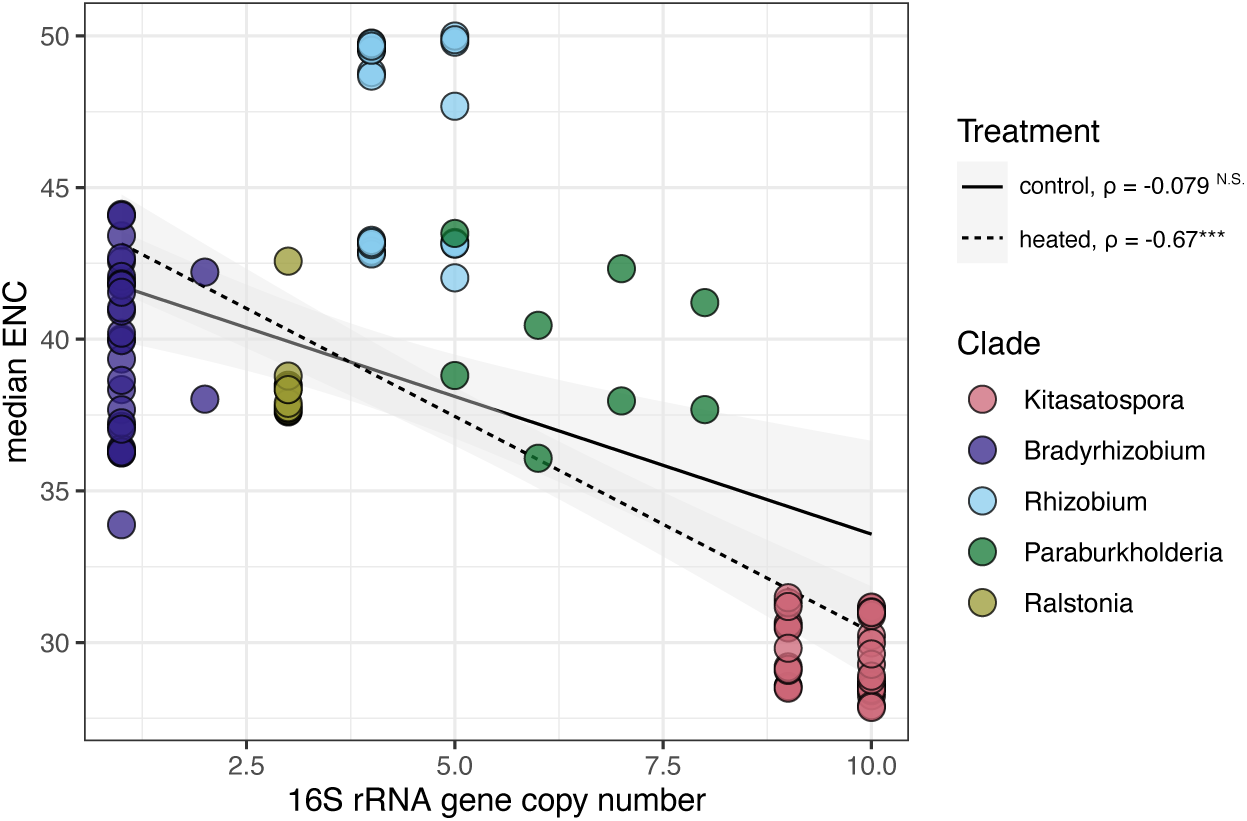
CUB versus 16S rRNA gene copy number. Plot illustrates the relationships between global codon usage bias (y-axis) and 16S rRNA gene copy number (x-axis) (Table S2). We omitted genome assemblies with over 20 contigs (see Harvard Forest soil warming experiment bacterial genome collection section). Circles show median values of strain-level ENC codon usage for each genome across all gene pools and are colored according to pangenome clade. Linear regression is plotted for heated (dashed line) and control (solid line) treatment. Shaded areas depict 95% confidence intervals. Spearman’s rho is reported, and asterisks note P-value *<* 0.0001.

Growing evidence suggests environmental drivers of microbial community-level codon usage. For instance, a recent cross-biome metagenomic study demonstrated direct impact of microbial habitat on codon bias and amino acid usage after controlling for other non-environmental variables Panda and Tuller (2023). Ultimately, a number of factors influence codon usage patterns in microbial genomes including genome wide G+C content Chen et al (2004) as well as tRNA abundances Ikemura (1985); Dong et al (1996). Global CUB correlates with microbial life history strategies, growth rate, and ecological adaptations Botzman and Margalit (2011); Roller et al (2013); Chen et al (2021), traits which appear to emerge in soil microbiomes exposed to long-term warming DeAngelis et al (2015); Liu et al (2021).

## 3 Conclusion

We examined genomes of bacteria exposed to three decades of climate warming to understand whether warmer soils select for common ecological traits, or if microbes also evolve and adapt to warmer soils. Warming caused a reduction in fungal but not bacterial biomass Frey et al (2008); DeAngelis et al (2015), suggesting a seasonal shift towards a more bacterial-dominated ecosystem and local adaptation to enable this change. We identified genomic signatures related to growth efficiency (Figures 3-4) and resource acquisition (Tables S4-S7), which supports our previous observations that depleted soil organic matter in warming plots leads to more oligotrophic life history strategies DeAngelis et al (2015) and utilization of more complex carbon substrates Pold et al (2016).

Microbial populations tend to adapt to local environmental conditions Kraemer and Boynton (2017). However, evidence for ecological adaptation (or lack thereof) varies across environmental, spatial, and phylogenetic distances Connor et al (2010); Prosser et al (2007); Leducq et al (2014); Kraemer and Kassen (2015). The question here is, does local adaptation act along the spatial and temporal scales of the Harvard Forest warming experiment? Based on an average generation time of two weeks for soil microbial communities estimated using 18O-labeled water Domeignoz-Horta et al (2022), 30 years corresponds to a minimum of 500 generations in our Harvard Forest system.

Under laboratory setting, microbes adapt to increasing temperatures rapidly Bennett et al (1992); Tenaillon et al (2012). In a previous study, after 1500 generations of selective temperature stress, the fungus *Neurospora* adapted to increased temperature through life history trait trade-offs, including increased resource allocation to spores but decreased growth rate and higher respiration Romero-Olivares et al (2015). Insights into microbial responses to climate change across longer time scales that impact ecosystem-level functions are narrow due to limited access to long-term warming experiments. Even more rare are observations of local adaptation *in situ*. A small, but growing, body of research demonstrates ecological and evolutionary responses resulting in adaptation to environmental change on similar timescales Martiny et al (2023). For example, in a recent reciprocal transplant study across a natural temperature gradient, ecological and evolutionary feedbacks were detected on a timescale of 1.5 years Chase et al (2021). In this analysis, consistent mutations were detected in genes related to nutrient acquisition and stress responses.

The pangenomes of five clades (Figure 2) reflect organisms with different nutrient acquisition strategies, based on carbohydrate-activated enzymes, functional gene content, and codon usage bias. Our comparative pangenome data supports our previous works suggesting microbial trait trade-offs between growth efficiency and carbon substrate utilization Domeignoz-Horta et al (2020); Pold et al (2016). Functional genomic potential of genomes from heated plots suggests increased competition under a regime of more limited resources due to chronic warming. Ultimately, population-level adaptations to climate change may aggregate at the community-level resulting in changes in biogeochemical functions Wallenstein and Hall (2012). Future research on the physiology and growth traits of these organisms will further strengthen our ability to predict microbial function from genome adaptation, and to better predict ecosystem function in a changing world from microbial eco-evolutionary responses to climate change.

## 4 Methods

### 4.1 Experimental site and bacterial culture collection

We isolated bacteria from the Harvard Forest long-term soil warming experiment plots between 2013–2020 using a range of enrichment conditions (Pold et al (2016); Domeignoz-Horta et al (2020) and Table S1) targeting taxa of interest. Briefly, we sampled soils (0–10 cm) using a stainless steel soil corer, manually separated mineral and organic horizons, and sieved (2 mm). We plated soil dilutions onto solid media (Table S1) and incubated at room temperature for 2-8 weeks. Following the initial enrichment, colonies were picked and streaked to isolation on 10% tryptic soy agar (TSA) or Reasoner’s 2A (R2A), genotyped with full length 16S rRNA gene sequences (27F-AGAGTTTGATCMTGGCTCAG and 1492R-TACGGYTACCTTGTTACGACTT Weisburg et al (1991)), and cryopreserved in 20% glycerol at -80 °C.

### 4.2 Genomic DNA extraction and whole genome sequencing

We inoculated colonies into 10 mL 10% tryptic soy broth (TSB), grew them to stationary phase at 28 °C with shaking, and pelleted the cells by centrifugation. We extracted high molecular weight genomic DNA (gDNA) from cell pellets using either the DNeasy Blood & Tissue Kit (Qiagen, Germantown, MD, USA) or a modified CTAB-lysozyme extraction protocol Larsen et al (2007) (with phenol-chloroform instead of chloroform and extended centrifugation times). We confirmed average fragment size of 30-50 kb with gel electrophoresis and quantified DNA using a Qubit Fluorometer (Thermo Fisher Scientific, Waltham, MA, USA) to verify at least 1 µg of gDNA. We determined DNA quality using a NanoDrop Spectrophotometer (Thermo Fisher Scientific, Waltham, MA, USA) with absorbance ratios 260/280 within 1.8-2.0 and 260/230 within 2.0-2.3.

A subset of genomes were sequenced and assembled at the DOE Joint Genome Institute (JGI) (Berkeley, CA, USA) using either PacBio RS (Pacific Biosciences, Menlo Park, CA, USA) or NovaSeq S4 (Illumina, San Diego, CA, USA) (Table S1). A subset of betaproteobacterial genomes were sequenced at SeqCenter (Pittsburgh, PA, USA) using NextSeq 2000 (Illumina, San Diego, CA, USA). All remaining genomes were sequenced in house using ONT MinION (Oxford Nanopore Technologies, Oxford, UK).

We prepared ONT MinION sequencing libraries using the Ligation Sequencing Kit SQK-LSK-109 (ONT, Oxford, UK). We multiplexed 6-8 genomes per sequencing run using the Native Barcoding Expansion Kit EXP-NBD104 (ONT, Oxford, UK). We skipped the Covaris g-TUBE shearing step to target long fragment DNA. Starting with 1 µg of DNA, we prepped samples with the NEBNext® FFPE DNA Repair Mix and NEBNext® Ultra™ II End Repair/dA-Tailing kits (New England Biolabs, Ipswich, MA, USA) and performed DNA cleanup Ampure XP beads (Beckman Coulter, Pasadena, CA, USA). We pooled approximately 150 ng of each barcoded sample for a final library of 700-1000 ng, and adapters were ligated to the sample with Blunt/TA ligase (New England Biolabs, Ipswich, MA, USA). The long fragment buffer was used in an extended 10 minute incubation at 37 °C to enrich for high molecular weight DNA. We primed the flow cell using the Flow Cell Priming Kit (ONT, Oxford, UK), mixed, and loaded approximately 15 fmols of the pooled library, sequencing buffer, and loading beads. Sequencing runs completed after 72 hours, and we ran high accuracy base calling on the raw fast5 data with the ONT software Guppy v4.5.4 or v6.1.1 (Oxford Nanopore Technologies, 2021).

### 4.3 ONT MinION genome assembly

For all genomes sequenced using ONT MinION, we performed either *de novo* long-read assembly or hybrid short and long-read assembly. We used Filtlong (https://github.com/rrwick/Filtlong) to sub-sample high quality reads (–min length 1000 –min mean q 85) to approximately 40X estimated genome coverage and Flye v2.8.1 Kolmogorov et al (2019) for *de novo* assembly. We used Racon v1.4.2 Vaser et al (2017) to generate a consensus assembly and performed final polishing with medaka v1.2.1 (Oxford Nanopore Technologies, 2018). For the subset of betaproteobacterial genomes with both short- and long-read sequence data, we used unicycler Wick et al (2017) to perform hybrid assembly. Quast Gurevich et al (2013) and CheckM Parks et al (2015) assessed the genome assembly quality. We considered a genome assembly high-quality if it had less than 5% contamination and more than 90% estimated completeness (Table S3) Bowers et al (2017).

### 4.4 Genome annotation

We conducted genomic analyses within the anvi’o v8 software ecosystem Eren et al (2021). Genome assemblies were imported into anvi’o with the program anvi-gen-contigs-database, and open reading frames were identified using Prodigal v2.6.3 Hyatt et al (2010). The program anvi-run-kegg-kofams assigned KEGG Ortholog (KO) functions to protein coding genes Aramaki et al (2020), and the program anvi-estimate-metabolism reconstructed the metabolic capabilities of each isolate genome Kanehisa et al (2012). Finally, we predicted catalytic and carbohydrate-binding module domains of carbohydrate-active enzymes (CAZymes) using anvi-run-cazymes to run Hidden Markov Model (HMM) searches against dbCAN HMMdb v11 Zhang et al (2018); Zheng et al (2023).

### 4.5 Pangenome construction

We constructed pangenomes for each clade with the program anvi-pan-genome. Protein comparisons were computed with DIAMOND Buchfink et al (2015) with minbit heuristic = 0.5 (default value) to eliminate weak matches. Finally, the Markov Cluster Algorithm (MCL) Dongen and Abreu-Goodger (2012), with an inflation value of 6, 8, or 10 depending on the phylogenetic relationships of the clade (Table 1), identified shared gene clusters (GCs) or collections of similar genes. Core GCs are defined as GCs present in all genomes. Accessory GCs are those present in at least one but not all genomes. Finally, singletons are GCs unique to a single genome. Within each clade, the program anvi-compute-genome-similarity computed pairwise average nucleotide identity using the software program pyANI and ANIb method Pritchard et al (2016).

### 4.6 Phylogenetic analysis

Phylogenetic relationships were reconstructed from conserved genes. 16S ribosomal RNA (rRNA) genes and single-copy ribosomal protein genes were identified with HMM searches using the command anvi-run-hmms with anvi’o databases Ribosomal RNA 16S and Bacteria 71. Nucloetide sequences for 16S rRNA genes and 38 concatenated ribosomal protein genes were extracted with the program anvi-get-sequences-for-hmm-hits and aligned with MUSCLE Edgar (2004). Maximum likelihood (ML) trees were built using the generalized time reversible nucleotide substitution model with gamma distributed rate heterogeneity among sites (GTRGAMMA) in RAxML -NG v1.2.2 Kozlov et al (2019), and bootstrap support was determined from 100 iterations. We used iTol Letunic and Bork (2019) to visualize trees. Finally, we used the R package phytools to perform phylogenetic ANOVA with 1000 simulations Revell (2012). We used BLASTn Camacho et al (2009) to query isolate 16S rRNA gene sequences at ¿99% identity against partial 16S rRNA gene sequences (254 bp) of the dominant subset community (n=155 OTUs, rank abundance) from a previous amplicon study conducted at the Harvard Forest warming experiment DeAngelis et al (2015).

### 4.7 Genome trait analyses

We used program anvi-compute-functional-enrichment-in-pan and anvicompute-metabolic-enrichment to identify individual gene clusters, functional annotations, and full metabolic pathway modules differently enriched in heated and control treatment groups Shaiber et al (2020). Discussions of functional annotations were based on unadjusted P-value cutoffs, but we acknowledge that this may generate false positives. We used principal coordinate analysis (PCoA) of Bray–Curtis dissimilarities to visualize differences in CAZyme composition between clades and warming experiment treatments with the R package phyloseq McMurdie and Holmes (2013) and permutational ANOVA to asses statistical differences following 999 permutations with vegan Oksanen et al (2022).

We used the R package coRdon Elek et al (2019) to calculate codon bias. First, we filtered genes less than 80 codons before calculating codon usage metrics including the effective number of codons (ENC) Wright (1990) and the Measure Independent of Length and Composition (MILC) Supek and Vlahoviček (2005). For MILC measurements we calculated genome wide codon bias with respect to a subset of ribosomal proteins, which we speculated to be highly expressed genes. All other statistical analyses were performed using R R Core Team (2022) statistical software, including FSA Ogle et al (2022) for Dunn’s post-hoc test of multiple comparisons.

## Data Availability Statement

All genomic data used in this study are publicly available. Genome sequence data is archived on the SRA under Bio-Projects PRJNA944970, PRJNA944974, PRJNA944975, PRJNA944977, and PRJNA944978 with BioSample accession numbers also referenced in Table S1. All draft genome assemblies are also archived on the DOE JGI’s Integrated Microbial Genomes and Microbiomes (IMG/M) under study names “Sequencing of bacterial isolates from soil warming experiment in Harvard Forest, Massachusetts, USA” and “Using genomics to understand microbial adaptation to soil warming” with GOLD Study IDs Gs0121248 and Gs0156716. IMG Taxon/Genome ID for each assembly is referenced in Table S1.

## Acknowledgments.

This project was conducted with support by a grant from the National Science Foundation (No. DEB-1749206). The soil warming experiments at Harvard Forest are maintained with support from the National Science Foundation Long Term Ecological Research Program (DEB-1832110) and a Long Term Research in Environmental Biology grant (DEB-1456610). We are grateful to Drs. Serita Frey and Mel Knorr for maintaining this infrastructure for our community. This work was also conducted with support (proposal 507977: 10.46936/10.25585/60008103 to M.J.C. and K.M.D.) from the U.S. Department of Energy Joint Genome Institute (https://ror.org/04xm1d337), a DOE Office of Science User Facility supported by the Office of Science of the U.S. Department of Energy operated under Contract No. DE-AC02-05CH11231. We are grateful to all the people who contributed to the isolation, sequencing, and annotation of isolates in this study, including (but probably not limited to) Erin Bergeron, Andrew Billings, Isabella Bushko, Gina Chaput, Emily Clark, Spencer Moore, Samantha Murphy, Grace Pold, Bianca Surjawan, and Wing Yin Tam. We also thank Adam Breister, Akieliah Robinson, and Madaris Serrano Perez for their helpful feedback on the manuscript.

## Author Contributions Statement

**Choudoir, MJ**: Conceptualization, Formal analysis, Writing - Original Draft, Visualization; **Narayanan, A**: Methodology, Resources; **Rodriquez-Ramos, D**: Methodology, Resources; **Simoes, R**: Resources; **Efroni, A**: Resources; **Sondrini, A**: Resources; **DeAngelis, KM**: Conceptualization, Writing - Review & Editing, Supervision, Funding acquisition.

