## Supplementary Information for "Pangenomes reveal genomic signatures of microbial adaptation to experimental soil warming"

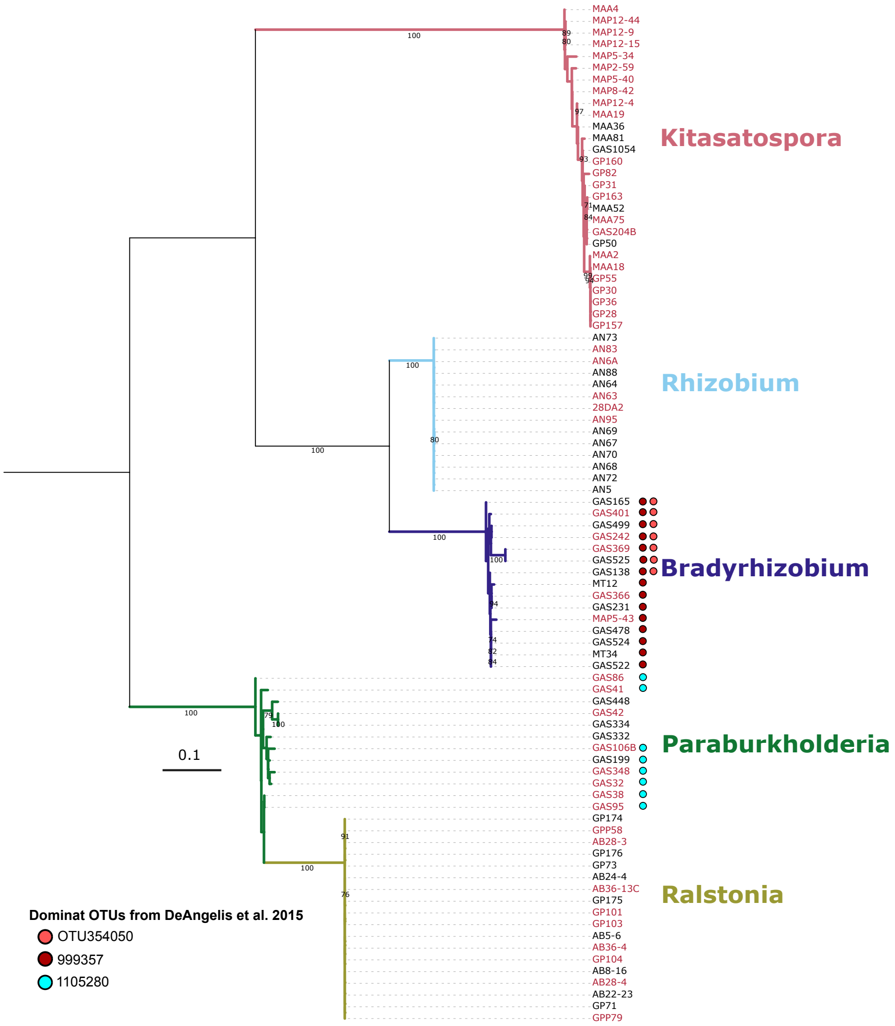


**Figure S1. 16S rRNA gene phylogeny**. Tree shows the phylogenetic relationships between all 91 isolates (Table S1) constructed from 16S rRNA gene nucleotide sequences derived from anvi'o HMM searches. Genome names are colored according to treatment group, with heated in red and control in black. Phylogenetic clades are also labeled and colored according to their pangenome group. Scale bar indicates nucleotide substitutions per site. Bootstrap values greater than 70 from 100 iterations are noted on tree branches. Tree is mid-rooted. Note, *Paraburkholderia* genomes GAS33, GAS82, GAS205, and GAS206C did not have 16S rRNA gene hits, so these strains are omitted from the tree. Circles indicate 16S rRNA gene sequences with > 99% identity to partial 16S rRNA gene sequences (254 bp) from dominant subset community (n=155 OTUs, rank abundance) from a previous amplicon study conducted at the Harvard Forest warming experiment (DeAngelis *et al.*, 2015). Alphaproteobacteria taxa NewOTU354050 and 999357 had greater relative abundances in heated communities, while Betaproteobacteria taxon 1105280 had greater relative abundances in control communities. None of these OTUs were indicator taxa.

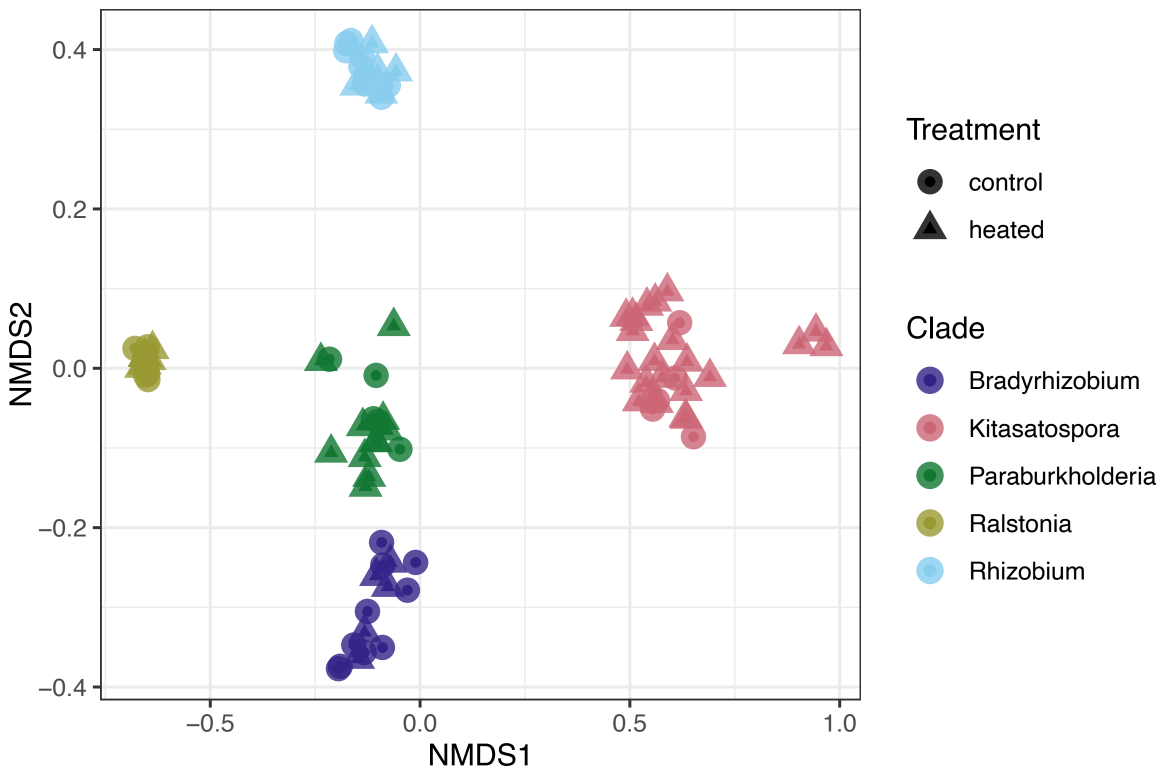


**Figure S2. CAZyme composition across clades.** Plot shows non-metric multidimensional scaling (NMDS) ordination of CAZyme composition (Bray-Curtis distances) across all genomes. Each point represents the collection of carbohydrate-active enzymes annotated with the dbCAN HMM database (HMMdb release 11.0) (Zhang *et al*., 2018; Zheng *et al*., 2023) for each genome. Points are colored by clade and shapes correspond to warming experiment treatment according to legend. Clade membership, but not treatment or the interaction between treatment and clade, was a strong predictor for CAZyme composition (PERMANOVA, P-value=0.001, $R^2$=0.85).

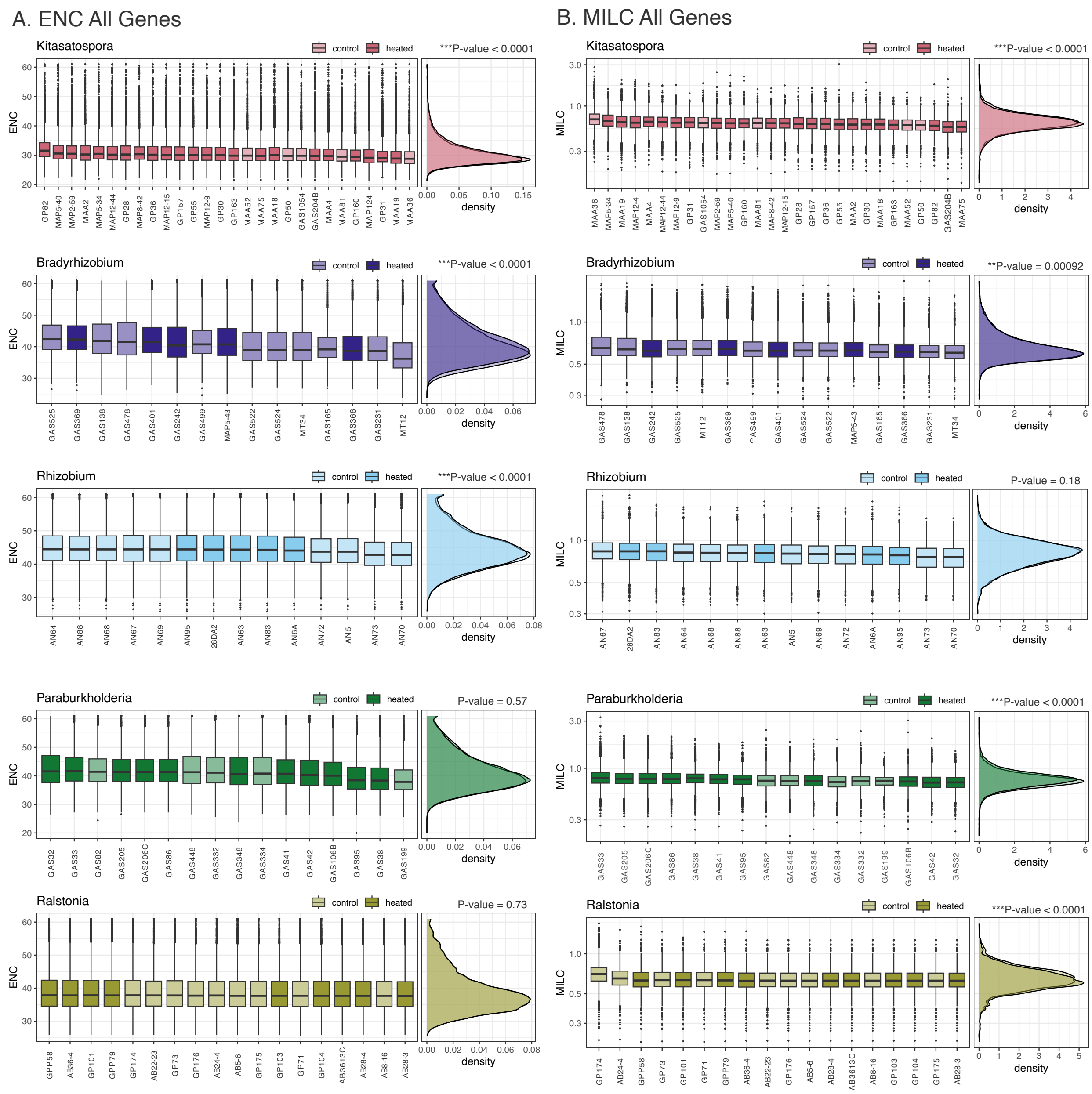


**Figure S3. Gene-level genome-wide codon usage distributions.** Panels illustrate global gene-level codon usage distributions for all strains across clades. Each boxplot shows the interquartile range of codon usage values for all genes, and the line is the median, whiskers are 1.5 interquartile range, and black points are outliers. Boxplots are colored by clade and treatment. Plots to the right show the codon usage density distributions by treatment. Figure S3A shows ENC (Wright, 1990) distributions, and Figure S3B shows MILC (Supek & Vlahoviček, 2005) distributions. We calculated MILC values against a set of conserved ribosomal protein genes. Wilcox rank sum test P-values for differences in codon usage between warming treatment are reported.

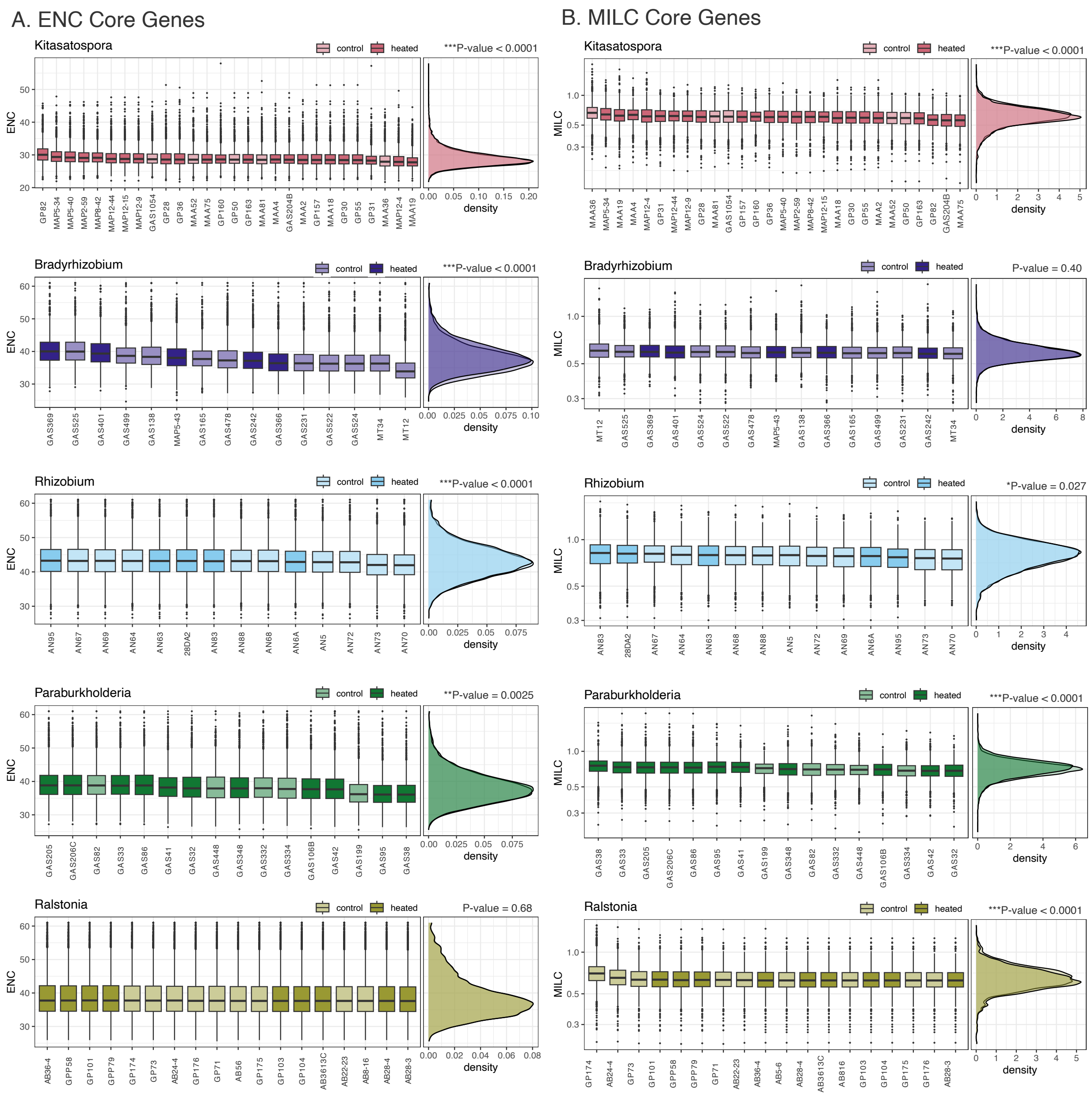


**Figure S4. Gene-level core genome codon usage distributions.** Panels illustrate gene-level core genome codon usage distributions for all strains across all clades. Each boxplot shows the interquartile range of codon usage values for core genes, and the line is the median, whiskers are 1.5 interquartile range, and black points are outliers. Boxplots are colored by clade and treatment. Plots to the right show the codon usage density distributions by treatment. Figure S4A shows ENC (Wright, 1990) distributions, and Figure S4B shows MILC (Supek & Vlahoviček, 2005) distributions. We calculated MILC values against a set of conserved ribosomal protein genes. Wilcox rank sum test P-values for differences in codon usage between warming treatment are reported.

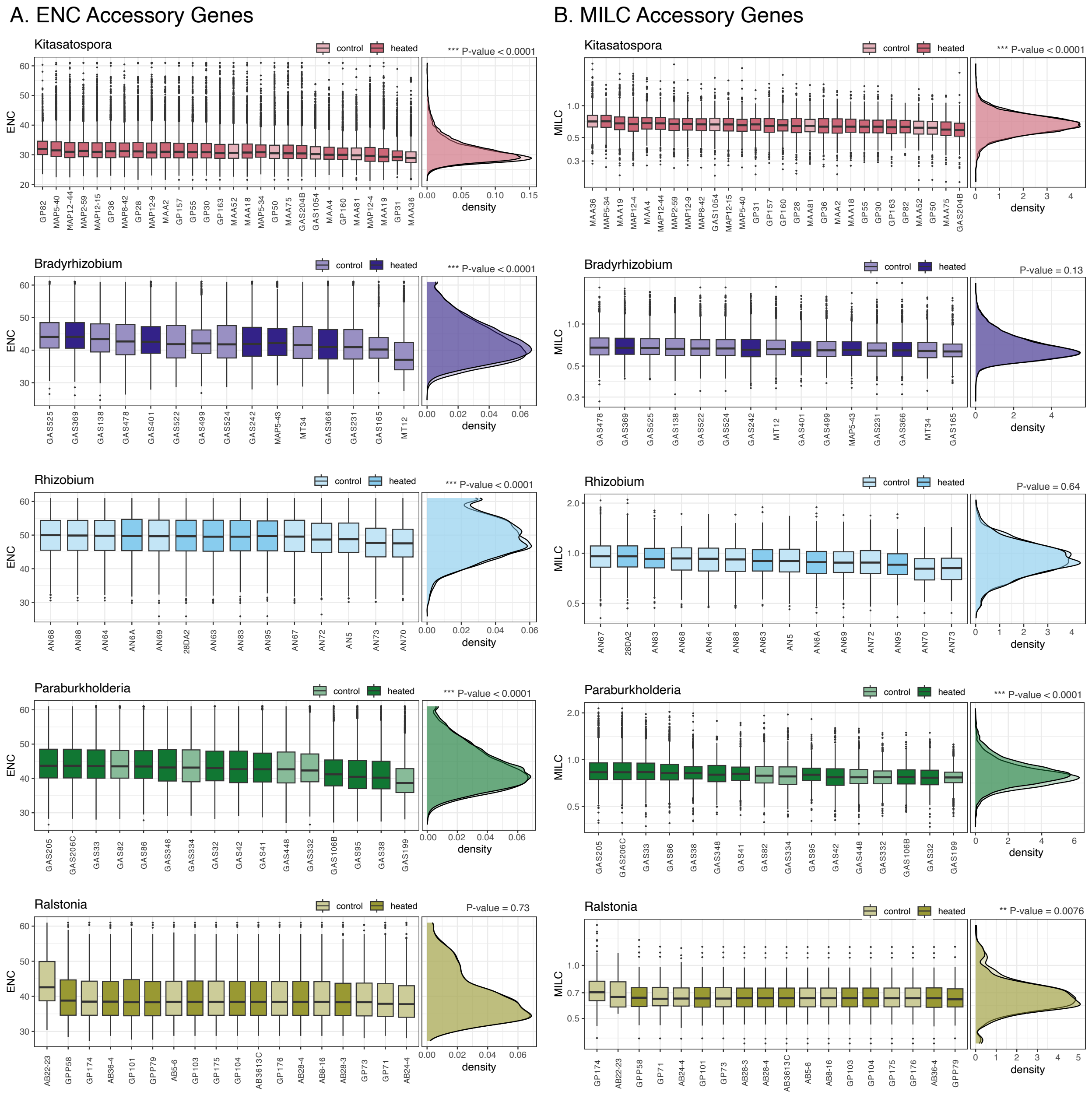


**Figure S5. Gene-level accessory genome codon usage distributions.** Panels illustrate the gene-level accessory genome codon usage distributions for all strains across all clades. Note, singletons genes were not included in the accessory gene pool for this analysis. Each boxplot shows the interquartile range of codon usage values for accessory genes, and the line is the median, whiskers are 1.5 interquartile range, and black points are outliers. Boxplots are colored by clade and treatment. Plots to the right show the codon usage density distributions by treatment. Figure S5A shows ENC (Wright, 1990) distributions, and Figure S5B shows MILC (Supek & Vlahoviček, 2005) distributions. We calculated MILC values against a set of conserved ribosomal protein genes. Wilcox rank sum test P-values for differences in codon usage between warming treatment are reported.

**Table S1. Isolation metadata for bacterial isolates.** The 91 isolates in this study include *Kitasatospora* spp., *Bradyrhizobium* spp., *Paraburkholderia* spp., and *Rhizobia* spp. clades. Columns report isolate ID, phylum, organism name, warming experiment treatment, soil horizon (na values are unknown), original isolation media, year isolated, genome sequencing platform, sequencing facility, NCBI BioSample accession number, and IMG taxon ID of the genome assembly. Media abbreviations include: VL55 amended with xylan, plant ploymers (PP), or readily oxidized carbon (ROC); humic acid vitamin agar (HVA); lignin soybean flour vitamin agar (LSFA); modified carboxymethyl cellulose (CMC); International *Streptomyces* Project-2 Medium (ISP2); and 1% glucose/0.4% potato infusion (1% glu). See references within (Pold *et al.*, 2016; Domeignoz-Horta *et al.*, 2020) for media formulations and additional isolation conditions. Note the 20 genomes in the sequencing facility column that were previously sequenced, indicated with asterisks. All other genomes were generated for this study.

**Table S2. Genome metadata for draft genome assemblies.** Columns report genome ID, pangenome clade, warming experiment treatment, number of contigs in the assembly, total assembly length, percent G+C, percent estimated genome completeness and redundancy (plus estimated confidence), total number of genes, average gene length, number of genes per kb, number of unique or singleton gene clusters, total number of gene clusters, 16S rRNA gene number, and the total number of carbohydrate-active enzyme (i.e., CAZymes) annotations. We used the program anvi-estimate-genome-completeness to estimate genome completion and redundancy based on presence of single-copy genes. We used the program anvi-run-hmms to determine 16S rRNA gene copy numbers with Hidden Markov Model (HMM) searches to anvi'o HMM sources Ribosomal_RNA_16S. We used anvi-run-cazymes to run HMM searches against dbCAN HMMdb v11 (Zhang *et al.*, 2018; Zheng *et al.*, 2023).

**Table S3. ONT minION assembly metadata and quality control metrics.** For all draft genomes sequenced and assembled in house using ONT minION, columns report isolate ID, phylum, pangenome clade, *de novo* or hybrid assembly (see Methods), assembly coverage (X), taxonomic classification determined by CheckM (Parks *et al.*, 2015), assembly length, percent G+C, number of contigs, N50, and completeness and contamination as estimated by CheckM.

**Table S4. Functional enrichment metrics for KOfam annotated gene clusters.** Columns are KOfam descriptions, enrichment score (see Methods and Shaiber *et al.*, 2020), unadjusted P-values, Q-values adjusting for multiple tests, associated warming experiment treatment, KO identifier, corresponding gene clusters, the portion of heated (p_Heated) and control (p_Control) genomes that harbor each functional annotation, and clade. Note, only functional annotations with unadjusted P-values < 0.05 are reported.

**Table S5. Functional enrichment metrics for KEGG annotated gene clusters.** Columns are KEGG module descriptions, enrichment score (see Methods and Shaiber *et al.*, 2020), unadjusted

P-values, Q-values adjusting for multiple tests, associated warming experiment treatment, KEGG identifier, corresponding gene clusters, the portion of heated (p_Heated) and control (p_Control) genomes that harbor each functional annotation, and clade. Note, only functional annotations with unadjusted P-values < 0.05 are reported.

**Table S6. Functional enrichment metrics for complete metabolic modules.** Columns are KEGG complete metabolic module descriptions (Kanehisa *et al.*, 2012), enrichment score (see Methods and Shaiber *et al*., 2020), unadjusted P-values, Q-values adjusting for multiple tests, associated warming experiment treatment, KEGG metabolic module identifier, corresponding genomes, the portion of heated (p_Heated) and control (p_Control) genomes that harbor each metabolic module, and clade. Note, only metabolic modules with unadjusted P-values < 0.05 are reported.

**Table S7. Functional enrichment metrics for carbohydrate-active enzymes (CAZymes)**. Columns are CAZyme HMM name, enzyme class, family activity description and EC number, enzyme substrate, enrichment score (see Methods and Shaiber *et al*., 2020), unadjusted P-values, Q-values adjusting for multiple tests, associated warming experiment treatment, predicted substrate, corresponding gene clusters, the portion of control (p_Control) and heated (p_Heated) genomes that harbor each CAZyme annotation, and clade. Note, only CAZyme annotations with unadjusted P-values < 0.05 are reported. Enzyme descriptions were parsed from the Carbohydrate-Active enZYmes Database webpages of www.cazy.org

### Table S1. Isolate Metadata

| Isolate | Phylum | Organism Name | Treatment | Horizon | Isolation Media | Isolation Date | Sequence Platform | Sequencing Facility | Biosample | IMG taxon ID |
|---|---|---|---|---|---|---|---|---|---|---|
| Actinobacteria *Kitasatospora* spp. clade | | | | | | | | | | |
| GAS1054 | Actinobacteria | Kitasatospora sp. GAS1054 | control | mineral | VL55 PP | 2014 | ONT MinION | DeAngelis Lab | SAMN33769326 | 2958935987 |
| GAS204B | Actinobacteria | Kitasatospora sp. GAS204B | heated | mineral | VL55 PP | 2014 | Illumina NovaSeq S4 | DOE JGI | SAMN33769410 | 2981703317 |
| GP28 | Actinobacteria | Kitasatospora sp. GP28 | heated | organic | oatmeal | 2014 | ONT MinION | DeAngelis Lab | SAMN33771170 | 2953957014 |
| GP30 | Actinobacteria | Kitasatospora sp. GP30 | heated | organic | oatmeal | 2014 | Illumina NovaSeq S4 | DOE JGI | SAMN33771171 | 2981846400 |
| GP31 | Actinobacteria | Kitasatospora sp. GP31 | heated | organic | oatmeal | 2014 | ONT MinION | DeAngelis Lab | SAMN33771172 | 2955925447 |
| GP36 | Actinobacteria | Kitasatospora sp. GP36 | heated | organic | oatmeal | 2014 | ONT MinION | DeAngelis Lab | SAMN33771173 | 2940444706 |
| GP50 | Actinobacteria | Kitasatospora sp. GP50 | control | mineral | oatmeal | 2014 | ONT MinION | DeAngelis Lab | SAMN33771174 | 2953929336 |
| GP55 | Actinobacteria | Kitasatospora sp. GP55 | heated | mineral | oatmeal | 2014 | PacBio RS, PacBio RS II | DOE JGI* | SAMN04515641 | 2617270923 |
| GP82 | Actinobacteria | Kitasatospora sp. GP82 | heated | mineral | oatmeal | 2014 | Illumina NovaSeq S4 | DOE JGI | SAMN33771176 | 2982040100 |
| GP157 | Actinobacteria | Kitasatospora sp. GP157 | heated | organic | HVA | 2014 | ONT MinION | DeAngelis Lab | SAMN33769410 | 2958916086 |
| GP160 | Actinobacteria | Kitasatospora sp. GP160 | heated | mineral | HVA | 2014 | ONT MinION | DeAngelis Lab | SAMN33771167 | 2958909016 |
| GP163 | Actinobacteria | Kitasatospora sp. GP163 | heated | mineral | LSFV | 2014 | ONT MinION | DeAngelis Lab | SAMN33771168 | 2953937211 |
| MAA2 | Actinobacteria | Kitasatospora sp. MAA2 | heated | organic | VL55 xylan | 2020 | ONT MinION | DeAngelis Lab | SAMN33771178 | 2953912599 |
| MAA4 | Actinobacteria | Kitasatospora sp. MAA4 | heated | organic | VL55 xylan | 2020 | Illumina NovaSeq S4 | DOE JGI | SAMN33771180 | 2982008629 |
| MAA18 | Actinobacteria | Streptomycetaceae bacterium MAA18 | heated | mineral | VL55 xylan | 2020 | Illumina NovaSeq S4 | DOE JGI | SAMN33771207 | 2981232351 |
| MAA19 | Actinobacteria | Kitasatospora sp. MAA19 | heated | mineral | VL55 xylan | 2020 | Illumina NovaSeq S4 | DOE JGI | SAMN33771177 | 2960241103 |
| MAA36 | Actinobacteria | Kitasatospora sp. MAA36 | control | organic | VL55 xylan | 2020 | ONT MinION | DeAngelis Lab | SAMN33771179 | 2958928483 |
| MAA52 | Actinobacteria | Kitasatospora sp. MAA52 | control | mineral | VL55 xylan | 2020 | ONT MinION | DeAngelis Lab | SAMN33771181 | 2940436776 |
| MAA75 | Actinobacteria | Kitasatospora sp. MAA75 | heated | organic | ISP2 | 2020 | ONT MinION | DeAngelis Lab | SAMN33771182 | 2953921507 |
| MAA81 | Actinobacteria | Kitasatospora sp. MAA81 | control | organic | ISP2 | 2020 | ONT MinION | DeAngelis Lab | SAMN33771183 | 2953904123 |
| MAP2-59 | Actinobacteria | Kitasatospora sp. MAP2-59 | heated | mineral | VL55 ROC | 2020 | ONT MinION | DeAngelis Lab | SAMN33771188 | 2958901192 |
| MAP5-34 | Actinobacteria | Kitasatospora sp. MAP5-34 | heated | mineral | ISP2 | 2020 | Illumina NovaSeq S4 | DOE JGI | SAMN33771189 | 2963575613 |
| MAP5-40 | Actinobacteria | Kitasatospora sp. MAP5-40 | heated | mineral | ISP2 | 2020 | ONT MinION | DeAngelis Lab | SAMN33771190 | 2953966083 |
| MAP8-42 | Actinobacteria | Kitasatospora sp. MAP8-42 | heated | mineral | VL55 ROC | 2020 | ONT MinION | DeAngelis Lab | SAMN33771191 | 2955909675 |
| MAP12-4 | Actinobacteria | Kitasatospora sp. MAP12-4 | heated | mineral | oatmeal | 2020 | ONT MinION | DeAngelis Lab | SAMN33771185 | 2939854634 |
| MAP12-9 | Actinobacteria | Kitasatospora sp. MAP12-9 | heated | mineral | oatmeal | 2020 | Illumina NovaSeq S4 | DOE JGI | SAMN33771187 | 2963582266 |
| MAP12-15 | Actinobacteria | Kitasatospora sp. MAP12-15 | heated | mineral | oatmeal | 2020 | Illumina NovaSeq S4 | DOE JGI | SAMN33771184 | 2982032058 |
| MAP12-44 | Actinobacteria | Kitasatospora sp. MAP12-44 | heated | organic | oatmeal | 2020 | PacBio Sequel IIe | DOE JGI | SAMN33771186 | 2974460287 |
| Alphaproteobacteria *Bradyrhizobium* spp. clade | | | | | | | | | | |
| GAS138 | Alphaproteobacteria | Bradyrhizobium erythrophlei GAS138 | control | mineral | VL55 PP | 2014 | PacBio RS, PacBio RS II | DOE JGI* | SAMN05443248 | 2695421015 |
| GAS165 | Alphaproteobacteria | Bradyrhizobium lablabi GAS165 | control | mineral | VL55 PP | 2014 | PacBio RS, PacBio RS II | DOE JGI* | SAMN05877939 | 2747843221 |
| GAS231 | Alphaproteobacteria | Afipia sp. GAS231 | control | mineral | VL55 PP | 2014 | PacBio RS, PacBio RS II | DOE JGI* | SAMN05444050 | 2690315678 |
| GAS242 | Alphaproteobacteria | Bradyrhizobium erythrophlei GAS242 | heated | mineral | VL55 PP | 2014 | PacBio RS, PacBio RS II | DOE JGI* | SAMN05444169 | 2695420948 |
| GAS366 | Alphaproteobacteria | Bradyrhizobium sp. GAS366 | heated | mineral | VL55 PP | 2014 | ONT MinION | DeAngelis Lab | SAMN33771221 | 2929879271 |
| GAS369 | Alphaproteobacteria | Bradyrhizobium canariense GAS369 | heated | mineral | VL55 PP | 2014 | PacBio RS, PacBio RS II | DOE JGI* | SAMN05444158 | 2693430033 |
| GAS401 | Alphaproteobacteria | Bradyrhizobium erythrophlei GAS401 | heated | na | VL55 PP | 2014 | PacBio RS, PacBio RS II | DOE JGI* | SAMN05444170 | 2695420919 |
| GAS478 | Alphaproteobacteria | Bradyrhizobium erythrophlei GAS478 | control | mineral | VL55 PP | 2014 | PacBio RS, PacBio RS II | DOE JGI* | SAMN05443247 | 2698537050 |
| GAS499 | Alphaproteobacteria | Bradyrhizobium lablabi GAS499 | control | mineral | VL55 PP | 2014 | PacBio RS, PacBio RS II | DOE JGI* | SAMN05444159 | 2698536816 |
| GAS522 | Alphaproteobacteria | Bradyrhizobium lablabi GAS522 | control | mineral | VL55 PP | 2014 | PacBio RS, PacBio RS II | DOE JGI* | SAMN05444171 | 2693429786 |
| GAS524 | Alphaproteobacteria | Bradyrhizobium ottawaense GAS524 | control | mineral | VL55 PP | 2014 | PacBio RS, PacBio RS II | DOE JGI* | SAMN05444163 | 2693430034 |
| GAS525 | Alphaproteobacteria | Afipia broomeae GAS525 | control | mineral | VL55 PP | 2014 | PacBio RS, PacBio RS II | DOE JGI* | SAMN05880572 | 2740892596 |
| MAP5-43 | Alphaproteobacteria | Bradyrhizobium sp. MAP5-43 | heated | mineral | ISP2 | 2020 | ONT MinION | DeAngelis Lab | SAMN33771222 | 2930805785 |
| MT12 | Alphaproteobacteria | Bradyrhizobium erythrophlei MT12 | control | organic | VL55 xylan | 2014 | PacBio RS, PacBio RS II | DOE JGI* | SAMN05444164 | 2690316366 |
| MT34 | Alphaproteobacteria | Bradyrhizobium lablabi MT34 | control | organic | VL55 xylan | 2014 | PacBio RS, PacBio RS II | DOE JGI* | SAMN05444321 | 2698536699 |

#### Table S1. Isolate Metadata

**Alphaproteobacteria *Rhizobium* spp. clade**

| | | | | | | | | | | |
|---|---|---|---|---|---|---|---|---|---|---|
| 28DA2 | Alphaproteobacteria | Rhizoboum sp. 28DA2 | heated | subsurface | BioSep; Bandounas ligni | 2013 | ONT MinION | DeAngelis Lab | SAMN33771228 | 2929580477 |
| AN5 | Alphaproteobacteria | Rhizobium sp. AN5 | control | surface | BioSep; Bandounas ligni | 2013 | PacBio RS, PacBio RS II | DOE JGI* | SAMN05216358 | 2617270923 |
| AN63 | Alphaproteobacteria | Rhizoboum sp. AN63 | heated | subsurface | BioSep; Bandounas ligni | 2013 | ONT MinION | DeAngelis Lab | SAMN33771229 | 2929873235 |
| AN64 | Alphaproteobacteria | Rhizoboum sp. AN64 | control | subsurface | BioSep; Bandounas ligni | 2013 | ONT MinION | DeAngelis Lab | SAMN33771230 | 2929597866 |
| AN67 | Alphaproteobacteria | Rhizoboum sp. AN67 | control | surface | BioSep; Bandounas ligni | 2013 | ONT MinION | DeAngelis Lab | SAMN33771231 | 2930909959 |
| AN68 | Alphaproteobacteria | Rhizobium sp. AN68 | control | surface | BioSep; Bandounas ligni | 2013 | ONT MinION | DeAngelis Lab | SAMN33771224 | 2935258951 |
| AN69 | Alphaproteobacteria | Rhizoboum sp. AN69 | control | surface | BioSep; Bandounas ligni | 2013 | ONT MinION | DeAngelis Lab | SAMN33771232 | 2935252894 |
| AN6A | Alphaproteobacteria | Rhizobium sp. AN6A | heated | subsurface | BioSep; Bandounas ligni | 2013 | PacBio RS, PacBio RS II | DOE JGI* | SAMN05216595 | 2619618868 |
| AN70 | Alphaproteobacteria | Rhizobium sp. AN70 | control | surface | BioSep; Bandounas ligni | 2013 | Illumina NovaSeq S4 | DOE JGI | SAMN33771226 | 2960282700 |
| AN72 | Alphaproteobacteria | Rhizoboum sp. AN72 | control | surface | BioSep; Bandounas ligni | 2013 | ONT MinION | DeAngelis Lab | SAMN33771233 | 2929586531 |
| AN73 | Alphaproteobacteria | Rhizobium sp. AN73 | control | surface | BioSep; Bandounas ligni | 2013 | ONT MinION | DeAngelis Lab | SAMN33771227 | 2935264491 |
| AN83 | Alphaproteobacteria | Rhizoboum sp. AN83 | heated | subsurface | BioSep; Bandounas ligni | 2013 | ONT MinION | DeAngelis Lab | SAMN33771234 | 2929591855 |
| AN88 | Alphaproteobacteria | Rhizoboum sp. AN88 | control | subsurface | BioSep; Bandounas ligni | 2013 | ONT MinION | DeAngelis Lab | SAMN33771235 | 2929574867 |
| AN95 | Alphaproteobacteria | Rhizoboum sp. AN95 | heated | subsurface | BioSep; Bandounas ligni | 2013 | ONT MinION | DeAngelis Lab | SAMN33771236 | 2930903789 |

**Betaproteobacteria *Paraburkholderia* spp. clade**

| | | | | | | | | | | |
|---|---|---|---|---|---|---|---|---|---|---|
| GAS32 | Betaproteobacteria | Paraburkholderia sp. GAS32 | heated | mineral | VL55 ROC | 2014 | Illumina NovaSeq S4 | DOE JGI | SAMN33771245 | 2981470370 |
| GAS33 | Betaproteobacteria | Paraburkholderia sp. GAS33 | heated | mineral | VL55 ROC | 2014 | Illumina NovaSeq S4 | DOE JGI | SAMN33771246 | 2981462429 |
| GAS38 | Betaproteobacteria | Paraburkholderia sp. GAS38 | heated | mineral | VL55 ROC | 2014 | Illumina NovaSeq S4 | DOE JGI | SAMN33771249 | 2981752396 |
| GAS41 | Betaproteobacteria | Paraburkholderia sp. GAS41 | heated | mineral | VL55 PP | 2014 | Illumina NovaSeq S4 | DOE JGI | SAMN33771250 | 2981392040 |
| GAS42 | Betaproteobacteria | Paraburkholderia sp. GAS42 | heated | mineral | VL55 PP | 2014 | Illumina NovaSeq S4 | DOE JGI | SAMN33771251 | 2981399414 |
| GAS82 | Betaproteobacteria | Paraburkholderia sp. GAS82 | control | mineral | VL55 PP | 2014 | Illumina NovaSeq S4 | DOE JGI | SAMN33771253 | 2981406600 |
| GAS86 | Betaproteobacteria | Paraburkholderia phenazinium GAS86 | heated | mineral | VL55 xylan | 2014 | PacBio RS II | DOE JGI* | SAMN05444168 | 2695421038 |
| GAS95 | Betaproteobacteria | Paraburkholderia phenazinium GAS95 | heated | mineral | modified CMC | 2014 | PacBio RS II | DOE JGI* | SAMN05444165 | 2695420309 |
| GAS106B | Betaproteobacteria | Paraburkholderia fungorum GAS106B | heated | mineral | modified CMC | 2014 | PacBio RS II | DOE JGI* | SAMN05443245 | 2690315676 |
| GAS199 | Betaproteobacteria | Paraburkholderia sp. GAS199 | control | mineral | VL55 PP | 2014 | Illumina NovaSeq S4 | DOE JGI | SAMN33771242 | 2981432666 |
| GAS205 | Betaproteobacteria | Paraburkholderia sp. GAS205 | heated | mineral | VL55 PP | 2014 | Illumina NovaSeq S4 | DOE JGI | SAMN33771243 | 2981440061 |
| GAS206C | Betaproteobacteria | Paraburkholderia sp. GAS206C | heated | mineral | VL55 PP | 2014 | Illumina NovaSeq S4 | DOE JGI | SAMN33771244 | 2981759879 |
| GAS332 | Betaproteobacteria | Burkholderia sp. GAS332 | control | mineral | modified CMC | 2014 | PacBio RS II | DOE JGI* | SAMN05444172 | 2695420918 |
| GAS334 | Betaproteobacteria | Paraburkholderia sp. GAS334 | control | mineral | VL55 PP | 2014 | Illumina NovaSeq S4 | DOE JGI | SAMN33771247 | 2981447645 |
| GAS348 | Betaproteobacteria | Paraburkholderia sp. GAS348 | heated | mineral | VL55 PP | 2014 | Illumina NovaSeq S4 | DOE JGI | SAMN33771248 | 2981455594 |
| GAS448 | Betaproteobacteria | Paraburkholderia sp. GAS448 | control | mineral | VL55 ROC | 2014 | Illumina NovaSeq S4 | DOE JGI | SAMN33771252 | 2981774849 |

**Betaproteobacteria *Ralstonia* spp. clade**

| | | | | | | | | | | |
|---|---|---|---|---|---|---|---|---|---|---|
| AB5-6 | Betaproteobacteria | Ralstonia sp. AB5-6 | control | organic | 1% nutrient broth | 2014 | ONT MinION, Illumina NextSeq 2000 | DeAngelis Lab, SeqCenter | SAMN33771260 | 2966226929 |
| AB8-16 | Betaproteobacteria | Ralstonia sp. AB8-16 | control | organic | 1% nutrient broth | 2014 | ONT MinION, Illumina NextSeq 2000 | DeAngelis Lab, SeqCenter | SAMN33771261 | 2966232277 |
| AB22-23 | Betaproteobacteria | Ralstonia sp. AB22-23 | control | mineral | 1% nutrient broth | 2014 | ONT MinION, Illumina NextSeq 2000 | DeAngelis Lab, SeqCenter | SAMN33771254 | 2966237621 |
| AB24-4 | Betaproteobacteria | Ralstonia sp. AB24-4 | control | mineral | 1% nutrient broth | 2014 | ONT MinION | DeAngelis Lab | SAMN33771255 | 2923570319 |
| AB28-3 | Betaproteobacteria | Ralstonia sp. AB28-3 | heated | mineral | 1% nutrient broth | 2014 | ONT MinION, Illumina NextSeq 2000 | DeAngelis Lab, SeqCenter | SAMN33771256 | 2966242759 |
| AB28-4 | Betaproteobacteria | Ralstonia sp. AB28-4 | heated | mineral | 1% nutrient broth | 2014 | ONT MinION, Illumina NextSeq 2000 | DeAngelis Lab, SeqCenter | SAMN33771257 | 2966248084 |
| AB36-4 | Betaproteobacteria | Ralstonia sp. AB36-4 | heated | mineral | 1% nutrient broth | 2014 | ONT MinION | DeAngelis Lab | SAMN33771259 | 2928862943 |
| AB36-13C | Betaproteobacteria | Ralstonia sp. AB36-13C | heated | mineral | 1% nutrient broth | 2014 | ONT MinION, Illumina NextSeq 2000 | DeAngelis Lab, SeqCenter | SAMN33771258 | 2966253427 |
| GP71 | Betaproteobacteria | Ralstonia sp. GP71 | control | organic | VL55 PP | 2014 | ONT MinION | DeAngelis Lab | SAMN33771268 | 2923564856 |
| GP73 | Betaproteobacteria | Ralstonia sp. GP73 | control | mineral | water agar | 2014 | ONT MinION | DeAngelis Lab | SAMN33771269 | 2923725988 |
| GP101 | Betaproteobacteria | Ralstonia sp. GP101 | heated | organic | VL55 PP | 2014 | ONT MinION | DeAngelis Lab | SAMN33771262 | 2923597253 |
| GP103 | Betaproteobacteria | Ralstonia sp. GP103 | heated | organic | VL55 PP | 2014 | ONT MinION, Illumina NextSeq 2000 | DeAngelis Lab, SeqCenter | SAMN33771263 | 2966258773 |
| GP104 | Betaproteobacteria | Ralstonia sp. GP104 | heated | organic | VL55 PP | 2014 | ONT MinION, Illumina NextSeq 2000 | DeAngelis Lab, SeqCenter | SAMN33771264 | 2966264122 |

#### Table S1. Isolate Metadata

| | | | | | | | | | | |
|---|---|---|---|---|---|---|---|---|---|---|
| GP174 | Betaproteobacteria | Ralstonia sp. GP174 | control | organic | VL55 PP | 2014 | ONT MinION | DeAngelis Lab | SAMN33771265 | 2923731573 |
| GP175 | Betaproteobacteria | Ralstonia sp. GP175 | control | mineral | water agar | 2014 | ONT MinION, Illumina NextSeq 2000 | DeAngelis Lab, SeqCenter | SAMN33771266 | 2966269469 |
| GP176 | Betaproteobacteria | Ralstonia sp. GP176 | control | mineral | water agar | 2014 | ONT MinION, Illumina NextSeq 2000 | DeAngelis Lab, SeqCenter | SAMN33771267 | 2966274816 |
| GPP58 | Betaproteobacteria | Ralstonia sp. GPP58 | heated | mineral | VL55 PP | 2014 | ONT MinION | DeAngelis Lab | SAMN33771270 | 2927050446 |
| GPP79 | Betaproteobacteria | Ralstonia sp. GPP79 | heated | mineral | VL55 PP | 2014 | ONT MinION | DeAngelis Lab | SAMN33771271 | 2923720395 |

#### Table S2. Genome Metadata

| Genome | Clade | Treatment | Contigs | Total Length | % G+C | % Completion | % Redundancy | Confidence | Genes | Ave Gene Length | Genes per Kb | Singleton GCs | Total GCs | 16S rRNA Genes | CAZymes |
|---|---|---|---|---|---|---|---|---|---|---|---|---|---|---|---|
| GAS1054 | Kitasatospora | control | 2 | 9526358 | 72.3 | 100 | 4.23 | 1 | 8632 | 945 | 0.91 | 643 | 7308 | 10 | 326 |
| GAS204B | Kitasatospora | heated | 84 | 8423638 | 71.9 | 98.6 | 8.45 | 1 | 7319 | 1014 | 0.87 | 47 | 6790 | 1 | 297 |
| GP157 | Kitasatospora | heated | 4 | 8969559 | 71.5 | 98.6 | 9.86 | 1 | 8156 | 954 | 0.91 | 19 | 7408 | 10 | 281 |
| GP160 | Kitasatospora | heated | 2 | 8159895 | 72.3 | 98.6 | 2.82 | 1 | 6762 | 1051 | 0.83 | 298 | 6127 | 10 | 240 |
| GP163 | Kitasatospora | heated | 5 | 8638857 | 71.8 | 98.6 | 7.04 | 0.9 | 7588 | 990 | 0.88 | 50 | 6841 | 9 | 308 |
| GP28 | Kitasatospora | heated | 4 | 8987849 | 71.5 | 97.2 | 9.86 | 1 | 8618 | 890 | 0.96 | 85 | 7471 | 10 | 306 |
| GP30 | Kitasatospora | heated | 110 | 8861088 | 71.6 | 98.6 | 7.04 | 1 | 7986 | 973 | 0.90 | 19 | 7399 | 1 | 271 |
| GP31 | Kitasatospora | heated | 2 | 9246900 | 72.7 | 98.6 | 8.45 | 1 | 7975 | 1014 | 0.86 | 908 | 7229 | 10 | 307 |
| GP36 | Kitasatospora | heated | 8 | 8071464 | 71.5 | 97.2 | 12.7 | 1 | 7918 | 867 | 0.98 | 88 | 6758 | 10 | 274 |
| GP50 | Kitasatospora | control | 2 | 8590657 | 71.8 | 98.6 | 8.45 | 1 | 7578 | 988 | 0.88 | 60 | 6846 | 9 | 299 |
| GP55 | Kitasatospora | heated | 3 | 8962713 | 71.5 | 98.6 | 7.04 | 1 | 7990 | 979 | 0.89 | 3 | 7388 | 10 | 270 |
| GP82 | Kitasatospora | heated | 138 | 8204036 | 70.7 | 100 | 8.45 | 1 | 7384 | 938 | 0.90 | 1599 | 6746 | 1 | 264 |
| MAA18 | Kitasatospora | heated | 81 | 8759053 | 71.7 | 98.6 | 7.04 | 1 | 7764 | 988 | 0.89 | 121 | 7170 | 1 | 267 |
| MAA19 | Kitasatospora | heated | 96 | 9091641 | 72.6 | 100 | 7.04 | 1 | 8119 | 968 | 0.89 | 437 | 7468 | 1 | 289 |
| MAA2 | Kitasatospora | heated | 6 | 9209497 | 71.4 | 98.6 | 7.04 | 1 | 8532 | 928 | 0.93 | 360 | 7627 | 10 | 288 |
| MAA36 | Kitasatospora | control | 3 | 7481034 | 73.1 | 98.6 | 5.63 | 1 | 7089 | 866 | 0.95 | 1061 | 6306 | 10 | 256 |
| MAA4 | Kitasatospora | heated | 71 | 8171757 | 71.7 | 98.6 | 7.04 | 1 | 7330 | 998 | 0.90 | 998 | 6713 | 1 | 284 |
| MAA52 | Kitasatospora | control | 3 | 8580784 | 71.8 | 98.6 | 8.45 | 1 | 7624 | 973 | 0.89 | 48 | 6804 | 9 | 304 |
| MAA75 | Kitasatospora | heated | 2 | 8524674 | 71.8 | 100 | 8.45 | 1 | 7538 | 984 | 0.88 | 190 | 6886 | 9 | 297 |
| MAA81 | Kitasatospora | control | 5 | 8784971 | 72.2 | 100 | 7.04 | 1 | 8106 | 935 | 0.92 | 1096 | 7063 | 9 | 322 |
| MAP12-15 | Kitasatospora | heated | 100 | 8934013 | 71.5 | 98.6 | 8.45 | 1 | 7785 | 1018 | 0.87 | 3 | 7147 | 1 | 296 |
| MAP12-4 | Kitasatospora | heated | 6 | 8976678 | 72.5 | 100 | 7.04 | 1 | 8514 | 882 | 0.95 | 457 | 7381 | 10 | 317 |
| MAP12-44 | Kitasatospora | heated | 4 | 9043878 | 71.4 | 98.6 | 8.45 | 1 | 7830 | 1019 | 0.87 | 17 | 7149 | 10 | 298 |
| MAP12-9 | Kitasatospora | heated | 72 | 8102036 | 71.6 | 98.6 | 4.23 | 1 | 7055 | 1019 | 0.87 | 1 | 6553 | 1 | 263 |
| MAP2-59 | Kitasatospora | heated | 9 | 8170709 | 71.1 | 100 | 11.3 | 1 | 7476 | 943 | 0.91 | 281 | 6705 | 9 | 355 |
| MAP5-34 | Kitasatospora | heated | 118 | 7318731 | 71.2 | 98.6 | 9.86 | 1 | 6448 | 985 | 0.88 | 982 | 5959 | 1 | 262 |
| MAP5-40 | Kitasatospora | heated | 7 | 7915022 | 71.2 | 97.2 | 14.1 | 0.9 | 7635 | 880 | 0.96 | 178 | 6643 | 9 | 393 |
| MAP8-42 | Kitasatospora | heated | 12 | 7813028 | 71.2 | 100 | 12.7 | 1 | 7149 | 946 | 0.92 | 92 | 6428 | 9 | 371 |
| GAS138 | Bradyrhizobium | control | 1 | 9092036 | 61.4 | 100 | 2.82 | 1 | 8434 | 883 | 0.93 | 1663 | 7532 | 1 | 206 |
| GAS165 | Bradyrhizobium | control | 3 | 6132387 | 62.5 | 100 | 2.82 | 1 | 5820 | 916 | 0.95 | 702 | 5492 | 1 | 155 |
| GAS231 | Bradyrhizobium | control | 1 | 7584136 | 62.6 | 100 | 2.82 | 1 | 7021 | 933 | 0.93 | 11 | 6516 | 1 | 182 |
| GAS242 | Bradyrhizobium | heated | 1 | 9184651 | 61.9 | 100 | 0 | 1 | 8622 | 876 | 0.94 | 1401 | 7637 | 1 | 202 |
| GAS366 | Bradyrhizobium | heated | 1 | 7583959 | 62.6 | 100 | 8.45 | 1 | 7240 | 897 | 0.95 | 74 | 6564 | 1 | 188 |

#### Table S2. Genome Metadata

| Genome | Clade | Treatment | Contigs | Total Length | % G+C | % Completion | % Redundancy | Confidence | Genes | Ave Gene Length | Genes per Kb | Singleton GCs | Total GCs | 16S rRNA Genes | CAZymes |
|---|---|---|---|---|---|---|---|---|---|---|---|---|---|---|---|
| GAS369 | Bradyrhizobium | heated | 1 | 7841944 | 60.9 | 100 | 7.04 | 1 | 7252 | 919 | 0.92 | 261 | 6737 | 1 | 174 |
| GAS401 | Bradyrhizobium | heated | 1 | 7525117 | 61.2 | 100 | 2.82 | 1 | 7107 | 893 | 0.94 | 1492 | 6508 | 1 | 184 |
| GAS478 | Bradyrhizobium | control | 4 | 11738562 | 61.4 | 100 | 2.82 | 1 | 11048 | 850 | 0.94 | 2137 | 8725 | 1 | 231 |
| GAS499 | Bradyrhizobium | control | 1 | 7909999 | 61.8 | 100 | 0 | 1 | 7330 | 917 | 0.93 | 954 | 6766 | 1 | 213 |
| GAS522 | Bradyrhizobium | control | 2 | 8269569 | 62.3 | 100 | 8.45 | 1 | 7801 | 896 | 0.94 | 67 | 6980 | 1 | 187 |
| GAS524 | Bradyrhizobium | control | 1 | 8339115 | 62.3 | 100 | 8.45 | 1 | 7921 | 890 | 0.95 | 136 | 7106 | 1 | 186 |
| GAS525 | Bradyrhizobium | control | 1 | 7951160 | 60.8 | 100 | 4.23 | 1 | 7353 | 920 | 0.92 | 417 | 6847 | 1 | 174 |
| MAP5-43 | Bradyrhizobium | heated | 1 | 8047392 | 62.1 | 98.6 | 1.41 | 1 | 7660 | 888 | 0.95 | 1300 | 6918 | 2 | 189 |
| MT12 | Bradyrhizobium | control | 2 | 8967172 | 63.8 | 100 | 5.63 | 1 | 8411 | 905 | 0.94 | 1741 | 7703 | 1 | 212 |
| MT34 | Bradyrhizobium | control | 1 | 8150868 | 62.3 | 100 | 7.04 | 1 | 7657 | 901 | 0.94 | 165 | 6901 | 1 | 190 |
| AN5 | Rhizobium | control | 5 | 5527901 | 58.6 | 100 | 0 | 1 | 5255 | 931 | 0.95 | 412 | 5095 | 4 | 173 |
| AN63 | Rhizobium | heated | 4 | 5617997 | 58.6 | 98.6 | 1.41 | 1 | 5767 | 851 | 1.03 | 30 | 5143 | 4 | 191 |
| AN64 | Rhizobium | control | 3 | 5639358 | 58.5 | 100 | 0 | 1 | 5413 | 917 | 0.96 | 30 | 5119 | 5 | 179 |
| AN67 | Rhizobium | control | 4 | 5619457 | 58.6 | 98.6 | 0 | 1 | 5885 | 830 | 1.05 | 54 | 5163 | 4 | 204 |
| AN68 | Rhizobium | control | 3 | 5639523 | 58.5 | 100 | 1.41 | 1 | 5375 | 924 | 0.95 | 19 | 5106 | 5 | 176 |
| AN69 | Rhizobium | control | 4 | 5618682 | 58.6 | 100 | 1.41 | 1 | 5834 | 839 | 1.04 | 30 | 5147 | 4 | 199 |
| AN6A | Rhizobium | heated | 4 | 5625739 | 58.6 | 100 | 0 | 1 | 5241 | 950 | 0.93 | 10 | 5036 | 4 | 173 |
| AN70 | Rhizobium | control | 22 | 5208341 | 59.2 | 100 | 1.41 | 1 | 4904 | 945 | 0.94 | 13 | 4750 | 1 | 158 |
| AN72 | Rhizobium | control | 3 | 5470029 | 58.8 | 100 | 0 | 1 | 5144 | 938 | 0.94 | 198 | 4906 | 4 | 176 |
| AN73 | Rhizobium | control | 3 | 5252559 | 59.1 | 100 | 2.82 | 1 | 5030 | 921 | 0.96 | 37 | 4771 | 5 | 166 |
| AN83 | Rhizobium | heated | 4 | 5617845 | 58.6 | 94.4 | 2.82 | 1 | 5748 | 851 | 1.02 | 42 | 5134 | 4 | 198 |
| AN88 | Rhizobium | control | 3 | 5642017 | 58.5 | 100 | 0 | 1 | 5420 | 916 | 0.96 | 26 | 5120 | 5 | 176 |
| AN95 | Rhizobium | heated | 4 | 5616373 | 58.6 | 97.2 | 1.41 | 1 | 5888 | 829 | 1.05 | 50 | 5160 | 4 | 206 |
| 28DA2 | Rhizobium | heated | 4 | 5618214 | 58.6 | 95.8 | 1.41 | 1 | 5793 | 845 | 1.03 | 46 | 5159 | 4 | 192 |
| GAS106B | Paraburkholderia | heated | 4 | 8581175 | 61.2 | 98.6 | 1.41 | 1 | 7431 | 974 | 0.87 | 1248 | 6882 | 8 | 230 |
| GAS199 | Paraburkholderia | control | 20 | 8196845 | 62.4 | 100 | 1.41 | 1 | 7166 | 986 | 0.87 | 992 | 6569 | 1 | 216 |
| GAS205 | Paraburkholderia | heated | 35 | 8284543 | 61.4 | 100 | 4.23 | 1 | 7360 | 975 | 0.89 | 1 | 6848 | 0 | 221 |
| GAS206C | Paraburkholderia | heated | 30 | 8284727 | 61.4 | 100 | 4.23 | 1 | 7353 | 976 | 0.89 | 1 | 6844 | 0 | 221 |
| GAS32 | Paraburkholderia | heated | 141 | 10460609 | 61.1 | 100 | 1.41 | 1 | 9732 | 905 | 0.93 | 1450 | 8485 | 1 | 239 |
| GAS33 | Paraburkholderia | heated | 38 | 8627607 | 61.3 | 98.6 | 4.23 | 1 | 7694 | 968 | 0.89 | 407 | 7118 | 0 | 228 |
| GAS332 | Paraburkholderia | control | 3 | 10416547 | 61.4 | 100 | 2.82 | 1 | 9124 | 972 | 0.88 | 1511 | 8113 | 7 | 257 |
| GAS334 | Paraburkholderia | control | 102 | 8203652 | 61.6 | 100 | 1.41 | 1 | 7624 | 904 | 0.93 | 719 | 6714 | 1 | 185 |
| GAS348 | Paraburkholderia | heated | 79 | 7262714 | 61.2 | 100 | 2.82 | 1 | 6476 | 942 | 0.89 | 829 | 5856 | 1 | 182 |

#### Table S2. Genome Metadata

| Genome | Clade | Treatment | Contigs | Total Length | % G+C | % Completion | % Redundancy | Confidence | Genes | Ave Gene Length | Genes per Kb | Singleton GCs | Total GCs | 16S rRNA Genes | CAZymes |
|---|---|---|---|---|---|---|---|---|---|---|---|---|---|---|---|
| GAS38 | Paraburkholderia | heated | 34 | 8268867 | 63.0 | 100 | 4.23 | 1 | 7262 | 987 | 0.88 | 255 | 6788 | 1 | 223 |
| GAS41 | Paraburkholderia | heated | 92 | 7839433 | 61.8 | 100 | 2.82 | 1 | 7094 | 937 | 0.90 | 838 | 6400 | 1 | 198 |
| GAS42 | Paraburkholderia | heated | 69 | 7534538 | 61.7 | 100 | 1.41 | 1 | 6921 | 923 | 0.92 | 557 | 6261 | 1 | 176 |
| GAS448 | Paraburkholderia | control | 168 | 10103218 | 61.9 | 100 | 1.41 | 1 | 9458 | 894 | 0.94 | 1816 | 8202 | 1 | 243 |
| GAS82 | Paraburkholderia | control | 31 | 8375761 | 61.4 | 100 | 4.23 | 1 | 7415 | 979 | 0.89 | 309 | 6885 | 0 | 227 |
| GAS86 | Paraburkholderia | heated | 2 | 8319467 | 61.4 | 100 | 4.23 | 1 | 7256 | 988 | 0.87 | 161 | 6729 | 5 | 224 |
| GAS95 | Paraburkholderia | heated | 3 | 8526942 | 62.8 | 100 | 4.23 | 1 | 7407 | 992 | 0.87 | 249 | 6855 | 6 | 228 |
| AB22-23 | Ralstonia | control | 7 | 5298867 | 63.6 | 94.4 | 2.82 | 0.9 | 4960 | 945 | 0.94 | 2 | 4724 | 3 | 117 |
| AB24-4 | Ralstonia | control | 4 | 5432636 | 63.6 | 98.6 | 2.82 | 1 | 5236 | 906 | 0.96 | 21 | 4861 | 3 | 125 |
| AB28-3 | Ralstonia | heated | 6 | 5483600 | 63.6 | 100 | 2.82 | 1 | 5133 | 944 | 0.94 | 1 | 4911 | 3 | 121 |
| AB28-4 | Ralstonia | heated | 5 | 5513759 | 63.6 | 100 | 2.82 | 1 | 5159 | 945 | 0.94 | 0 | 4913 | 3 | 121 |
| AB36-13C | Ralstonia | heated | 5 | 5513759 | 63.6 | 100 | 2.82 | 1 | 5159 | 945 | 0.94 | 0 | 4912 | 3 | 121 |
| AB36-4 | Ralstonia | heated | 6 | 5579439 | 63.5 | 100 | 5.63 | 1 | 5304 | 923 | 0.95 | 11 | 4916 | 3 | 122 |
| AB5-6 | Ralstonia | control | 5 | 5513753 | 63.6 | 100 | 2.82 | 1 | 5164 | 944 | 0.94 | 0 | 4914 | 3 | 121 |
| AB8-16 | Ralstonia | control | 5 | 5513543 | 63.6 | 100 | 2.82 | 1 | 5159 | 945 | 0.94 | 0 | 4913 | 3 | 121 |
| GP101 | Ralstonia | heated | 5 | 5513656 | 63.6 | 98.6 | 4.23 | 1 | 5359 | 899 | 0.97 | 21 | 4912 | 3 | 125 |
| GP103 | Ralstonia | heated | 6 | 5515304 | 63.6 | 100 | 2.82 | 1 | 5162 | 944 | 0.94 | 0 | 4914 | 3 | 121 |
| GP104 | Ralstonia | heated | 6 | 5515238 | 63.6 | 100 | 2.82 | 1 | 5161 | 944 | 0.94 | 0 | 4914 | 3 | 121 |
| GP174 | Ralstonia | control | 5 | 5513800 | 63.6 | 98.6 | 4.23 | 1 | 5352 | 899 | 0.97 | 25 | 4926 | 3 | 126 |
| GP175 | Ralstonia | control | 6 | 5515256 | 63.6 | 100 | 2.82 | 1 | 5162 | 944 | 0.94 | 0 | 4914 | 3 | 121 |
| GP176 | Ralstonia | control | 18 | 5553901 | 63.6 | 100 | 2.82 | 1 | 5211 | 942 | 0.94 | 24 | 4940 | 3 | 121 |
| GP71 | Ralstonia | control | 4 | 5432770 | 63.6 | 98.6 | 4.23 | 1 | 5201 | 915 | 0.96 | 15 | 4851 | 3 | 126 |
| GP73 | Ralstonia | control | 5 | 5477586 | 63.6 | 98.6 | 5.63 | 1 | 5311 | 899 | 0.97 | 16 | 4908 | 3 | 129 |
| GPP58 | Ralstonia | heated | 5 | 5513689 | 63.6 | 98.6 | 8.45 | 1 | 5346 | 902 | 0.97 | 23 | 4931 | 3 | 125 |
| GPP79 | Ralstonia | heated | 5 | 5513696 | 63.6 | 98.6 | 4.23 | 1 | 5318 | 906 | 0.96 | 17 | 4921 | 3 | 126 |

#### Table S3. MinION Assembly Stats

| Isolate | Phylum | Clade | Assembly Method | Coverage (X) | CheckM Lineage | Assembly Length | % G+C | Contigs | N50 | Completeness | Contamination |
|---|---|---|---|---|---|---|---|---|---|---|---|
| GAS1054 | Actinobacteria | Kitasatospora | de novo | 37.8 | o__Actinomycetales | 9526358 | 72.3 | 2 | 9465601 | 95.7 | 2.9 |
| GP28 | Actinobacteria | Kitasatospora | de novo | 40.1 | o__Actinomycetales | 8987849 | 71.5 | 4 | 8885772 | 98.2 | 0.8 |
| GP31 | Actinobacteria | Kitasatospora | de novo | 38.9 | o__Actinomycetales | 9246900 | 72.7 | 2 | 9218386 | 99.3 | 3.2 |
| GP36 | Actinobacteria | Kitasatospora | de novo | 44.6 | o__Actinomycetales | 8071464 | 71.5 | 8 | 2279891 | 95.0 | 0.8 |
| GP50 | Actinobacteria | Kitasatospora | de novo | 41.9 | o__Actinomycetales | 8590657 | 71.8 | 2 | 8097869 | 99.5 | 0.9 |
| GP157 | Actinobacteria | Kitasatospora | de novo | 40.1 | o__Actinomycetales | 8969559 | 71.5 | 4 | 8878309 | 99.5 | 0.8 |
| GP160 | Actinobacteria | Kitasatospora | de novo | 44.1 | o__Actinomycetales | 8159895 | 72.3 | 2 | 6036722 | 99.0 | 3.4 |
| GP163 | Actinobacteria | Kitasatospora | de novo | 41.7 | o__Actinomycetales | 8638857 | 71.8 | 5 | 8084037 | 99.0 | 0.9 |
| MAA2 | Actinobacteria | Kitasatospora | de novo | 39.1 | o__Actinomycetales | 9209497 | 71.4 | 6 | 8574763 | 97.6 | 1.6 |
| MAA36 | Actinobacteria | Kitasatospora | de novo | 42.8 | o__Actinomycetales | 7481034 | 73.1 | 3 | 7229582 | 94.7 | 2.1 |
| MAA52 | Actinobacteria | Kitasatospora | de novo | 47.2 | o__Actinomycetales | 8580784 | 71.8 | 3 | 7998586 | 98.4 | 1.4 |
| MAA75 | Actinobacteria | Kitasatospora | de novo | 42.2 | o__Actinomycetales | 8524674 | 71.8 | 2 | 8012833 | 100.0 | 1.4 |
| MAA81 | Actinobacteria | Kitasatospora | de novo | 41.0 | o__Actinomycetales | 8784971 | 72.2 | 5 | 8512193 | 97.0 | 0.7 |
| MAP2-59 | Actinobacteria | Kitasatospora | de novo | 44.1 | o__Actinomycetales | 8170709 | 71.1 | 9 | 8011956 | 99.5 | 1.2 |
| MAP5-40 | Actinobacteria | Kitasatospora | de novo | 45.5 | o__Actinomycetales | 7915022 | 71.2 | 7 | 7591491 | 97.6 | 0.6 |
| MAP8-42 | Actinobacteria | Kitasatospora | de novo | 46.1 | o__Actinomycetales | 7813028 | 71.2 | 12 | 6256654 | 98.8 | 0.6 |
| MAP12-4 | Actinobacteria | Kitasatospora | de novo | 45.1 | o__Actinomycetales | 8976678 | 72.5 | 6 | 8304589 | 96.9 | 2.4 |
| GAS366 | Alphaproteobacteria | Bradyrhizobium | de novo | 32.0 | f__Bradyrhizobiaceae | 7583959 | 62.6 | 1 | 7583959 | 99.3 | 1.4 |
| MAP5-43 | Alphaproteobacteria | Bradyrhizobium | de novo | 41.9 | f__Bradyrhizobiaceae | 8047392 | 62.1 | 1 | 8047392 | 99.3 | 0.9 |
| 28DA2 | Alphaproteobacteria | Rhizobium | de novo | 32.0 | f__Rhizobiaceae | 5618214 | 58.6 | 4 | 2869860 | 96.8 | 0.1 |
| AN63 | Alphaproteobacteria | Rhizobium | de novo | 44.9 | f__Rhizobiaceae | 5617997 | 58.6 | 4 | 2869899 | 97.4 | 0.1 |
| AN64 | Alphaproteobacteria | Rhizobium | de novo | 31.9 | f__Rhizobiaceae | 5639358 | 58.5 | 3 | 2110257 | 99.2 | 0.8 |
| AN67 | Alphaproteobacteria | Rhizobium | de novo | 44.8 | f__Rhizobiaceae | 5619457 | 58.6 | 4 | 2870042 | 97.6 | 0.1 |
| AN68 | Alphaproteobacteria | Rhizobium | de novo | 44.7 | f__Rhizobiaceae | 5639523 | 58.5 | 3 | 2110415 | 99.6 | 0.7 |
| AN69 | Alphaproteobacteria | Rhizobium | de novo | 44.9 | f__Rhizobiaceae | 5618682 | 58.6 | 4 | 2870003 | 97.8 | 0.1 |
| AN72 | Alphaproteobacteria | Rhizobium | de novo | 32.9 | f__Rhizobiaceae | 5470029 | 58.8 | 3 | 2915437 | 99.8 | 0.6 |
| AN73 | Alphaproteobacteria | Rhizobium | de novo | 48.0 | f__Rhizobiaceae | 5252559 | 59.2 | 3 | 2912879 | 99.5 | 0.7 |
| AN83 | Alphaproteobacteria | Rhizobium | de novo | 32.0 | f__Rhizobiaceae | 5617845 | 58.6 | 4 | 2869877 | 98.6 | 0.1 |
| AN88 | Alphaproteobacteria | Rhizobium | de novo | 31.9 | f__Rhizobiaceae | 5642017 | 58.5 | 3 | 2112953 | 98.7 | 0.8 |
| AN95 | Alphaproteobacteria | Rhizobium | de novo | 44.9 | f__Rhizobiaceae | 5616373 | 58.6 | 4 | 2870026 | 97.3 | 0.1 |
| AB5-6 | Betaproteobacteria | Ralstonia | hybrid | 174.7 | f__Burkholderiaceae | 5513753 | 63.6 | 5 | 3537649 | 99.9 | 0.5 |
| AB8-16 | Betaproteobacteria | Ralstonia | hybrid | 146.8 | f__Burkholderiaceae | 5513543 | 63.6 | 5 | 3537481 | 99.9 | 0.5 |
| AB22-23 | Betaproteobacteria | Ralstonia | hybrid | 184.9 | f__Burkholderiaceae | 5298867 | 63.6 | 7 | 1386635 | 92.6 | 0.5 |
| AB24-4 | Betaproteobacteria | Ralstonia | de novo | 39.8 | f__Burkholderiaceae | 5432636 | 63.6 | 4 | 3537574 | 99.0 | 0.5 |

#### Table S3. MinION Assembly Stats

| Isolate | Phylum | Clade | Assembly Method | Coverage (X) | CheckM Lineage | Assembly Length | % G+C | Contigs | N50 | Completeness | Contamination |
|---|---|---|---|---|---|---|---|---|---|---|---|
| AB36-4 | Betaproteobacteria | Ralstonia | de novo | 38.7 | f__Burkholderiaceae | 5579439 | 63.5 | 6 | 3564194 | 98.8 | 0.5 |
| GP71 | Betaproteobacteria | Ralstonia | de novo | 39.8 | f__Burkholderiaceae | 5432770 | 63.6 | 4 | 3537633 | 99.1 | 0.5 |
| GP73 | Betaproteobacteria | Ralstonia | de novo | 39.4 | f__Burkholderiaceae | 5477586 | 63.6 | 5 | 3537750 | 97.9 | 0.5 |
| GP101 | Betaproteobacteria | Ralstonia | de novo | 39.2 | f__Burkholderiaceae | 5513656 | 63.6 | 5 | 3537632 | 98.9 | 0.5 |
| GP104 | Betaproteobacteria | Ralstonia | hybrid | 187.3 | f__Burkholderiaceae | 5515238 | 63.6 | 6 | 3537649 | 99.9 | 0.5 |
| GP174 | Betaproteobacteria | Ralstonia | de novo | 39.2 | f__Burkholderiaceae | 5513800 | 63.6 | 5 | 3537701 | 98.2 | 0.5 |
| GP176 | Betaproteobacteria | Ralstonia | hybrid | 190.9 | f__Burkholderiaceae | 5553306 | 63.6 | 16 | 1386635 | 99.9 | 0.5 |
| GPP58 | Betaproteobacteria | Ralstonia | de novo | 39.2 | f__Burkholderiaceae | 5513689 | 63.6 | 5 | 3537615 | 98.5 | 0.5 |
| GPP79 | Betaproteobacteria | Ralstonia | de novo | 39.2 | f__Burkholderiaceae | 5512301 | 63.6 | 5 | 3537617 | 98.7 | 0.5 |

#### Table S4. KOfam Functional Enrichment

| Description | Enrichment Score | P-value | Q-value | Associated Treatment | KO Identifier | Gene Cluster IDs | p_Heated | p_Control | Clade |
|---|---|---|---|---|---|---|---|---|---|
| nicotinamide mononucleotide transporter | 9.91 | 0.002 | 0.75 | heated | K03811 | GC_00002777 | 1.0 | 0.6 | Kitasatospora |
| gentisate 1,2-dioxygenase [EC:1.13.11.4] | 8.12 | 0.004 | 0.75 | control | K00450 | GC_00005029 | 0.3 | 1.0 | Kitasatospora |
| (5-formylfuran-3-yl)methyl phosphate synthase [EC:4.2.3.153] | 8.12 | 0.004 | 0.75 | control | K09733 | GC_00002980 | 0.3 | 1.0 | Kitasatospora |
| pentalenene oxygenase [EC:1.14.15.32] | 8.12 | 0.004 | 0.75 | control | K15907 | GC_00006887, GC_00010110, GC_00016724, GC_00029715 | 0.3 | 1.0 | Kitasatospora |
| Rrf2 family transcriptional regulator, repressor of oqxAB | 7.89 | 0.005 | 0.75 | control | K19587 | GC_00007190 | 0.2 | 0.8 | Kitasatospora |
| cell division protease FtsH [EC:3.4.24.-] | 7.02 | 0.008 | 0.75 | control | K03798 | GC_00005770, GC_00006673 | 0.3 | 1.0 | Kitasatospora |
| thiamine kinase [EC:2.7.1.89] | 7.02 | 0.008 | 0.75 | control | K07251 | GC_00004527 | 0.3 | 1.0 | Kitasatospora |
| toxin FitB [EC:3.1.-.-] | 7.02 | 0.008 | 0.75 | control | K07062 | GC_00004283, GC_00020036 | 0.3 | 1.0 | Kitasatospora |
| maleate isomerase [EC:5.2.1.1] | 7.02 | 0.008 | 0.75 | control | K01799 | GC_00005486, GC_00028536, GC_00028579 | 0.3 | 1.0 | Kitasatospora |
| polysaccharide biosynthesis protein PslG | 6.39 | 0.011 | 0.75 | control | K21000 | GC_00007371, GC_00017393 | 0.2 | 0.8 | Kitasatospora |
| magnesium-dependent phosphatase 1 [EC:3.1.3.48 3.1.3.-] | 6.39 | 0.011 | 0.75 | control | K17619 | GC_00008076, GC_00017122 | 0.2 | 0.8 | Kitasatospora |
| arsenite oxidase small subunit [EC:1.20.2.1 1.20.9.1] | 6.39 | 0.011 | 0.75 | control | K08355 | GC_00006620 | 0.2 | 0.8 | Kitasatospora |
| acyl-CoA oxidase [EC:1.3.3.6] | 6.39 | 0.011 | 0.75 | control | K00232 | GC_00007179, GC_00009968, GC_00013461, GC_00023709 | 0.2 | 0.8 | Kitasatospora |
| RpiR family transcriptional regulator, carbohydrate utilization regulator | 6.39 | 0.011 | 0.75 | control | K19337 | GC_00006408 | 0.2 | 0.8 | Kitasatospora |
| small membrane protein | 6.39 | 0.011 | 0.75 | control | K09153 | GC_00006341 | 0.2 | 0.8 | Kitasatospora |
| antitoxin Phd | 6.39 | 0.011 | 0.75 | heated | K19165 | GC_00003864, GC_00009763 | 0.8 | 0.2 | Kitasatospora |
| dimethylglycine oxidase [EC:1.5.3.10] | 6.39 | 0.011 | 0.75 | heated | K00309 | GC_00003488 | 0.8 | 0.2 | Kitasatospora |
| glutamate transport system ATP-binding protein [EC:7.4.2.1] | 6.39 | 0.011 | 0.75 | heated | K10008 | GC_00000458 | 0.8 | 0.2 | Kitasatospora |
| glutamate transport system substrate-binding protein | 6.39 | 0.011 | 0.75 | heated | K10005 | GC_00002724 | 0.8 | 0.2 | Kitasatospora |
| serine O-acetyltransferase [EC:2.3.1.30] | 6.09 | 0.014 | 0.75 | heated | K00640 | GC_00004630, GC_00025323, GC_00029649 | 0.6 | 0.0 | Kitasatospora |
| NTF2-related export protein 1/2 | 5.46 | 0.019 | 0.75 | control | K14285 | GC_00015502 | 0.0 | 0.4 | Kitasatospora |
| heme exporter protein B | 5.46 | 0.019 | 0.75 | control | K02194 | GC_00016395 | 0.0 | 0.4 | Kitasatospora |
| L-proline 4-hydroxylase [EC:1.14.11.57] | 5.46 | 0.019 | 0.75 | control | K21615 | GC_00024079, GC_00025558, GC_00028750 | 0.0 | 0.4 | Kitasatospora |
| prephenate decarboxylase [EC:4.1.1.100] | 5.46 | 0.019 | 0.75 | control | K19546 | GC_00016699 | 0.0 | 0.4 | Kitasatospora |
| damage-control phosphatase, subfamily III [EC:3.1.3.-] | 5.46 | 0.019 | 0.75 | heated | K23114 | GC_00003704, GC_00008583 | 1.0 | 0.6 | Kitasatospora |
| zinc/manganese transport system ATP-binding protein | 5.46 | 0.019 | 0.75 | heated | K02074 | GC_00000338 | 1.0 | 0.6 | Kitasatospora |
| pilus assembly protein CpaB | 5.46 | 0.019 | 0.75 | heated | K02279 | GC_00003359, GC_00009250, GC_00017467 | 1.0 | 0.6 | Kitasatospora |
| phosphoenolpyruvate phosphomutase [EC:5.4.2.9] | 5.38 | 0.020 | 0.75 | control | K01841 | GC_00010589, GC_00028165 | 0.1 | 0.6 | Kitasatospora |
| 2-hydroxy-6-oxonona-2,4-dienedioate hydrolase [EC:3.7.1.14] | 5.38 | 0.020 | 0.75 | control | K05714 | GC_00009176 | 0.1 | 0.6 | Kitasatospora |
| 2,3-dihydroxyphenylpropionate 1,2-dioxygenase [EC:1.13.11.16] | 5.38 | 0.020 | 0.75 | control | K05713 | GC_00008123 | 0.1 | 0.6 | Kitasatospora |
| phosphonopyruvate decarboxylase [EC:4.1.1.82] | 5.38 | 0.020 | 0.75 | control | K09459 | GC_00010304, GC_00021033 | 0.1 | 0.6 | Kitasatospora |
| hercynylcysteine S-oxide lyase [EC:4.4.1.36] | 5.38 | 0.020 | 0.75 | control | K18913 | GC_00009766 | 0.1 | 0.6 | Kitasatospora |
| starvation-inducible DNA-binding protein | 5.38 | 0.020 | 0.75 | heated | K04047 | GC_00003474, GC_00025296 | 0.9 | 0.4 | Kitasatospora |
| ATP-dependent RNA helicase DeaD [EC:3.6.4.13] | 5.38 | 0.020 | 0.75 | heated | K05592 | GC_00003429 | 0.9 | 0.4 | Kitasatospora |
| putative chitinase | 5.28 | 0.022 | 0.75 | control | K03791 | GC_00003830 | 0.4 | 1.0 | Kitasatospora |
| TetR/AcrR family transcriptional regulator, fatty acid metabolism regulator protein | 5.20 | 0.023 | 0.75 | control | K13770 | GC_00005998 | 0.3 | 0.8 | Kitasatospora |

#### Table S4. KOfam Functional Enrichment

| Description | Enrichment Score | P-value | Q-value | Associated Treatment | KO Identifier | Gene Cluster IDs | p_Heated | p_Control | Clade |
|---|---|---|---|---|---|---|---|---|---|
| alpha-N-acetylglucosaminidase [EC:3.2.1.50] | 5.20 | 0.023 | 0.75 | control | K01205 | GC_00005322 | 0.3 | 0.8 | Kitasatospora |
| peptidoglycan DL-endopeptidase RipA [EC:3.4.-.-] | 5.20 | 0.023 | 0.75 | control | K21473 | GC_00005972 | 0.3 | 0.8 | Kitasatospora |
| polyprenyl-phospho-N-acetylgalactosaminyl synthase | 5.20 | 0.023 | 0.75 | control | K22907 | GC_00005503 | 0.3 | 0.8 | Kitasatospora |
| protein HIRA/HIR1 | 5.20 | 0.023 | 0.75 | control | K11293 | GC_00005634, GC_00017673 | 0.3 | 0.8 | Kitasatospora |
| colanic acid/amylovoran biosynthesis protein WcaK/AmsJ | 5.20 | 0.023 | 0.75 | control | K16710 | GC_00007252, GC_00010530 | 0.3 | 0.8 | Kitasatospora |
| 5-carboxymethyl-2-hydroxymuconate isomerase [EC:5.3.3.10] | 5.20 | 0.023 | 0.75 | heated | K01826 | GC_00004867, GC_00015586, GC_00016351, GC_00018400, GC_00019132 | 0.7 | 0.2 | Kitasatospora |
| protein PhnA | 5.20 | 0.023 | 0.75 | heated | K06193 | GC_00003716 | 0.7 | 0.2 | Kitasatospora |
| putative ATP-dependent endonuclease of the OLD family | 5.20 | 0.023 | 0.75 | heated | K07459 | GC_00010889, GC_00015153, GC_00016745, GC_00018242, GC_00022573, GC_00023176, GC_00024225, GC_00024358, GC_00024735, GC_00028275 | 0.7 | 0.2 | Kitasatospora |
| polyether ionophore transport system permease protein | 5.20 | 0.023 | 0.75 | heated | K25149 | GC_00005968, GC_00009675, GC_00017311 | 0.7 | 0.2 | Kitasatospora |
| polyether ionophore transport system ATP-binding protein | 5.20 | 0.023 | 0.75 | heated | K25150 | GC_00003620 | 0.7 | 0.2 | Kitasatospora |
| gamma-polyglutamate biosynthesis protein CapC | 4.77 | 0.029 | 0.75 | control | K22116 | GC_00030495 | 0.0 | 0.2 | Kitasatospora |
| lactate permease | 4.77 | 0.029 | 0.75 | control | K03303 | GC_00029165 | 0.0 | 0.2 | Kitasatospora |
| streptogrisin C [EC:3.4.21.-] | 4.77 | 0.029 | 0.75 | control | K18546 | GC_00021229 | 0.0 | 0.2 | Kitasatospora |
| C3 family ADP-ribosyltransferase [EC:2.4.2.-] | 4.77 | 0.029 | 0.75 | control | K11044 | GC_00028645 | 0.0 | 0.2 | Kitasatospora |
| 2-methylcitrate dehydratase [EC:4.2.1.79] | 4.77 | 0.029 | 0.75 | control | K01720 | GC_00027771 | 0.0 | 0.2 | Kitasatospora |
| ATP-dependent helicase IRC3 [EC:5.6.2.-] | 4.77 | 0.029 | 0.75 | control | K17677 | GC_00027236 | 0.0 | 0.2 | Kitasatospora |
| iron(III)-enterobactin esterase [EC:3.1.1.108] | 4.77 | 0.029 | 0.75 | control | K07214 | GC_00030704 | 0.0 | 0.2 | Kitasatospora |
| gamma-polyglutamate synthase [EC:6.3.2.-] | 4.77 | 0.029 | 0.75 | control | K01932 | GC_00021332 | 0.0 | 0.2 | Kitasatospora |
| cobalamin transport system substrate-binding protein | 4.77 | 0.029 | 0.75 | control | K25034 | GC_00025894 | 0.0 | 0.2 | Kitasatospora |
| extracellular factor (EF) 3-hydroxypalmitic acid methyl ester biosynthesis protein | 4.77 | 0.029 | 0.75 | control | K19620 | GC_00027366 | 0.0 | 0.2 | Kitasatospora |
| adenosylcobinamide hydrolase [EC:3.5.1.90] | 4.77 | 0.029 | 0.75 | control | K08260 | GC_00028111 | 0.0 | 0.2 | Kitasatospora |
| phenylpyruvate C(3)-methyltransferase [EC:2.1.1.281] | 4.77 | 0.029 | 0.75 | control | K21457 | GC_00026745 | 0.0 | 0.2 | Kitasatospora |
| catechol O-methyltransferase [EC:2.1.1.6] | 4.77 | 0.029 | 0.75 | control | K00545 | GC_00030359 | 0.0 | 0.2 | Kitasatospora |
| (+)-beta-caryophyllene/(+)-caryolan-1-ol synthase [EC:4.2.3.89 4.2.1.138] | 4.77 | 0.029 | 0.75 | control | K18111 | GC_00025119 | 0.0 | 0.2 | Kitasatospora |
| ureidoacrylate peracid hydrolase [EC:3.5.1.110] | 4.77 | 0.029 | 0.75 | control | K09020 | GC_00028512 | 0.0 | 0.2 | Kitasatospora |
| cytochrome P450 family 20 subfamily A [EC:1.14.-.-] | 4.77 | 0.029 | 0.75 | control | K07435 | GC_00029795 | 0.0 | 0.2 | Kitasatospora |
| Xaa-Arg dipeptidase [EC:3.4.13.4] | 4.77 | 0.029 | 0.75 | control | K26141 | GC_00030282 | 0.0 | 0.2 | Kitasatospora |
| flagellin | 4.77 | 0.029 | 0.75 | control | K02406 | GC_00028736 | 0.0 | 0.2 | Kitasatospora |
| taurine dioxygenase [EC:1.14.11.17] | 4.77 | 0.029 | 0.75 | control | K03119 | GC_00027188 | 0.0 | 0.2 | Kitasatospora |
| GTP pyrophosphokinase [EC:2.7.6.5] | 4.77 | 0.029 | 0.75 | control | K07816 | GC_00025630 | 0.0 | 0.2 | Kitasatospora |
| Delta3-Delta2-enoyl-CoA isomerase [EC:5.3.3.8] | 4.77 | 0.029 | 0.75 | control | K07517 | GC_00027945 | 0.0 | 0.2 | Kitasatospora |
| AraC family transcriptional regulator, positive regulator of tynA and feaB | 4.77 | 0.029 | 0.75 | heated | K14063 | GC_00002222, GC_00003471, GC_00004334, GC_00005247, GC_00006834, GC_00011801, GC_00014302, GC_00016293, GC_00016448, GC_00016676, GC_00017172, GC_00027737, GC_00028606, GC_00030372 | 1.0 | 0.8 | Kitasatospora |

#### Table S4. KOfam Functional Enrichment

| Description | Enrichment Score | P-value | Q-value | Associated Treatment | KO Identifier | Gene Cluster IDs | p_Heated | p_Control | Clade |
|---|---|---|---|---|---|---|---|---|---|
| isobutyryl-CoA mutase small subunit [EC:5.4.99.13] | 4.77 | 0.029 | 0.75 | heated | K25821 | GC_00002532 | 1.0 | 0.8 | Kitasatospora |
| viologen exporter family transport system ATP-binding protein | 4.77 | 0.029 | 0.75 | heated | K25156 | GC_00002381, GC_00028986 | 1.0 | 0.8 | Kitasatospora |
| D-methionine transport system permease protein | 4.77 | 0.029 | 0.75 | heated | K02072 | GC_00002546 | 1.0 | 0.8 | Kitasatospora |
| viologen exporter family transport system permease protein | 4.77 | 0.029 | 0.75 | heated | K25155 | GC_00002582, GC_00002599, GC_00024439, GC_00025307 | 1.0 | 0.8 | Kitasatospora |
| ADP-dependent NAD(P)H-hydrate dehydratase / NAD(P)H-hydrate epimerase [EC:4.2.1.136 5.1.99.6] | 4.77 | 0.029 | 0.75 | heated | K23997 | GC_00002700 | 1.0 | 0.8 | Kitasatospora |
| 16S rRNA (adenine1518-N6/adenine1519-N6)-dimethyltransferase [EC:2.1.1.182] | 4.77 | 0.029 | 0.75 | heated | K02528 | GC_00002593 | 1.0 | 0.8 | Kitasatospora |
| glutathione transport system ATP-binding protein | 4.77 | 0.029 | 0.75 | heated | K13892 | GC_00000525 | 1.0 | 0.8 | Kitasatospora |
| preprotein translocase subunit SecE | 4.77 | 0.029 | 0.75 | heated | K03073 | GC_00002541, GC_00006353 | 1.0 | 0.8 | Kitasatospora |
| molybdopterin adenylyltransferase [EC:2.7.7.75] | 4.77 | 0.029 | 0.75 | heated | K03831 | GC_00002648 | 1.0 | 0.8 | Kitasatospora |
| L-lactate dehydrogenase complex protein LldE | 4.77 | 0.029 | 0.75 | heated | K18928 | GC_00000271 | 1.0 | 0.8 | Kitasatospora |
| L-lactate dehydrogenase complex protein LldG | 4.77 | 0.029 | 0.75 | heated | K00782 | GC_00003526, GC_00005866, GC_00015007 | 1.0 | 0.8 | Kitasatospora |
| aerobic C4-dicarboxylate transport protein | 4.77 | 0.029 | 0.75 | heated | K11103 | GC_00002438 | 1.0 | 0.8 | Kitasatospora |
| arsenate reductase (glutaredoxin) [EC:1.20.4.1] | 4.77 | 0.029 | 0.75 | heated | K00537 | GC_00002692 | 1.0 | 0.8 | Kitasatospora |
| endoglycosylceramidase [EC:3.2.1.123] | 4.77 | 0.029 | 0.75 | heated | K05991 | GC_00003040, GC_00016243 | 1.0 | 0.8 | Kitasatospora |
| copper transport protein | 4.77 | 0.029 | 0.75 | heated | K14166 | GC_00000505, GC_00011661 | 1.0 | 0.8 | Kitasatospora |
| ribulose-phosphate 3-epimerase [EC:5.1.3.1] | 4.77 | 0.029 | 0.75 | heated | K01783 | GC_00002559, GC_00030699 | 1.0 | 0.8 | Kitasatospora |
| cell division protein FtsL | 4.77 | 0.029 | 0.75 | heated | K03586 | GC_00002712, GC_00003115, GC_00018835 | 1.0 | 0.8 | Kitasatospora |
| ATP synthase protein I | 4.77 | 0.029 | 0.75 | heated | K02116 | GC_00002555, GC_00019837 | 1.0 | 0.8 | Kitasatospora |
| O-acetylserine/cysteine efflux transporter | 4.77 | 0.029 | 0.75 | heated | K15268 | GC_00002719, GC_00003350, GC_00003980, GC_00027552 | 1.0 | 0.8 | Kitasatospora |
| branched-chain amino acid transport system substrate-binding protein | 4.77 | 0.029 | 0.75 | heated | K01999 | GC_00002608, GC_00011727, GC_00012062, GC_00012931, GC_00022008 | 1.0 | 0.8 | Kitasatospora |
| TetR/AcrR family transcriptional regulator, regulator of biofilm formation and stress response | 4.77 | 0.029 | 0.75 | heated | K23778 | GC_00004758, GC_00004805, GC_00005278, GC_00005352, GC_00005852, GC_00006167, GC_00014423, GC_00017419 | 1.0 | 0.8 | Kitasatospora |
| thiamine-phosphate pyrophosphorylase [EC:2.5.1.3] | 4.77 | 0.029 | 0.75 | heated | K00788 | GC_00002677 | 1.0 | 0.8 | Kitasatospora |
| endoribonuclease LACTB2 [EC:3.1.27.-] | 4.77 | 0.029 | 0.75 | heated | K16639 | GC_00001308 | 1.0 | 0.8 | Kitasatospora |
| NAD+ diphosphatase [EC:3.6.1.22] | 4.77 | 0.029 | 0.75 | heated | K03426 | GC_00002612, GC_00016870 | 1.0 | 0.8 | Kitasatospora |
| heme oxygenase (mycobilin-producing) [EC:1.14.99.57] | 4.77 | 0.029 | 0.75 | heated | K21481 | GC_00000560 | 1.0 | 0.8 | Kitasatospora |
| starch synthase (maltosyl-transferring) [EC:2.4.99.16] | 4.77 | 0.029 | 0.75 | heated | K16147 | GC_00000219 | 1.0 | 0.8 | Kitasatospora |
| O-acetyl-ADP-ribose deacetylase [EC:3.1.1.106] | 4.77 | 0.029 | 0.75 | heated | K23518 | GC_00002654 | 1.0 | 0.8 | Kitasatospora |
| nitrite reductase (NADH) large subunit [EC:1.7.1.15] | 4.57 | 0.033 | 0.78 | heated | K00362 | GC_00004972 | 0.5 | 0.0 | Kitasatospora |
| lanthionine-containing peptide SapB | 4.57 | 0.033 | 0.78 | heated | K24913 | GC_00005053 | 0.5 | 0.0 | Kitasatospora |
| uroporphyrinogen-III synthase [EC:4.2.1.75] | 4.57 | 0.033 | 0.78 | heated | K01719 | GC_00005173, GC_00012627 | 0.5 | 0.0 | Kitasatospora |
| calicheamicin 3'-O-methyl-rhamnosyltransferase [EC:2.4.1.-] | 4.57 | 0.033 | 0.78 | heated | K21263 | GC_00005620, GC_00009704 | 0.5 | 0.0 | Kitasatospora |
| NDP-mannose synthase | 4.57 | 0.033 | 0.78 | control | K21210 | GC_00004251 | 0.5 | 1.0 | Kitasatospora |
| NDP-hexose 4,6-dehydratase | 4.57 | 0.033 | 0.78 | control | K21211 | GC_00004270 | 0.5 | 1.0 | Kitasatospora |
| ketoreductase [EC:1.1.1.-] | 4.57 | 0.033 | 0.78 | control | K12420 | GC_00005945, GC_00009532, GC_00021125 | 0.5 | 1.0 | Kitasatospora |

#### Table S4. KOfam Functional Enrichment

| Description | Enrichment Score | P-value | Q-value | Associated Treatment | KO Identifier | Gene Cluster IDs | p_Heated | p_Control | Clade |
|---|---|---|---|---|---|---|---|---|---|
| NitT/TauT family transport system permease protein | 4.57 | 0.033 | 0.78 | control | K02050 | GC_00004322, GC_00011643 | 0.5 | 1.0 | Kitasatospora |
| acetyl-CoA carboxylase, biotin carboxylase subunit [EC:6.4.1.2 6.3.4.14] | 4.23 | 0.040 | 0.92 | control | K01961 | GC_00005507 | 0.3 | 0.8 | Kitasatospora |
| spermidine/putrescine transport system permease protein | 4.23 | 0.040 | 0.92 | control | K11071 | GC_00005431, GC_00005532 | 0.3 | 0.8 | Kitasatospora |
| spermidine/putrescine transport system ATP-binding protein [EC:7.6.2.11] | 4.23 | 0.040 | 0.92 | control | K11072 | GC_00005558 | 0.3 | 0.8 | Kitasatospora |
| 4,5-DOPA dioxygenase extradiol [EC:1.13.11.-] | 4.23 | 0.040 | 0.92 | control | K15777 | GC_00010082, GC_00011523 | 0.3 | 0.8 | Kitasatospora |
| D-amino-acid oxidase [EC:1.4.3.3] | 3.98 | 0.046 | 0.95 | control | K00273 | GC_00008207 | 0.2 | 0.6 | Kitasatospora |
| lantibiotic bacteriocin | 3.98 | 0.046 | 0.95 | control | K20482 | GC_00006294 | 0.2 | 0.6 | Kitasatospora |
| adenylate cyclase [EC:4.6.1.1] | 3.98 | 0.046 | 0.95 | control | K01768 | GC_00011602, GC_00016893, GC_00023100 | 0.2 | 0.6 | Kitasatospora |
| esterase FrsA [EC:3.1.-.-] | 3.98 | 0.046 | 0.95 | control | K11750 | GC_00010997, GC_00020805 | 0.2 | 0.6 | Kitasatospora |
| aminocarboxymuconate-semialdehyde decarboxylase [EC:4.1.1.45] | 3.98 | 0.046 | 0.95 | heated | K03392 | GC_00004626, GC_00008818 | 0.8 | 0.4 | Kitasatospora |
| choline dehydrogenase [EC:1.1.99.1] | 3.98 | 0.046 | 0.95 | heated | K00108 | GC_00003646, GC_00029120 | 0.8 | 0.4 | Kitasatospora |
| 2-phosphosulfolactate phosphatase [EC:3.1.3.71] | 3.94 | 0.047 | 0.95 | heated | K05979 | GC_00005869, GC_00008057 | 0.5 | 0.0 | Kitasatospora |
| assimilatory nitrate reductase catalytic subunit [EC:1.7.99.-] | 3.94 | 0.047 | 0.95 | heated | K00372 | GC_00005201 | 0.5 | 0.0 | Kitasatospora |
| assimilatory nitrate reductase electron transfer subunit [EC:1.7.99. | 3.94 | 0.047 | 0.95 | heated | K00360 | GC_00005371 | 0.5 | 0.0 | Kitasatospora |
| nitrite reductase (NADH) small subunit [EC:1.7.1.15] | 3.94 | 0.047 | 0.95 | heated | K00363 | GC_00005380 | 0.5 | 0.0 | Kitasatospora |
| CRISPR system Cascade subunit CasD | 3.94 | 0.047 | 0.95 | control | K19125 | GC_00006893, GC_00007159, GC_00008758, GC_00017209 | 0.5 | 1.0 | Kitasatospora |
| superoxide dismutase, Fe-Mn family [EC:1.15.1.1] | 3.94 | 0.047 | 0.95 | control | K04564 | GC_00004131 | 0.5 | 1.0 | Kitasatospora |
| erythromycin esterase [EC:3.1.1.-] | 3.94 | 0.047 | 0.95 | control | K06880 | GC_00004004, GC_00012095, GC_00016333, GC_00028112, GC_00029602 | 0.5 | 1.0 | Kitasatospora |
| iron uptake system component EfeO | 3.94 | 0.047 | 0.95 | control | K07224 | GC_00004752, GC_00016137 | 0.5 | 1.0 | Kitasatospora |
| adenylate cyclase, class 2 [EC:4.6.1.1] | 3.94 | 0.047 | 0.95 | control | K05873 | GC_00003152, GC_00030091 | 0.5 | 1.0 | Kitasatospora |
| N-acyl-D-amino-acid deacylase [EC:3.5.1.81] | 3.94 | 0.047 | 0.95 | control | K06015 | GC_00003995 | 0.5 | 1.0 | Kitasatospora |
| mercuric ion transport protein | 7.50 | 0.006 | 1.00 | control | K19058 | GC_00004249, GC_00004888 | 0.4 | 1.0 | Bradyrhizobium |
| MerR family transcriptional regulator, mercuric resistance operon regulatory protein | 6.56 | 0.010 | 1.00 | control | K08365 | GC_00005825, GC_00017188, GC_00020937 | 0.0 | 0.7 | Bradyrhizobium |
| aldose sugar dehydrogenase [EC:1.1.5.-] | 5.00 | 0.025 | 1.00 | control | K21430 | GC_00004609 | 0.2 | 0.8 | Bradyrhizobium |
| flavin prenyltransferase [EC:2.5.1.129] | 5.00 | 0.025 | 1.00 | heated | K03186 | GC_00006040, GC_00009484, GC_00010750 | 0.8 | 0.2 | Bradyrhizobium |
| N-acetylglucosaminyl-diphospho-decaprenol L-rhamnosyltransferase [EC:2.4.1.289] | 5.00 | 0.025 | 1.00 | control | K16870 | GC_00005966, GC_00019054, GC_00024041, GC_00024221 | 0.0 | 0.6 | Bradyrhizobium |
| oleate hydratase [EC:4.2.1.53] | 4.62 | 0.032 | 1.00 | heated | K10254 | GC_00014562, GC_00024883 | 0.4 | 0.0 | Bradyrhizobium |
| glucosyl-3-phosphoglycerate synthase [EC:2.4.1.266] | 4.62 | 0.032 | 1.00 | heated | K13693 | GC_00012003 | 0.4 | 0.0 | Bradyrhizobium |
| decaprenyl-phosphate phosphoribosyltransferase [EC:2.4.2.45] | 4.62 | 0.032 | 1.00 | heated | K14136 | GC_00016509, GC_00018289 | 0.4 | 0.0 | Bradyrhizobium |
| carboxylesterase [EC:3.1.1.1] | 4.62 | 0.032 | 1.00 | heated | K03928 | GC_00011598 | 0.4 | 0.0 | Bradyrhizobium |
| alcohol dehydrogenase [EC:1.1.1.-] | 4.62 | 0.032 | 1.00 | heated | K18369 | GC_00020023, GC_00022054 | 0.4 | 0.0 | Bradyrhizobium |
| ribonucleoside-diphosphate reductase beta chain [EC:1.17.4.1] | 4.62 | 0.032 | 1.00 | heated | K00526 | GC_00013345 | 0.4 | 0.0 | Bradyrhizobium |
| 4-aminobutyrate---pyruvate transaminase [EC:2.6.1.96] | 4.62 | 0.032 | 1.00 | heated | K16871 | GC_00012424 | 0.4 | 0.0 | Bradyrhizobium |
| (R,R)-butanediol dehydrogenase / meso-butanediol dehydrogenase / diacetyl reductase [EC:1.1.1.4 1.1.1.- 1.1.1.303] | 4.62 | 0.032 | 1.00 | heated | K00004 | GC_00019124, GC_00022146 | 0.4 | 0.0 | Bradyrhizobium |

#### Table S4. KOfam Functional Enrichment

| Description | Enrichment Score | P-value | Q-value | Associated Treatment | KO Identifier | Gene Cluster IDs | p_Heated | p_Control | Clade |
|---|---|---|---|---|---|---|---|---|---|
| peptidoglycan LD-endopeptidase CwlK [EC:3.4.-.-] | 4.62 | 0.032 | 1.00 | heated | K17733 | GC_00012620, GC_00020628 | 0.4 | 0.0 | Bradyrhizobium |
| vitamin B12 transporter | 4.62 | 0.032 | 1.00 | heated | K16092 | GC_00011777 | 0.4 | 0.0 | Bradyrhizobium |
| pyrimidine oxygenase [EC:1.14.99.46] | 4.62 | 0.032 | 1.00 | heated | K09018 | GC_00018396, GC_00023840 | 0.4 | 0.0 | Bradyrhizobium |
| succinyl-CoA:acetate CoA-transferase [EC:2.8.3.18] | 4.62 | 0.032 | 1.00 | heated | K18118 | GC_00012404 | 0.4 | 0.0 | Bradyrhizobium |
| MFS transporter, Spinster family, sphingosine-1-phosphate transporter | 4.62 | 0.032 | 1.00 | control | K23677 | GC_00003539, GC_00009082, GC_00009895 | 0.6 | 1.0 | Bradyrhizobium |
| phytanoyl-CoA hydroxylase [EC:1.14.11.18] | 4.62 | 0.032 | 1.00 | control | K00477 | GC_00003798, GC_00004803, GC_00020322, GC_00023954 | 0.6 | 1.0 | Bradyrhizobium |
| spermidine synthase [EC:2.5.1.16] | 4.62 | 0.032 | 1.00 | control | K00797 | GC_00004761, GC_00004893, GC_00015455, GC_00020520 | 0.6 | 1.0 | Bradyrhizobium |
| cell filamentation protein, protein adenylyltransferase [EC:2.7.7.10 | 4.62 | 0.032 | 1.00 | control | K04095 | GC_00006262, GC_00010315, GC_00010399, GC_00010634, GC_00012206, GC_00013808, GC_00015965, GC_00021162, GC_00021214, GC_00022608, GC_00026007 | 0.6 | 1.0 | Bradyrhizobium |
| pimeloyl-[acyl-carrier protein] methyl ester esterase [EC:3.1.1.85] | 4.62 | 0.032 | 1.00 | control | K02170 | GC_00003531 | 0.6 | 1.0 | Bradyrhizobium |
| bifunctional enzyme CysN/CysC [EC:2.7.7.4 2.7.1.25] | 4.26 | 0.039 | 1.00 | heated | K00955 | GC_00007761 | 0.6 | 0.1 | Bradyrhizobium |
| diphthine-ammonia ligase [EC:6.3.1.14] | 4.26 | 0.039 | 1.00 | heated | K06927 | GC_00007675 | 0.6 | 0.1 | Bradyrhizobium |
| porin | 4.26 | 0.039 | 1.00 | heated | K07267 | GC_00012483, GC_00013359, GC_00015225, GC_00017427, GC_00017871, GC_00021685 | 0.6 | 0.1 | Bradyrhizobium |
| ribosome-dependent ATPase | 4.26 | 0.039 | 1.00 | heated | K13926 | GC_00006975 | 0.6 | 0.1 | Bradyrhizobium |
| restriction system protein | 4.26 | 0.039 | 1.00 | heated | K07448 | GC_00013073, GC_00014310, GC_00020243, GC_00024998 | 0.6 | 0.1 | Bradyrhizobium |
| cobalamin biosynthesis protein CobC | 4.26 | 0.039 | 1.00 | control | K02225 | GC_00006016, GC_00009971, GC_00012944 | 0.4 | 0.9 | Bradyrhizobium |
| alpha-terpineol hydroxylase | 4.26 | 0.039 | 1.00 | control | K24391 | GC_00004166 | 0.4 | 0.9 | Bradyrhizobium |
| antitoxin Phd | 7.78 | 0.005 | 0.37 | control | K19165 | GC_00005733, GC_00006702, GC_00006805 | 0.0 | 0.8 | Rhizobium |
| para-nitrobenzyl esterase [EC:3.1.1.-] | 7.78 | 0.005 | 0.37 | control | K03929 | GC_00004995, GC_00005404, GC_00005681 | 0.0 | 0.8 | Rhizobium |
| FAD-dependent urate hydroxylase [EC:1.14.13.113] | 7.78 | 0.005 | 0.37 | control | K22879 | GC_00004951 | 0.0 | 0.8 | Rhizobium |
| LysR family transcriptional regulator, benzoate and cis,cis-muconate-responsive activator of ben and cat genes | 7.78 | 0.005 | 0.37 | control | K21757 | GC_00005208, GC_00006886 | 0.0 | 0.8 | Rhizobium |
| cell filamentation protein, protein adenylyltransferase [EC:2.7.7.10 | 7.78 | 0.005 | 0.37 | control | K04095 | GC_00005084, GC_00005477, GC_00006353, GC_00006468, GC_00006890, GC_00007645 | 0.0 | 0.8 | Rhizobium |
| alanine dehydrogenase [EC:1.4.1.1] | 7.78 | 0.005 | 0.37 | control | K19244 | GC_00005227, GC_00006715 | 0.0 | 0.8 | Rhizobium |
| gamma-glutamylcyclotransferase [EC:4.3.2.9] | 7.78 | 0.005 | 0.37 | heated | K00682 | GC_00004500 | 1.0 | 0.2 | Rhizobium |
| RpiR family transcriptional regulator, repressor of rpiB and als oper | 7.78 | 0.005 | 0.37 | heated | K23238 | GC_00004616 | 1.0 | 0.2 | Rhizobium |
| TetR/AcrR family transcriptional regulator, repressor of the mexAB-oprM multidrug resistance operon | 7.78 | 0.005 | 0.37 | heated | K18135 | GC_00004719 | 1.0 | 0.2 | Rhizobium |
| alcohol dehydrogenase [EC:1.1.1.-] | 7.78 | 0.005 | 0.37 | heated | K18369 | GC_00004556 | 1.0 | 0.2 | Rhizobium |
| cobalt-zinc-cadmium efflux system protein | 7.78 | 0.005 | 0.37 | heated | K16264 | GC_00004825 | 1.0 | 0.2 | Rhizobium |
| primary-amine oxidase [EC:1.4.3.21] | 7.78 | 0.005 | 0.37 | heated | K00276 | GC_00004899 | 1.0 | 0.2 | Rhizobium |
| tellurite methyltransferase [EC:2.1.1.265] | 7.78 | 0.005 | 0.37 | heated | K16868 | GC_00004640 | 1.0 | 0.2 | Rhizobium |
| DNA sulfur modification protein DndE | 7.78 | 0.005 | 0.37 | heated | K19172 | GC_00004722 | 1.0 | 0.2 | Rhizobium |
| MFS transporter, DHA1 family, multidrug resistance protein B | 7.78 | 0.005 | 0.37 | heated | K08152 | GC_00002364 | 1.0 | 0.2 | Rhizobium |

#### Table S4. KOfam Functional Enrichment

| Description | Enrichment Score | P-value | Q-value | Associated Treatment | KO Identifier | Gene Cluster IDs | p_Heated | p_Control | Clade |
|---|---|---|---|---|---|---|---|---|---|
| 3-hydroxy-9,10-secoandrosta-1,3,5(10)-triene-9,17-dione monooxygenase [EC:1.14.14.12] | 7.78 | 0.005 | 0.37 | heated | K16047 | GC_00001250 | 1.0 | 0.2 | Rhizobium |
| glucose dehydrogenase [EC:1.1.5.9] | 7.78 | 0.005 | 0.37 | heated | K19813 | GC_00004917 | 1.0 | 0.2 | Rhizobium |
| DNA sulfur modification protein DndD | 7.78 | 0.005 | 0.37 | heated | K19171 | GC_00003866 | 1.0 | 0.2 | Rhizobium |
| dimethylamine monooxygenase subunit B [EC:1.14.13.238] | 7.78 | 0.005 | 0.37 | heated | K22343 | GC_00004797 | 1.0 | 0.2 | Rhizobium |
| trimethylamine monooxygenase [EC:1.14.13.148] | 7.78 | 0.005 | 0.37 | heated | K18277 | GC_00004889 | 1.0 | 0.2 | Rhizobium |
| DNA sulfur modification protein DndB | 7.78 | 0.005 | 0.37 | heated | K19169 | GC_00004337, GC_00004549 | 1.0 | 0.2 | Rhizobium |
| limonene 1,2-monooxygenase [EC:1.14.13.107] | 7.78 | 0.005 | 0.37 | heated | K14733 | GC_00004418 | 1.0 | 0.2 | Rhizobium |
| dimethylamine monooxygenase subunit C [EC:1.14.13.238] | 7.78 | 0.005 | 0.37 | heated | K22344 | GC_00004781 | 1.0 | 0.2 | Rhizobium |
| N,N-dimethylformamidase large subunit [EC:3.5.1.56] | 7.78 | 0.005 | 0.37 | heated | K03418 | GC_00004675 | 1.0 | 0.2 | Rhizobium |
| RNA-directed DNA polymerase [EC:2.7.7.49] | 7.78 | 0.005 | 0.37 | heated | K00986 | GC_00004998 | 1.0 | 0.2 | Rhizobium |
| fructose 5-dehydrogenase cytochrome subunit | 7.78 | 0.005 | 0.37 | heated | K23275 | GC_00004878 | 1.0 | 0.2 | Rhizobium |
| DNA sulfur modification protein DndC | 7.78 | 0.005 | 0.37 | heated | K19170 | GC_00003922 | 1.0 | 0.2 | Rhizobium |
| 4,5-dihydroxyphthalate decarboxylase [EC:4.1.1.55] | 7.78 | 0.005 | 0.37 | heated | K04102 | GC_00003733 | 1.0 | 0.2 | Rhizobium |
| dimethylamine monooxygenase subunit A [EC:1.14.13.238] | 7.78 | 0.005 | 0.37 | heated | K22342 | GC_00004841 | 1.0 | 0.2 | Rhizobium |
| protein-tyrosine phosphatase [EC:3.1.3.48] | 7.78 | 0.005 | 0.37 | heated | K01104 | GC_00004710 | 1.0 | 0.2 | Rhizobium |
| fructose 5-dehydrogenase small subunit [EC:1.1.5.14] | 7.78 | 0.005 | 0.37 | heated | K23274 | GC_00004906 | 1.0 | 0.2 | Rhizobium |
| DNA-binding protein HU-beta | 6.64 | 0.010 | 0.68 | control | K03530 | GC_00004426, GC_00005335 | 0.2 | 0.9 | Rhizobium |
| Rrf2 family transcriptional regulator, nitric oxide-sensitive transcriptional repressor | 5.83 | 0.016 | 0.77 | heated | K13771 | GC_00004586 | 1.0 | 0.3 | Rhizobium |
| aspartate 1-decarboxylase [EC:4.1.1.11] | 5.83 | 0.016 | 0.77 | heated | K01579 | GC_00004562 | 1.0 | 0.3 | Rhizobium |
| 5-methylcytosine-specific restriction enzyme A [EC:3.1.21.-] | 5.83 | 0.016 | 0.77 | heated | K07451 | GC_00003918, GC_00007265, GC_00007760 | 1.0 | 0.3 | Rhizobium |
| peptidyl carrier protein | 5.83 | 0.016 | 0.77 | heated | K21183 | GC_00004498 | 1.0 | 0.3 | Rhizobium |
| DNA mismatch endonuclease, patch repair protein [EC:3.1.-.-] | 5.83 | 0.016 | 0.77 | heated | K07458 | GC_00004580 | 1.0 | 0.3 | Rhizobium |
| MbtH protein | 5.83 | 0.016 | 0.77 | heated | K05375 | GC_00004590 | 1.0 | 0.3 | Rhizobium |
| D-glycero-D-manno-heptose 1,7-bisphosphate phosphatase [EC:3.1.3.82 3.1.3.83] | 5.83 | 0.016 | 0.77 | control | K03273 | GC_00005376, GC_00006750 | 0.0 | 0.7 | Rhizobium |
| Bacteriophage probable baseplate hub protein | 5.83 | 0.016 | 0.77 | control | K06905 | GC_00005334, GC_00006740 | 0.0 | 0.7 | Rhizobium |
| mRNA interferase HigB [EC:3.1.-.-] | 4.38 | 0.036 | 0.77 | heated | K19166 | GC_00005049 | 0.8 | 0.2 | Rhizobium |
| cyclohexyl-isocyanide hydratase [EC:4.2.1.103] | 4.32 | 0.038 | 0.77 | heated | K18199 | GC_00004551 | 1.0 | 0.4 | Rhizobium |
| FAD:protein FMN transferase [EC:2.7.1.180] | 4.32 | 0.038 | 0.77 | heated | K03734 | GC_00004153 | 1.0 | 0.4 | Rhizobium |
| alginate O-acetyltransferase complex protein AlgI | 4.32 | 0.038 | 0.77 | heated | K19294 | GC_00004548 | 1.0 | 0.4 | Rhizobium |
| filamentous hemagglutinin | 4.32 | 0.038 | 0.77 | heated | K15125 | GC_00000253 | 1.0 | 0.4 | Rhizobium |
| limonene-1,2-epoxide hydrolase [EC:3.3.2.8] | 4.32 | 0.038 | 0.77 | heated | K10533 | GC_00004522 | 1.0 | 0.4 | Rhizobium |
| LuxR family transcriptional regulator, activator of conjugal transfer of Ti plasmids | 4.32 | 0.038 | 0.77 | heated | K19732 | GC_00004631, GC_00006943 | 1.0 | 0.4 | Rhizobium |
| mannuronan 5-epimerase [EC:5.1.3.37] | 4.32 | 0.038 | 0.77 | heated | K01795 | GC_00000274 | 1.0 | 0.4 | Rhizobium |
| 5-(hydroxymethyl)furfural/furfural oxidase [EC:1.1.3.47 1.1.3.-] | 4.32 | 0.038 | 0.77 | heated | K16873 | GC_00004392 | 1.0 | 0.4 | Rhizobium |
| GDP-mannose 6-dehydrogenase [EC:1.1.1.132] | 4.32 | 0.038 | 0.77 | heated | K00066 | GC_00004521 | 1.0 | 0.4 | Rhizobium |
| death on curing protein | 4.32 | 0.038 | 0.77 | heated | K07341 | GC_00004476 | 1.0 | 0.4 | Rhizobium |

#### Table S4. KOfam Functional Enrichment

| Description | Enrichment Score | P-value | Q-value | Associated Treatment | KO Identifier | Gene Cluster IDs | p_Heated | p_Control | Clade |
|---|---|---|---|---|---|---|---|---|---|
| TraR antiactivator | 4.32 | 0.038 | 0.77 | heated | K20272 | GC_00004748, GC_00006213 | 1.0 | 0.4 | Rhizobium |
| competence protein ComFC | 4.32 | 0.038 | 0.77 | heated | K02242 | GC_00004545 | 1.0 | 0.4 | Rhizobium |
| acyl homoserine lactone synthase [EC:2.3.1.184] | 4.32 | 0.038 | 0.77 | heated | K22955 | GC_00004249 | 1.0 | 0.4 | Rhizobium |
| inhibitor of KinA | 4.32 | 0.038 | 0.77 | heated | K06351 | GC_00004683, GC_00006346 | 1.0 | 0.4 | Rhizobium |
| type IV secretion system protein TrbH | 4.32 | 0.038 | 0.77 | heated | K20267 | GC_00004449 | 1.0 | 0.4 | Rhizobium |
| mannuronan synthase [EC:2.4.1.33] | 4.32 | 0.038 | 0.77 | heated | K19290 | GC_00003953, GC_00004326 | 1.0 | 0.4 | Rhizobium |
| alginate O-acetyltransferase complex protein AlgJ | 4.32 | 0.038 | 0.77 | heated | K19295 | GC_00004465 | 1.0 | 0.4 | Rhizobium |
| type IV secretion system protein TrbG | 4.32 | 0.038 | 0.77 | heated | K20532 | GC_00004401 | 1.0 | 0.4 | Rhizobium |
| periplasmic protein CpxP/Spy | 4.32 | 0.038 | 0.77 | heated | K06006 | GC_00004380 | 1.0 | 0.4 | Rhizobium |
| DNA (cytosine-5)-methyltransferase 1 [EC:2.1.1.37] | 4.32 | 0.038 | 0.77 | heated | K00558 | GC_00004599, GC_00004666, GC_00004941, GC_00004961, GC_00007065, GC_00007853 | 1.0 | 0.4 | Rhizobium |
| 5'-nucleotidase [EC:3.1.3.5] | 4.32 | 0.038 | 0.77 | heated | K02566 | GC_00000965 | 1.0 | 0.4 | Rhizobium |
| type IV secretion system protein TrbC | 4.32 | 0.038 | 0.77 | heated | K20528 | GC_00004421, GC_00007376 | 1.0 | 0.4 | Rhizobium |
| hemolysin activation/secretion protein | 4.32 | 0.038 | 0.77 | heated | K07326 | GC_00003970 | 1.0 | 0.4 | Rhizobium |
| large repetitive protein | 4.32 | 0.038 | 0.77 | heated | K20276 | GC_00004422, GC_00006445 | 1.0 | 0.4 | Rhizobium |
| 2,5-diketo-D-gluconate reductase A [EC:1.1.1.346] | 4.32 | 0.038 | 0.77 | heated | K06221 | GC_00004475 | 1.0 | 0.4 | Rhizobium |
| TetR/AcrR family transcriptional regulator, cholesterol catabolism regulator | 4.32 | 0.038 | 0.77 | heated | K22107 | GC_00004524 | 1.0 | 0.4 | Rhizobium |
| alginate O-acetyltransferase complex protein AlgF | 4.32 | 0.038 | 0.77 | heated | K19296 | GC_00004373 | 1.0 | 0.4 | Rhizobium |
| alginate biosynthesis protein AlgX | 4.32 | 0.038 | 0.77 | heated | K19293 | GC_00004437 | 1.0 | 0.4 | Rhizobium |
| DNA-damage-inducible protein J | 4.32 | 0.038 | 0.77 | heated | K07473 | GC_00000546 | 1.0 | 0.4 | Rhizobium |
| type IV secretion system protein TrbK | 4.32 | 0.038 | 0.77 | heated | K20555 | GC_00004756, GC_00006167 | 1.0 | 0.4 | Rhizobium |
| aspartate-semialdehyde dehydrogenase [EC:1.2.1.-] | 4.32 | 0.038 | 0.77 | control | K15786 | GC_00005174 | 0.0 | 0.6 | Rhizobium |
| tRNA (cmo5U34)-methyltransferase [EC:2.1.1.-] | 4.32 | 0.038 | 0.77 | control | K15256 | GC_00005096, GC_00005337 | 0.0 | 0.6 | Rhizobium |
| type IV secretion system protein VirB4 [EC:7.4.2.8] | 4.32 | 0.038 | 0.77 | control | K03199 | GC_00005112 | 0.0 | 0.6 | Rhizobium |
| arylformamidase [EC:3.5.1.9] | 4.32 | 0.038 | 0.77 | control | K07130 | GC_00005162, GC_00005447, GC_00006745 | 0.0 | 0.6 | Rhizobium |
| type IV secretion system protein VirB9 | 4.32 | 0.038 | 0.77 | control | K03204 | GC_00005255 | 0.0 | 0.6 | Rhizobium |
| type IV secretion system protein VirB7 | 4.32 | 0.038 | 0.77 | control | K03202 | GC_00005142 | 0.0 | 0.6 | Rhizobium |
| 2-dehydro-3-deoxy-L-rhamnonate dehydrogenase (NAD+) [EC:1.1.1.401] | 4.32 | 0.038 | 0.77 | control | K21883 | GC_00005163, GC_00006435 | 0.0 | 0.6 | Rhizobium |
| ectoine hydrolase [EC:3.5.4.44] | 4.32 | 0.038 | 0.77 | control | K15783 | GC_00005157 | 0.0 | 0.6 | Rhizobium |
| mannopine transport system substrate-binding protein | 4.32 | 0.038 | 0.77 | control | K11077 | GC_00005262 | 0.0 | 0.6 | Rhizobium |
| type IV secretion system protein VirB3 | 4.32 | 0.038 | 0.77 | control | K03198 | GC_00005115 | 0.0 | 0.6 | Rhizobium |
| chromate reductase, NAD(P)H dehydrogenase (quinone) | 4.32 | 0.038 | 0.77 | control | K19784 | GC_00005218 | 0.0 | 0.6 | Rhizobium |
| aldehyde dehydrogenase (NAD+) [EC:1.2.1.3] | 4.32 | 0.038 | 0.77 | control | K00128 | GC_00004655 | 0.0 | 0.6 | Rhizobium |
| mannopine transport system permease protein | 4.32 | 0.038 | 0.77 | control | K11078 | GC_00005173, GC_00005176 | 0.0 | 0.6 | Rhizobium |
| type IV secretion system protein VirB10 | 4.32 | 0.038 | 0.77 | control | K03195 | GC_00005230 | 0.0 | 0.6 | Rhizobium |
| GntR family transcriptional regulator, colanic acid and biofilm gene transcriptional regulator | 4.32 | 0.038 | 0.77 | control | K13654 | GC_00005150 | 0.0 | 0.6 | Rhizobium |
| simple sugar transport system ATP-binding protein [EC:7.5.2.-] | 4.32 | 0.038 | 0.77 | control | K02056 | GC_00005144 | 0.0 | 0.6 | Rhizobium |

#### Table S4. KOfam Functional Enrichment

| Description | Enrichment Score | P-value | Q-value | Associated Treatment | KO Identifier | Gene Cluster IDs | p_Heated | p_Control | Clade |
|---|---|---|---|---|---|---|---|---|---|
| putative Mg2+ transporter-C (MgtC) family protein | 4.32 | 0.038 | 0.77 | control | K07507 | GC_00005152 | 0.0 | 0.6 | Rhizobium |
| type IV secretion system protein VirB6 | 4.32 | 0.038 | 0.77 | control | K03201 | GC_00005147 | 0.0 | 0.6 | Rhizobium |
| type IV secretion system protein VirB1 | 4.32 | 0.038 | 0.77 | control | K03194 | GC_00005128 | 0.0 | 0.6 | Rhizobium |
| IclR family transcriptional regulator, blcABC operon repressor | 4.32 | 0.038 | 0.77 | control | K20539 | GC_00005178 | 0.0 | 0.6 | Rhizobium |
| mannopine transport system ATP-binding protein | 4.32 | 0.038 | 0.77 | control | K11080 | GC_00005161 | 0.0 | 0.6 | Rhizobium |
| transposase, IS30 family | 4.32 | 0.038 | 0.77 | control | K07482 | GC_00005650, GC_00006268 | 0.0 | 0.6 | Rhizobium |
| type IV secretion system protein VirB11 [EC:7.4.2.8] | 4.32 | 0.038 | 0.77 | control | K03196 | GC_00005046 | 0.0 | 0.6 | Rhizobium |
| 8-hydroxy-5-deazaflavin:NADPH oxidoreductase [EC:1.5.1.40] | 4.32 | 0.038 | 0.77 | control | K06988 | GC_00005793, GC_00006394 | 0.0 | 0.6 | Rhizobium |
| N2-acetyl-L-2,4-diaminobutanoate deacetylase [EC:3.5.1.125] | 4.32 | 0.038 | 0.77 | control | K15784 | GC_00005195 | 0.0 | 0.6 | Rhizobium |
| type IV secretion system protein VirB2 | 4.32 | 0.038 | 0.77 | control | K03197 | GC_00005098 | 0.0 | 0.6 | Rhizobium |
| type IV secretion system protein VirB8 | 4.32 | 0.038 | 0.77 | control | K03203 | GC_00005103 | 0.0 | 0.6 | Rhizobium |
| antitoxin FitA | 4.32 | 0.038 | 0.77 | control | K21495 | GC_00005093 | 0.0 | 0.6 | Rhizobium |
| DeoR family transcriptional regulator, deoxyribose operon represso | 4.32 | 0.038 | 0.77 | control | K11534 | GC_00004672 | 0.0 | 0.6 | Rhizobium |
| type IV secretion system protein VirB5 | 4.32 | 0.038 | 0.77 | control | K03200 | GC_00005229 | 0.0 | 0.6 | Rhizobium |
| glutamate dehydrogenase (NADP+) [EC:1.4.1.4] | 4.32 | 0.038 | 0.77 | control | K00262 | GC_00005253 | 0.0 | 0.6 | Rhizobium |
| regulator of nucleoside diphosphate kinase | 4.32 | 0.038 | 0.77 | control | K06140 | GC_00005102, GC_00005119 | 0.0 | 0.6 | Rhizobium |
| 3-demethoxyubiquinol 3-hydroxylase [EC:1.14.99.60] | 4.32 | 0.038 | 0.77 | control | K06134 | GC_00005231 | 0.0 | 0.6 | Rhizobium |
| D-alanyl-D-alanine carboxypeptidase [EC:3.4.16.4] | 4.32 | 0.038 | 0.77 | control | K01286 | GC_00005584, GC_00006158 | 0.0 | 0.6 | Rhizobium |
| L-2,4-diaminobutyrate transaminase [EC:2.6.1.76] | 4.32 | 0.038 | 0.77 | control | K15785 | GC_00005082 | 0.0 | 0.6 | Rhizobium |
| SlyX protein | 4.20 | 0.040 | 0.80 | control | K03745 | GC_00003990 | 0.6 | 1.0 | Rhizobium |
| chorismate mutase [EC:5.4.99.5] | 4.20 | 0.040 | 0.80 | control | K04092 | GC_00004002, GC_00005864 | 0.6 | 1.0 | Rhizobium |
| large subunit ribosomal protein L27 | 4.20 | 0.040 | 0.80 | control | K02899 | GC_00003987 | 0.6 | 1.0 | Rhizobium |
| flagellar biosynthesis protein | 4.20 | 0.040 | 0.80 | heated | K04061 | GC_00006076 | 0.4 | 0.0 | Rhizobium |
| antitoxin HigA-1 | 8.12 | 0.004 | 1.00 | heated | K21498 | GC_00004018, GC_00004720, GC_00010539, GC_00010540, GC_00017723, GC_00022049, GC_00024334 | 1.0 | 0.4 | Paraburkholderia |
| 3-oxoisoapionate-4-phosphate transcarboxylase/hydrolase [EC:3.7.1.28] | 8.12 | 0.004 | 1.00 | control | K01559 | GC_00011190, GC_00013200 | 0.0 | 0.6 | Paraburkholderia |
| antitoxin ParD1/3/4 | 8.12 | 0.004 | 1.00 | control | K07746 | GC_00011027, GC_00022627 | 0.0 | 0.6 | Paraburkholderia |
| 3-oxoisoapionate kinase [EC:2.7.1.231] | 8.12 | 0.004 | 1.00 | control | K23247 | GC_00011463, GC_00013087 | 0.0 | 0.6 | Paraburkholderia |
| acrylyl-CoA reductase (NADPH) [EC:1.3.1.-] | 8.12 | 0.004 | 1.00 | control | K19745 | GC_00008334 | 0.0 | 0.6 | Paraburkholderia |
| aromatic-L-amino-acid/L-tryptophan decarboxylase [EC:4.1.1.28 4.1.1.105] | 8.12 | 0.004 | 1.00 | control | K01593 | GC_00011166, GC_00012791 | 0.0 | 0.6 | Paraburkholderia |
| LuxR family transcriptional regulator, transcriptional regulator of spore coat protein | 8.04 | 0.005 | 1.00 | control | K01994 | GC_00008552, GC_00015838 | 0.1 | 0.8 | Paraburkholderia |
| carnitine-CoA ligase [EC:6.2.1.48] | 5.66 | 0.017 | 1.00 | control | K02182 | GC_00008118, GC_00010237, GC_00012914, GC_00019298, GC_00021041, GC_00022782, GC_00024366 | 0.4 | 1.0 | Paraburkholderia |
| alpha-galactosidase [EC:3.2.1.22] | 5.60 | 0.018 | 1.00 | control | K07407 | GC_00007528, GC_00022964 | 0.2 | 0.8 | Paraburkholderia |
| toxin CcdB | 5.60 | 0.018 | 1.00 | control | K19163 | GC_00009832, GC_00012944, GC_00021954 | 0.2 | 0.8 | Paraburkholderia |
| ATP-dependent Clp protease ATP-binding subunit ClpC | 5.60 | 0.018 | 1.00 | control | K03696 | GC_00007410, GC_00008183 | 0.2 | 0.8 | Paraburkholderia |

#### Table S4. KOfam Functional Enrichment

| Description | Enrichment Score | P-value | Q-value | Associated Treatment | KO Identifier | Gene Cluster IDs | p_Heated | p_Control | Clade |
|---|---|---|---|---|---|---|---|---|---|
| cation:H+ antiporter | 5.60 | 0.018 | 1.00 | control | K07301 | GC_00006657 | 0.2 | 0.8 | Paraburkholderia |
| antitoxin CcdA | 5.60 | 0.018 | 1.00 | control | K19164 | GC_00009506, GC_00009507, GC_00018685, GC_00020449 | 0.2 | 0.8 | Paraburkholderia |
| lysozyme [EC:3.2.1.17] | 5.60 | 0.018 | 1.00 | control | K01185 | GC_00010956, GC_00012539, GC_00014544, GC_00014860, GC_00017216, GC_00019240 | 0.2 | 0.8 | Paraburkholderia |
| deoxyribodipyrimidine photo-lyase [EC:4.1.99.3] | 5.03 | 0.025 | 1.00 | heated | K01669 | GC_00003501 | 1.0 | 0.6 | Paraburkholderia |
| putative colanic acid biosynthesis glycosyltransferase WcaI | 5.03 | 0.025 | 1.00 | heated | K03208 | GC_00000853, GC_00024058 | 1.0 | 0.6 | Paraburkholderia |
| alkyldihydroxyacetonephosphate synthase [EC:2.5.1.26] | 5.03 | 0.025 | 1.00 | control | K00803 | GC_00013098 | 0.0 | 0.4 | Paraburkholderia |
| 3-oxocholest-4-en-26-oyl-CoA dehydrogenase beta subunit [EC:1.3.99.-] | 5.03 | 0.025 | 1.00 | control | K22819 | GC_00010244 | 0.0 | 0.4 | Paraburkholderia |
| membrane fusion protein, type I secretion system | 5.03 | 0.025 | 1.00 | control | K12537 | GC_00013006 | 0.0 | 0.4 | Paraburkholderia |
| MarR family transcriptional regulator, 2-MHQ and catechol-resistance regulon repressor | 5.03 | 0.025 | 1.00 | control | K15973 | GC_00011401 | 0.0 | 0.4 | Paraburkholderia |
| 4-hydroxybutyrate dehydrogenase / sulfolactaldehyde 3-reductase [EC:1.1.1.61 1.1.1.373] | 5.03 | 0.025 | 1.00 | control | K08318 | GC_00011856 | 0.0 | 0.4 | Paraburkholderia |
| 3,4-dihydroxy-9,10-secoandrosta-1,3,5(10)-triene-9,17-dione 4,5-dioxygenase [EC:1.13.11.25] | 5.03 | 0.025 | 1.00 | control | K16049 | GC_00013586, GC_00018746 | 0.0 | 0.4 | Paraburkholderia |
| D-apionate oxidoisomerase [EC:1.1.1.421] | 5.03 | 0.025 | 1.00 | control | K23245 | GC_00012444 | 0.0 | 0.4 | Paraburkholderia |
| ATP-binding cassette, subfamily C, type I secretion system permease/ATPase | 5.03 | 0.025 | 1.00 | control | K12536 | GC_00013686 | 0.0 | 0.4 | Paraburkholderia |
| N-acetylglucosaminyldiphosphoundecaprenol N-acetyl-beta-D-mannosaminyltransferase [EC:2.4.1.187] | 5.03 | 0.025 | 1.00 | control | K05946 | GC_00012630 | 0.0 | 0.4 | Paraburkholderia |
| 3-hydroxybutyryl-CoA dehydratase [EC:4.2.1.55] | 5.03 | 0.025 | 1.00 | control | K17865 | GC_00013298 | 0.0 | 0.4 | Paraburkholderia |
| 5-methylcytosine-specific restriction enzyme subunit McrC | 5.03 | 0.025 | 1.00 | control | K19147 | GC_00016739, GC_00024226 | 0.0 | 0.4 | Paraburkholderia |
| antitoxin FitA | 5.03 | 0.025 | 1.00 | control | K21495 | GC_00015636, GC_00022972 | 0.0 | 0.4 | Paraburkholderia |
| curli production protein | 5.03 | 0.025 | 1.00 | control | K04336 | GC_00013201 | 0.0 | 0.4 | Paraburkholderia |
| fumarate hydratase subunit beta [EC:4.2.1.2] | 5.03 | 0.025 | 1.00 | control | K01678 | GC_00011596 | 0.0 | 0.4 | Paraburkholderia |
| GntR family transcriptional regulator, sialic acid-inducible nan operon repressor | 5.03 | 0.025 | 1.00 | control | K22104 | GC_00011398 | 0.0 | 0.4 | Paraburkholderia |
| glycosyl transferase, family 25 | 5.03 | 0.025 | 1.00 | control | K07270 | GC_00012466 | 0.0 | 0.4 | Paraburkholderia |
| 5-oxoprolinase (ATP-hydrolysing) [EC:3.5.2.9] | 5.03 | 0.025 | 1.00 | control | K01469 | GC_00012069 | 0.0 | 0.4 | Paraburkholderia |
| fumarate hydratase subunit alpha [EC:4.2.1.2] | 5.03 | 0.025 | 1.00 | control | K01677 | GC_00012745 | 0.0 | 0.4 | Paraburkholderia |
| 2,4-dienoyl-CoA reductase [(3E)-enoyl-CoA-producing], peroxisomal [EC:1.3.1.124] | 5.03 | 0.025 | 1.00 | control | K13237 | GC_00011423, GC_00013011 | 0.0 | 0.4 | Paraburkholderia |
| GntR family transcriptional regulator, glc operon transcriptional activator | 5.03 | 0.025 | 1.00 | control | K11474 | GC_00012112 | 0.0 | 0.4 | Paraburkholderia |
| sulfopyruvate decarboxylase subunit beta [EC:4.1.1.79] | 5.03 | 0.025 | 1.00 | control | K13039 | GC_00010425 | 0.0 | 0.4 | Paraburkholderia |
| dimethylsulfone monooxygenase [EC:1.14.14.35] | 5.03 | 0.025 | 1.00 | control | K17228 | GC_00012582 | 0.0 | 0.4 | Paraburkholderia |
| general L-amino acid transport system substrate-binding protein | 5.03 | 0.025 | 1.00 | control | K09969 | GC_00009259, GC_00014341 | 0.0 | 0.4 | Paraburkholderia |
| mannosyltransferase [EC:2.4.1.-] | 5.03 | 0.025 | 1.00 | control | K14340 | GC_00013899, GC_00020125 | 0.0 | 0.4 | Paraburkholderia |
| glycerol uptake operon antiterminator | 5.03 | 0.025 | 1.00 | control | K02443 | GC_00012818 | 0.0 | 0.4 | Paraburkholderia |
| sulfite dehydrogenase (cytochrome) subunit B [EC:1.8.2.1] | 4.75 | 0.029 | 1.00 | control | K00386 | GC_00010228, GC_00021755 | 0.1 | 0.6 | Paraburkholderia |

#### Table S4. KOfam Functional Enrichment

| Description | Enrichment Score | P-value | Q-value | Associated Treatment | KO Identifier | Gene Cluster IDs | p_Heated | p_Control | Clade |
|---|---|---|---|---|---|---|---|---|---|
| LytTR family transcriptional regulator, CO-responsive transcriptional regulator RcoM | 4.75 | 0.029 | 1.00 | control | K21696 | GC_00008244 | 0.1 | 0.6 | Paraburkholderia |
| 2-oxopent-4-enoate/cis-2-oxohex-4-enoate hydratase [EC:4.2.1.80 4.2.1.132] | 4.75 | 0.029 | 1.00 | control | K18364 | GC_00008491 | 0.1 | 0.6 | Paraburkholderia |
| phosphatidylinositol-3-phosphatase [EC:3.1.3.64] | 4.75 | 0.029 | 1.00 | control | K21302 | GC_00008634, GC_00023561 | 0.1 | 0.6 | Paraburkholderia |
| zinc D-Ala-D-Ala dipeptidase [EC:3.4.13.22] | 4.75 | 0.029 | 1.00 | control | K08641 | GC_00008012 | 0.1 | 0.6 | Paraburkholderia |
| sulfite dehydrogenase (cytochrome) subunit A [EC:1.8.2.1] | 4.75 | 0.029 | 1.00 | control | K05301 | GC_00008106 | 0.1 | 0.6 | Paraburkholderia |
| salicylate 5-hydroxylase large subunit [EC:1.14.13.172] | 4.75 | 0.029 | 1.00 | control | K18242 | GC_00007681 | 0.1 | 0.6 | Paraburkholderia |
| TetR/AcrR family transcriptional regulator, mexJK operon transcriptional repressor | 4.75 | 0.029 | 1.00 | heated | K18301 | GC_00004474, GC_00007246, GC_00012732, GC_00016990, GC_00018820 | 0.9 | 0.4 | Paraburkholderia |
| acetoacetyl-CoA synthetase [EC:6.2.1.16] | 4.75 | 0.029 | 1.00 | heated | K01907 | GC_00005396, GC_00010168, GC_00018859 | 0.9 | 0.4 | Paraburkholderia |
| toxin HigB-1 | 4.75 | 0.029 | 1.00 | heated | K07334 | GC_00003962, GC_00008900, GC_00010510 | 0.9 | 0.4 | Paraburkholderia |
| toxin CptA | 4.75 | 0.029 | 1.00 | heated | K19168 | GC_00003930 | 0.9 | 0.4 | Paraburkholderia |
| IclR family transcriptional regulator, mhp operon transcriptional activator | 4.36 | 0.037 | 1.00 | control | K05818 | GC_00008314, GC_00008806, GC_00009979, GC_00011757, GC_00014364, GC_00016559, GC_00021690 | 0.5 | 1.0 | Paraburkholderia |
| HTH-type transcriptional regulator, competence development regulator | 4.36 | 0.037 | 1.00 | heated | K22299 | GC_00006163, GC_00010744 | 0.5 | 0.0 | Paraburkholderia |
| mRNA interferase YafQ [EC:3.1.-.-] | 4.36 | 0.037 | 1.00 | heated | K19157 | GC_00006181 | 0.5 | 0.0 | Paraburkholderia |
| D-galactose 1-dehydrogenase [EC:1.1.1.48] | 3.88 | 0.049 | 1.00 | heated | K00035 | GC_00004983 | 0.7 | 0.2 | Paraburkholderia |
| citronellol/citronellal dehydrogenase | 3.88 | 0.049 | 1.00 | control | K13775 | GC_00005054, GC_00007500, GC_00016332 | 0.3 | 0.8 | Paraburkholderia |
| isobutyryl-CoA mutase [EC:5.4.99.13] | 3.88 | 0.049 | 1.00 | control | K11942 | GC_00006141 | 0.3 | 0.8 | Paraburkholderia |
| protein PsiE | 3.88 | 0.049 | 1.00 | control | K13256 | GC_00007231, GC_00008902 | 0.3 | 0.8 | Paraburkholderia |
| benzoate-CoA ligase [EC:6.2.1.25] | 3.88 | 0.049 | 1.00 | control | K04110 | GC_00006098 | 0.3 | 0.8 | Paraburkholderia |
| type IV secretion system protein TrbG | 3.88 | 0.049 | 1.00 | control | K20532 | GC_00006581, GC_00014624 | 0.3 | 0.8 | Paraburkholderia |
| choline-sulfatase [EC:3.1.6.6] | 3.88 | 0.049 | 1.00 | control | K01133 | GC_00006116 | 0.3 | 0.8 | Paraburkholderia |
| XRE family transcriptional regulator, aerobic/anaerobic benzoate catabolism transcriptional regulator | 3.88 | 0.049 | 1.00 | control | K15546 | GC_00006001, GC_00017904 | 0.3 | 0.8 | Paraburkholderia |
| molybdate transport system permease protein | 3.88 | 0.049 | 1.00 | control | K02018 | GC_00005996 | 0.3 | 0.8 | Paraburkholderia |
| benzoyl-CoA 2,3-epoxidase subunit A [EC:1.14.13.208] | 3.88 | 0.049 | 1.00 | control | K15511 | GC_00006088 | 0.3 | 0.8 | Paraburkholderia |
| ribulokinase [EC:2.7.1.16] | 3.88 | 0.049 | 1.00 | control | K24707 | GC_00006147 | 0.3 | 0.8 | Paraburkholderia |
| benzoyl-CoA-dihydrodiol lyase [EC:4.1.2.44] | 3.88 | 0.049 | 1.00 | control | K15513 | GC_00006136 | 0.3 | 0.8 | Paraburkholderia |
| type IV secretion system protein TrbF | 3.88 | 0.049 | 1.00 | control | K20531 | GC_00006552, GC_00017401 | 0.3 | 0.8 | Paraburkholderia |
| entericidin A | 3.88 | 0.049 | 1.00 | control | K16347 | GC_00005951 | 0.3 | 0.8 | Paraburkholderia |
| L-lactate dehydrogenase (cytochrome) [EC:1.1.2.3] | 3.88 | 0.049 | 1.00 | control | K00101 | GC_00005524 | 0.3 | 0.8 | Paraburkholderia |
| benzoyl-CoA 2,3-epoxidase subunit B [EC:1.14.13.208] | 3.88 | 0.049 | 1.00 | control | K15512 | GC_00006056 | 0.3 | 0.8 | Paraburkholderia |
| (S)-mandelate dehydrogenase [EC:1.1.99.31] | 3.88 | 0.049 | 1.00 | control | K15054 | GC_00006560, GC_00009472, GC_00017903 | 0.3 | 0.8 | Paraburkholderia |

#### Table S5. KEGG Module Functional Enrichment

| Description | Enrichment Score | P-value | Q-value | Associated Treatment | KEGG Identifier | Gene Cluster IDs | p_Heated | p_Control | Clade |
|---|---|---|---|---|---|---|---|---|---|
| Methanofuran biosynthesis | 8.12 | 0.004 | 0.86 | control | M00935 | GC_00002980, GC_00017119 | 0.3 | 1.0 | Kitasatospora |
| Nicotinate degradation, nicotinate => fumarate | 7.02 | 0.008 | 0.86 | control | M00622 | GC_00005486, GC_00028536, GC_00028579 | 0.3 | 1.0 | Kitasatospora |
| Calicheamicin biosynthesis, calicheamicinone => calicheamicin | 7.02 | 0.008 | 0.86 | heated | M00833 | GC_00005614, GC_00005620, GC_00009704, GC_00016311, GC_00026255 | 0.7 | 0.0 | Kitasatospora |
| Malonate semialdehyde pathway, propanoyl-CoA => acetyl-CoA;beta-Oxidation;beta-Oxidation, peroxisome, VLCFA;Jasmonic acid biosynthesis | 6.39 | 0.011 | 0.87 | control | M00013;M00087; M00861;M00113 | GC_00007179, GC_00009968, GC_00013461, GC_00023709 | 0.2 | 0.8 | Kitasatospora |
| Cysteine biosynthesis, serine => cysteine | 6.09 | 0.014 | 0.87 | heated | M00021 | GC_00004630, GC_00025323, GC_00029039, GC_00029649 | 0.6 | 0.0 | Kitasatospora |
| Fosfomycin biosynthesis, phosphoenolpyruvate => fosfomycin | 5.38 | 0.020 | 0.91 | control | M00903 | GC_00010304, GC_00021033 | 0.1 | 0.6 | Kitasatospora |
| Heparan sulfate degradation | 5.20 | 0.023 | 0.91 | control | M00078 | GC_00005322 | 0.3 | 0.8 | Kitasatospora |
| Homoprotocatechuate degradation, homoprotocatechuate => 2-oxohept-3-enedioate | 5.20 | 0.023 | 0.91 | heated | M00533 | GC_00004867, GC_00015586, GC_00016351, GC_00018400, GC_00019132 | 0.7 | 0.2 | Kitasatospora |
| Dihydrokalafungin biosynthesis, octaketide => dihydrokalafungin | 4.77 | 0.029 | 1.00 | heated | M00779 | GC_00004048, GC_00006202, GC_00006714, GC_00011263, GC_00011682, GC_00012840, GC_00016375 | 1.0 | 0.8 | Kitasatospora |
| Dissimilatory nitrate reduction, nitrate => ammonia | 4.57 | 0.033 | 1.00 | heated | M00530 | GC_00004972, GC_00005380, GC_00020999 | 0.5 | 0.0 | Kitasatospora |
| Betacyanin biosynthesis, L-tyrosine => amaranthin | 4.23 | 0.040 | 1.00 | control | M00961 | GC_00010082, GC_00011523 | 0.3 | 0.8 | Kitasatospora |
| Tryptophan metabolism, tryptophan => kynurenine => 2-aminomuconate | 3.98 | 0.046 | 1.00 | heated | M00038 | GC_00004626, GC_00008818 | 0.8 | 0.4 | Kitasatospora |
| Coenzyme M biosynthesis | 3.94 | 0.047 | 1.00 | heated | M00358 | GC_00005869, GC_00008057 | 0.5 | 0.0 | Kitasatospora |
| Assimilatory nitrate reduction, nitrate => ammonia;Nitrate assimilation | 3.94 | 0.047 | 1.00 | heated | M00531;M00615 | GC_00005201, GC_00005371 | 0.5 | 0.0 | Kitasatospora |
| Ubiquinone biosynthesis, prokaryotes, chorismate (+ polyprenyl-PP) => ubiquinol;C5 isoprenoid biosynthesis, mevalonate pathway, archaea | 5.00 | 0.025 | 1.00 | heated | M00117;M00849 | GC_00006040, GC_00009484, GC_00010750 | 0.8 | 0.2 | Bradyrhizobium |
| Pimeloyl-ACP biosynthesis, BioC-BioH pathway, malonyl-ACP => pimeloyl-ACP | 4.62 | 0.032 | 1.00 | control | M00572 | GC_00003531 | 0.6 | 1.0 | Bradyrhizobium |
| Methionine salvage pathway;Polyamine biosynthesis, arginine => agmatine => putrescine => spermidine | 4.62 | 0.032 | 1.00 | control | M00034;M00133 | GC_00004761, GC_00004893, GC_00015455, GC_00020520 | 0.6 | 1.0 | Bradyrhizobium |
| GABA (gamma-Aminobutyrate) shunt | 4.26 | 0.039 | 1.00 | heated | M00027 | GC_00010877, GC_00012424, GC_00025959 | 0.6 | 0.1 | Bradyrhizobium |
| Phthalate degradation, phthalate => protocatechuate | 7.78 | 0.005 | 0.77 | heated | M00623 | GC_00003733 | 1.0 | 0.2 | Rhizobium |
| Pantothenate biosynthesis, valine/L-aspartate => pantothenate | 5.83 | 0.016 | 0.77 | heated | M00119 | GC_00004562 | 1.0 | 0.3 | Rhizobium |

#### Table S5. KEGG Module Functional Enrichment

| Description | Enrichment Score | P-value | Q-value | Associated Treatment | KEGG Identifier | Gene Cluster IDs | p_Heated | p_Control | Clade |
|---|---|---|---|---|---|---|---|---|---|
| C-1027 beta-amino acid moiety biosynthesis, tyrosine => 3-chloro-4,5-dihydroxy-beta-phenylalanyl-PCP;Maduropeptin beta-hydroxy acid moiety biosynthesis, tyrosine => 3-(4-hydroxyphenyl)-3-oxopropanoyl-PCP;Kedarcidin 2-aza-3-chloro-beta-tyrosine moiety biosynthesis, azatyrosine => 2-aza-3-chloro-beta-tyrosyl-PCP | 5.83 | 0.016 | 0.77 | heated | M00827;M00828; M00832 | GC_00004498 | 1.0 | 0.3 | Rhizobium |
| Nocardicin A biosynthesis, L-pHPG + arginine + serine => nocardicin A | 5.83 | 0.016 | 0.77 | heated | M00736 | GC_00004590 | 1.0 | 0.3 | Rhizobium |
| ADP-L-glycero-D-manno-heptose biosynthesis | 5.83 | 0.016 | 0.77 | control | M00064 | GC_00005369, GC_00005374, GC_00005376, GC_00006750, GC_00006849 | 0.0 | 0.7 | Rhizobium |
| Ubiquinone biosynthesis, prokaryotes, chorismate (+ polyprenyl-PP) => ubiquinol;Ubiquinone biosynthesis, eukaryotes, 4-hydroxybenzoate + polyprenyl-PP => ubiquinol | 4.32 | 0.038 | 0.90 | control | M00117;M00128 | GC_00005231 | 0.0 | 0.6 | Rhizobium |
| Tryptophan metabolism, tryptophan => kynurenine => 2-aminomuconate;NAD biosynthesis, tryptophan => quinolinate => NAD | 4.32 | 0.038 | 0.90 | control | M00038;M00912 | GC_00005162, GC_00005447, GC_00005773, GC_00006745 | 0.0 | 0.6 | Rhizobium |
| GABA biosynthesis, eukaryotes, putrescine => GABA;Pantothenate biosynthesis, 2-oxoisovalerate/spermine => pantothenate | 4.32 | 0.038 | 0.90 | control | M00135;M00913 | GC_00004655 | 0.0 | 0.6 | Rhizobium |
| Ectoine degradation, ectoine => aspartate | 4.32 | 0.038 | 0.90 | control | M00919 | GC_00005082, GC_00005157, GC_00005174, GC_00005195 | 0.0 | 0.6 | Rhizobium |
| Helicobacter pylori pathogenicity signature, cagA pathogenicity island | 4.32 | 0.038 | 0.90 | control | M00564 | GC_00005046 | 0.0 | 0.6 | Rhizobium |
| Phenylalanine biosynthesis, chorismate => phenylpyruvate => phenylalanine;Tyrosine biosynthesis, chorismate => HPP => tyrosine;Tyrosine biosynthesis, chorismate => arogenate => tyrosine | 4.20 | 0.040 | 0.90 | control | M00024;M00025; M00040 | GC_00004002, GC_00005864 | 0.6 | 1.0 | Rhizobium |
| Catecholamine biosynthesis, tyrosine => dopamine => noradrenaline => adrenaline;Melatonin biosynthesis, animals, tryptophan => serotonin => melatonin;Melatonin biosynthesis, plants, tryptophan => serotonin => melatonin | 8.12 | 0.004 | 0.73 | control | M00042;M00037; M00936 | GC_00011166, GC_00012791 | 0.0 | 0.6 | Paraburkholderia |
| Vancomycin resistance, D-Ala-D-Lac type | 8.04 | 0.005 | 0.73 | control | M00651 | GC_00008012, GC_00017324, GC_00022669 | 0.1 | 0.8 | Paraburkholderia |
| Citrate cycle (TCA cycle, Krebs cycle);Citrate cycle, second carbon oxidation, 2-oxoglutarate => oxaloacetate;Reductive citrate cycle (Arnon-Buchanan cycle);Dicarboxylate-hydroxybutyrate cycle;Incomplete reductive citrate cycle, acetyl-CoA => oxoglutarate;Anoxygenic photosynthesis in green nonsulfur bacteria;Anoxygenic photosynthesis in green sulfur bacteria | 5.03 | 0.025 | 1.00 | control | M00009;M00011; M00173; M00374; M00620; M00613; M00614 | GC_00011596, GC_00012745 | 0.0 | 0.4 | Paraburkholderia |
| Ethylmalonyl pathway | 5.03 | 0.025 | 1.00 | control | M00373 | GC_00013298 | 0.0 | 0.4 | Paraburkholderia |
| Salicylate degradation, salicylate => gentisate | 4.75 | 0.029 | 1.00 | control | M00638 | GC_00007681, GC_00008591 | 0.1 | 0.6 | Paraburkholderia |
| Anoxygenic photosynthesis in green nonsulfur bacteria | 3.88 | 0.049 | 1.00 | control | M00613 | GC_00006141 | 0.3 | 0.8 | Paraburkholderia |

#### Table S6. Metabolic Module Functional Enrichment

| Description | Enrichment Score | P-value | Q-value | Associated Treatment | Module Identifier | Genomes | p_Heated | p_Control | Clade |
|---|---|---|---|---|---|---|---|---|---|
| Citrate cycle, first carbon oxidation, oxaloacetate => 2-oxoglutarate | 9.91 | 0.002 | 0.10 | heated | M00010 | GP157, GP160, GP163, GP28, GP31, GP36, GP50, GP55, MAA19, MAA2, MAA52, MAA75, MAA81, MAP12-4, MAP12-9, MAP2-59, MAP5-34, MAP5-40, MAP8-42, GAS204B, MAP12-44, MAP12-15, MAA4, MAA18, GP30, GP82 | 1.0 | 0.6 | Kitasatospora |
| Reductive citrate cycle (Arnon-Buchanan cycle) | 9.91 | 0.002 | 0.10 | heated | M00173 | GP157, GP160, GP163, GP28, GP31, GP36, GP50, GP55, MAA19, MAA2, MAA52, MAA75, MAA81, MAP12-4, MAP12-9, MAP2-59, MAP5-34, MAP5-40, MAP8-42, GAS204B, MAP12-44, MAP12-15, MAA4, MAA18, GP30, GP82 | 1.0 | 0.6 | Kitasatospora |
| Trans-cinnamate degradation, trans-cinnamate => acetyl-CoA | 5.38 | 0.020 | 0.49 | control | M00545 | GAS1054, GP163, GP50, MAA52, MAA75, GAS204B | 0.1 | 0.6 | Kitasatospora |
| GABA biosynthesis, eukaryotes, putrescine => GABA | 4.77 | 0.029 | 0.49 | heated | M00135 | GAS1054, GP157, GP160, GP163, GP28, GP31, GP36, GP50, GP55, MAA19, MAA2, MAA52, MAA75, MAA81, MAP12-4, MAP12-9, MAP25-9, MAP5-34, MAP5-40, MAP8-42, GAS204B, MAP124-4, MAP12-15, MAA4, MAA18, GP30, GP82 | 1.0 | 0.8 | Kitasatospora |
| Glyoxylate cycle | 4.77 | 0.029 | 0.49 | heated | M00012 | GP157, GP160, GP163, GP28, GP31, GP36, GP50, GP55, MAA19, MAA2, MAA36, MAA52, MAA75, MAA81, MAP12-4, MAP129, MAP2-59, MAP5-34, MAP5-40, MAP8-42, GAS204B, MAP12-44, MAP12-15, MAA4, MAA18, GP30, GP82 | 1.0 | 0.8 | Kitasatospora |
| Pantothenate biosynthesis, 2-oxoisovalerate/spermine => pantothenate | 4.77 | 0.029 | 0.49 | heated | M00913 | GAS1054, GP157, GP160, GP163, GP28, GP31, GP36, GP50, GP55, MAA19, MAA2, MAA52, MAA75, MAA81, MAP12-4, MAP12-9, MAP2-59, MAP5-34, MAP5-40, MAP8-42, GAS204B, MAP12-44, MAP12-15, MAA4, MAA18, GP30, GP82 | 1.0 | 0.8 | Kitasatospora |
| beta-Oxidation | 4.77 | 0.029 | 0.49 | heated | M00087 | GAS1054, GP157, GP160, GP163, GP28, GP31, GP36, GP50, GP55, MAA19, MAA2, MAA52, MAA75, MAA81, MAP12-4, MAP12-9, MAP2-59, MAP5-34, MAP5-40, MAP8-42, GAS204B, MAP12-44, MAP12-15, MAA4, MAA18, GP30, GP82 | 1.0 | 0.8 | Kitasatospora |
| beta-Lactam resistance, Bla system | 4.62 | 0.032 | 1.00 | heated | M00627 | GAS242, MAP5-43 | 0.4 | 0.0 | Bradyrhizobium |
| Purine degradation, xanthine => urea | 7.78 | 0.005 | 0.53 | control | M00546 | AN5, AN64, AN68, AN70, AN72, AN73, AN88 | 0.0 | 0.8 | Rhizobium |
| Assimilatory nitrate reduction, nitrate => ammonia | 6.64 | 0.010 | 0.53 | control | M00531 | AN5, AN64, AN68, AN69, AN6A, AN70, AN72, AN73, AN88 | 0.2 | 0.9 | Rhizobium |
| Ectoine degradation, ectoine => aspartate | 4.32 | 0.038 | 0.61 | control | M00919 | AN5, AN64, AN68, AN72, AN88 | 0.0 | 0.6 | Rhizobium |

#### Table S6. Metabolic Module Functional Enrichment

| Description | Enrichment Score | P-value | Q-value | Associated Treatment | Module Identifier | Genomes | p_Heated | p_Control | Clade |
|---|---|---|---|---|---|---|---|---|---|
| Methionine degradation | 4.32 | 0.038 | 0.61 | heated | M00035 | AN5, AN63, AN67, AN69, AN6A, AN72, AN83, AN95, 28DA2 | 1.0 | 0.4 | Rhizobium |
| Pantothenate biosynthesis, 2-oxoisovalerate/spermine => pantothenate | 4.32 | 0.038 | 0.61 | control | M00913 | AN64, AN68, AN70, AN73, AN88 | 0.0 | 0.6 | Rhizobium |
| Cytochrome c oxidase, prokaryotes | 4.20 | 0.040 | 0.61 | control | M00155 | AN5, AN63, AN64, AN67, AN68, AN69, AN6A, AN70, AN72, AN73, AN83, AN88 | 0.6 | 1.0 | Rhizobium |
| Multidrug resistance, efflux pump MexJK-OprM | 4.20 | 0.040 | 0.61 | heated | M00642 | AN63, AN83 | 0.4 | 0.0 | Rhizobium |

#### Table S7. CAZyme Functional Enrichment

| CAZyme HMM | Enzyme Class | Family Activity Description | Substrate | Enrichment Score | P-value | Q-value | Associated Treatment | Gene Cluster IDs | p_Heated | p_Control | Clade |
|---|---|---|---|---|---|---|---|---|---|---|---|
| GH39.hmm | Glycoside hydrolases | 3-O-β-L-arabinopyranosyl-α-L-arabinofuranosidase (EC 3.2.1.-); Endo-α-L-rhamnosidase (EC 3.2.1.-); β-glucosidase (EC 3.2.1.21); 3-O-α-D-galactosyl-α-L-arabinofuranosidase (EC 3.2.1.215); β-galactosidase (EC 3.2.1.23); Xylan β-1,4-xylosidase (EC 3.2.1.37); Exo-β-1,4-glucanase / cellodextrinase (EC 3.2.1.74); α-L-iduronidase (EC 3.2.1.76); | xylan, cellobiose, cellulose | 6.39 | 0.011 | 0.73 | control | GC_00007371, GC_00017393 | 0.2 | 0.8 | Kitasatospora |
| GH19.hmm | Glycoside hydrolases | Chitinase (EC 3.2.1.14); Lysozyme (EC 3.2.1.17); [reducing end] exo-chitinase (EC 3.2.1.201); | chitin | 5.28 | 0.022 | 0.73 | control | GC_00003830 | 0.4 | 1.0 | Kitasatospora |
| GH89.hmm | Glycoside hydrolases | α-N-acetylglucosaminidase (EC 3.2.1.50); | | 5.20 | 0.023 | 0.73 | control | GC_00005322 | 0.3 | 0.8 | Kitasatospora |
| GH13_21.hmm | Glycoside hydrolases | [retaining] α-amylase (EC 3.2.1.1); α-glucosidase (EC 3.2.1.20); | starch, glycogen | 4.77 | 0.029 | 0.73 | control | GC_00023147 | 0.0 | 0.2 | Kitasatospora |
| PL31.hmm | Polysaccharide Lyases | Endo-β-1,4-glucuronan lyase (EC 4.2.2.14); Poly(β-mannuronate) lyase / M-specific alginate lyase (EC 4.2.2.3); | alginate | 4.77 | 0.029 | 0.73 | heated | GC_00000510, GC_00006367, GC_00019336 | 1.0 | 0.8 | Kitasatospora |
| GH13_3.hmm | Glycoside hydrolases | α-1,4-glucan: phosphate α-maltosyltransferase (EC 2.4.99.16); | | 4.77 | 0.029 | 0.73 | heated | GC_00000219 | 1.0 | 0.8 | Kitasatospora |
| GH114.hmm | Glycoside hydrolases | [retaining] endo-α-1,4-galactosaminidase (EC 3.2.1.109); | | 4.57 | 0.033 | 0.73 | control | GC_00004350, GC_00020399 | 0.5 | 1.0 | Kitasatospora |
| GH119.hmm | Glycoside hydrolases | [retaining] α-amylase (EC 3.2.1.1); | starch | 4.23 | 0.040 | 0.78 | control | GC_00005030, GC_00013232 | 0.3 | 0.8 | Kitasatospora |
| GH33.hmm | Glycoside hydrolases | Trans-sialidase (EC 2.4.1.-); 2-keto-3-deoxynononic acid hydrolase / KDNase (EC 3.2.1.-); Kdo hydrolase (EC 3.2.1.124); Exo-α-sialidase (EC 3.2.1.18); Anhydrosialidase (EC 4.2.2.15); | | 3.94 | 0.047 | 0.82 | heated | GC_00005197, GC_00009621 | 0.5 | 0.0 | Kitasatospora |
| GH74.hmm | Glycoside hydrolases | Oligoxyloglucan reducing-end-specific cellobiohydrolase (EC 3.2.1.150); Xyloglucan-specific endo-β-1,4-glucanase / endo-xyloglucanase (EC 3.2.1.151); Endo-β-1,4-glucanase (EC 3.2.1.4); | xyloglucan, cellulose | 4.26 | 0.039 | 1.00 | heated | GC_00010635, GC_00011244, GC_00017774, GC_00018033, GC_00018895, GC_00021200, GC_00025217 | 0.6 | 0.1 | Bradyrhizobium |
| CE19.hmm | Carbhodrate Esterases | Pectin methylesterase (EC 3.1.1.11); | | 4.32 | 0.038 | 0.86 | control | GC_00005263 | 0.6 | 0.0 | Rhizobium |

#### Table S7. CAZyme Functional Enrichment

| CAZyme HMM | Enzyme Class | Family Activity Description | Substrate | Enrichment Score | P-value | Q-value | Associated Treatment | Gene Cluster IDs | p_Heated | p_Control | Clade |
|---|---|---|---|---|---|---|---|---|---|---|---|
| GT2_Glyco_ tranf_2_3.hmm | Glycosyl-transferases | cellulose synthase (EC 2.4.1.12); chitin synthase (EC 2.4.1.16); dolichyl-phosphate beta-D-mannosyltransferase (EC 2.4.1.83); dolichyl-phosphate beta-glucosyltransferase (EC 2.4.1.117); N-acetylglucosaminyltransferase (EC 2.4.1.-); N-acetylgalactosaminyltransferase (EC 2.4.1.-); hyaluronan synthase (EC 2.4.1.212); chitin oligosaccharide synthase (EC 2.4.1.-); beta-1,3-glucan synthase (EC 2.4.1.34); beta-1,4-mannan synthase (EC 2.4.1.-); beta-mannosylphosphodecaprenol-mannooligosaccharide alpha-1,6-mannosyltransferase (EC 2.4.1.199); UDP-Galf: rhamnopyranosyl-N-acetylglucosaminyl-PP-decaprenol beta-1,4/1,5-galactofuranosyltransferase (EC 2.4.1.287); UDP-Galf: galactofuranosyl-galactofuranosyl-rhamnosyl-N-acetylglucosaminyl-PP-decaprenol beta-1,5/1,6-galactofuranosyltransferase (EC 2.4.1.288); dTDP-L-Rha: N-acetylglucosaminyl-PP-decaprenol alpha-1,3-L-rhamnosyltransferase (EC 2.4.1.289); alternating beta-1,3/4-N-acetylmannan synthase (2.4.1.-); UDP-GlcA: N-acetylglucosaminyl-proteoglycan beta-1,4-glucuronosyltransferase (EC 2.4.1.225); [inverting] UDP-Glc: glycocin S-beta-glucosyltransferase (EC 2.4.1.-); [inverting] UDP-Glc: protein O-beta-glucosyltransferase (EC 2.4.1.-) | | 4.32 | 0.038 | 0.86 | heated | GC_00003953 | 0.4 | 1.0 | Rhizobium |
| PL5.hmm | Polysaccharide Lyases | Endo-β-1,4-glucuronan lyase (EC 4.2.2.14); Poly(β-mannuronate) lyase / M-specific alginate lyase (EC 4.2.2.3); | alginate | 4.32 | 0.038 | 0.86 | heated | GC_00004263, GC_00004445 | 0.4 | 1.0 | Rhizobium |
| GH24.hmm | Glycoside hydrolases | Lysozyme (EC 3.2.1.17); | | 5.60 | 0.018 | 0.74 | control | GC_00010956, GC_00012539, GC_00014544, GC_00014860, GC_00017216, GC_00017413, GC_00019240 | 0.2 | 0.8 | Paraburkholderia |
| GT25.hmm | Glycosyl-transferases | LPS β-1,4-N-acetylgalactosaminyltransferase (EC 2.4.1.-); UDP-Gal: β-1,4-galactosyltransferase (EC 2.4.1.-); UDP-Glc: hydroxylysine O-glucosyltransferase (EC 2.4.1.-); β-1,2-galactosyltransferase (EC 2.4.1.-); Occidiofungin β-xylosyltransferase (EC 2.4.2.-); | | 5.03 | 0.025 | 0.74 | control | GC_00012466 | 0.0 | 0.4 | Paraburkholderia |
| AA5.hmm | Auxiliary Activities | Raffinose oxidase (EC 1.1.3.-); Alcohol oxidase (EC 1.1.3.13); 5-(hydroxymethyl)furfural oxidase (EC 1.1.3.47); Aryl alcohol oxidase (EC 1.1.3.7); Galactose oxidase (EC 1.1.3.9); Glyoxal oxidase (EC 1.2.3.15); | raffinose, galactose | 5.03 | 0.025 | 0.74 | control | GC_00012899, GC_00020189, GC_00020567, GC_00024242 | 0.0 | 0.4 | Paraburkholderia |
| GT26.hmm | Glycosyl-transferases | UDP-Gal: β-1,4-galactosyltransferase (EC 2.4.1.-); UDP-Glc: β-1,4-glucosyltransferase (EC 2.4.1.-); UDP-ManNAc: β-N-acetyl-mannosaminyltransferase (EC 2.4.1.-); UDP-ManNAcA: β-N-acetyl mannosaminuronyltransferase (EC 2.4.1.-); | | 5.03 | 0.025 | 0.74 | control | GC_00012630 | 0.0 | 0.4 | Paraburkholderia |
| GH92.hmm | Glycoside hydrolases | α-1,4-mannosidase (EC 3.2.1.-); Mannosyl-1-phosphodiester α-1,P-mannosidase (EC 3.2.1.-); Mannosyl-oligosaccharide α-1,3-mannosidase (EC 3.2.1.-); Mannosyl-oligosaccharide α-1,2-mannosidase (EC 3.2.1.113); α-mannosidase (EC 3.2.1.24); | | 3.88 | 0.049 | 0.74 | heated | GC_00004487, GC_00024113 | 0.7 | 0.2 | Paraburkholderia |
